# Orthogonal Shared Basis Factorization: Cross-species gene expression analysis using a common expression subspace

**DOI:** 10.1101/2022.08.26.505467

**Authors:** Amal Thomas

## Abstract

One of the main challenges in analyzing gene expression profiles across species is the dependence on determining corresponding genes between species. Homology-based approaches fail to account for the contribution of non-homologous genes to the phenotype, genes’ functional divergence, and rewiring of pathways. Homology-independent methods based on joint matrix factorization provide a potential solution, but biological interpretations with existing approaches are difficult. We developed a novel joint matrix factorization method that we call the orthogonal shared basis factorization (OSBF) to compare functionally similar phenotypes across species. OSBF utilizes a similar correlation structure within individual datasets to estimate interpretable matrix factors. This homology-independent approach places cellular phenotypes in a common coordinate system that can summarize gene expression patterns shared by different organisms and quantifies the role of all genes in the phenotype independent of their homology relationships and annotation. OSBF is available on GitHub.

## 1 Introduction

Comparative analysis of gene expression profiles across species has important applications in the fields of biology and medicine. Cross-species gene expression studies have addressed many biomedical research questions, including characterizing cellular phenotype, disease states, and aging (Ginis et al., 2004; Sweet-Cordero et al., 2005; De Magalhães et al., 2009; Mueller et al., 2017). Comparing gene expression data of different tissues from multiple species have identified conserved transcriptional programs driving the development and evolution of organs (Brawand et al., 2011; Merkin et al., 2012; Cardoso-Moreira et al., 2019). Conserved co-expression modules across organisms and similarity of expression patterns in developmental stages were discovered across animal phyla (Gerstein et al., 2014).

Genes participating in the same pathway or part of the same protein complex are often co-regulated and exhibit correlated expression profiles (Kim et al., 2001; Segal et al., 2003). However, similar expression patterns do not necessarily imply that genes are functionally related. For instance, apparent co-expression of genes can occur by chance as a result of transcriptional regulation. As a result, in studies limited to single species, it would be difficult to distinguish functionally important genes from those accidentally regulated (Stuart et al., 2003). Conservation of co-expression patterns in multiple species is a powerful criterion to distinguish functionally important genes from a set of co-regulated genes. Several studies have used gene co-expression information across different species to infer gene functions (Bergmann et al., 2004; Lefebvre et al., 2005; van Dam et al., 2015; Obayashi et al., 2019). In studies co-analyzing gene expression profiles across species, the usual analysis strategy starts with selecting a set of homologous genes shared by the species. The assumption here is that the shared genes, usually restricted to one-to-one orthologs, perform similar roles in different species. The relationships between the transcriptomic profiles of multiple species and the genes contributing to phenotypes are then investigated.

The relationship between gene expression and phenotype is rarely simple, and predicting phenotypic relationships based on orthologs is not always straightforward. Although many cellular processes are conserved across species, some or even all components of a pathway can be replaced by genes with independent evolutionary origin (Koonin et al., 1996). Studies on enzymatic evolution have identified structurally distinct non-homologous enzymes catalyzing the same biochemical reaction (Omelchenko et al., 2010; Galperin & Koonin, 2012). Many of the genes performing essential functions in bacterial groups do not have a shared evolutionary origin (Soucy et al., 2015). Functional divergence of human-mouse orthologs has been observed, including genes involved in DNA repair, inflammation pathways, and development (Hirano et al., 2007; Yashiro et al., 2000; Liu et al., 2010). There is evidence for homologous genes and regulatory networks adapted for a different function in both plants and animals (Kajala et al., 2021; Carelli et al., 2018). Species or lineage-specific genes potentially coding for novel proteins may also arise by partial gene duplication, shuffling of domains between genes, or completely de novo from non-coding regions (Zhang et al., 2004; Toll-Riera et al., 2009; Tautz & Domazet-Lošo, 2011; Chen et al., 2010). Collectively, the re-purposing of genes, proteins, and regulatory networks implies that independent strategies exist in nature to achieve the same molecular function, and orthology does not guarantee equivalent functions. Hence, cross-species comparative approaches focussing just on orthologs fail to identify a complete set of functionally relevant genes related to a phenotype.

A different category of methods for co-analyzing gene expression profiles independent of orthology mapping between species is based on joint matrix factorization. Using generalized singular value decomposition (GSVD) (Van Loan, 1976), Alter et al. (2003) compared cell cycle expression profiles of human and yeast. This idea was further extended to more than two species to develop higher-order GSVD (HO GSVD) (Ponnapalli et al., 2011). These methods decompose expression profiles of functionally similar phenotypes to estimate species-specific factors and a shared factor: a common expression subspace. They showed that orthology-independent cross-species analysis can be performed using this common subspace. Using cell-cycle gene expression datasets, these approaches have shown examples of genes with highly conserved sequences across species but with significantly different cell-cycle peak times. Although these methods offer a potential solution for orthology-independent analysis, the estimated common expression subspace is not designed to represent biological processes conserved across species. Most of the dimensions of the expression subspace represent biological processes specific for individual species. The species-specific factors and the shared subspace in HO GSVD allow redundancy, making it challenging to differentiate the contribution of genes to different phenotypes. As a result, identifying genes relevant for different phenotypes is not possible using these approaches.

In this work, we developed a novel joint matrix factorization called orthogonal shared basis factorization (OSBF) to estimate an expression subspace common to all species that represents the conserved correlation relationships between the phenotypes. The method factorizes the gene expression profiles of functionally similar phenotypes from multiple species into a species-specific loading factor, a species-specific diagonal factor, and a common expression subspace. For each species, the loadings for independent linear combination of genes is stored in different columns of the loading factor, and the amount of correlation information represented by different dimensions of the common expression subspace is stored in the diagonal factor. Finally, these factors are estimated such that the reconstructed expression profiles using these factors are close to the original gene expression profiles. The species-specific loading factor quantifies the contribution of all genes to different phenotypes associated with the gene expression profiles, and allows us to identify functionally relevant genes.

Using OSBF, cross-species gene expression analysis of six tissue types from eight vertebrate species shows that more than 40% of genes important for tissue phenotype are not one-to-one orthologs shared in all species. Furthermore, 6-10% of these genes do not have orthologs in other species. Several examples of gene expression divergence of orthologs that can be identified by comparing homologs are also detected in our analysis. Without relying on homology and annotation classification, our approach identifies functionally relevant genes for tissue phenotype, including known gene markers, non-coding genes, pseudogenes, and other less-studied genes.

## 2 Results

In this section, we first provide the details of our joint matrix factorization, how the different factors are estimated, and their interpretation. Then we demonstrate how cross-species gene expression comparisons are performed independent of the homology mapping between genes. Finally, we show the application of our method by analyzing public RNA-Seq profiles of different vertebrate tissue types.

### 2.1 Orthogonal Shared Basis Factorization

Consider a set of *k* real matrices 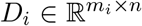, each with full column rank. Each *D*_*i*_ represents the expression of *n* tissues (columns) in species *i* with *m*_*i*_ genes (rows) annotated for that species. The *p*-th column of each *D*_*i*_ matrix corresponds to the gene expression profile of a functionally similar tissue representing a similar phenotype across species. The number of rows is different for each *D*_*i*_ matrix, but they all have the same number of columns. We define orthogonal shared basis factorization (OSBF) as

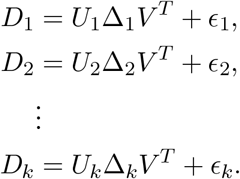

Each 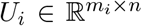 is a species-specific loading matrix with orthonormal columns (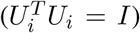) and Δ_*i*_ ∈ ℝ^*n*×*n*^ is a diagonal matrix with positive values.. The right basis matrix *V* ∈ ℝ^*n*×*n*^ is an orthogonal matrix and identical in all the *k* matrix factorizations (Additional file 1: Figure S1). In OSBF, we define the common space (*V*) as the subspace of row space (ℝ^*n*^) shared by all *k* matrices representing correlation relationships between the columns. An orthonormal basis of this shared subspace is used as the axes of the common space. We estimate the components of the matrix factorization by minimizing the total reconstruction error given by 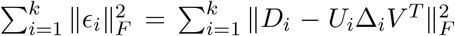. We designed an alternating least square algorithm to minimize the total factorization error (see Methods). Our algorithm determines the optimal learning rate in each update step and guarantees convergence to a stable function value (Additional file 2). A schematic summarizing the joint factorization of gene expression matrices from different species using OSBF is shown in Figure 1.

**Figure 1:**
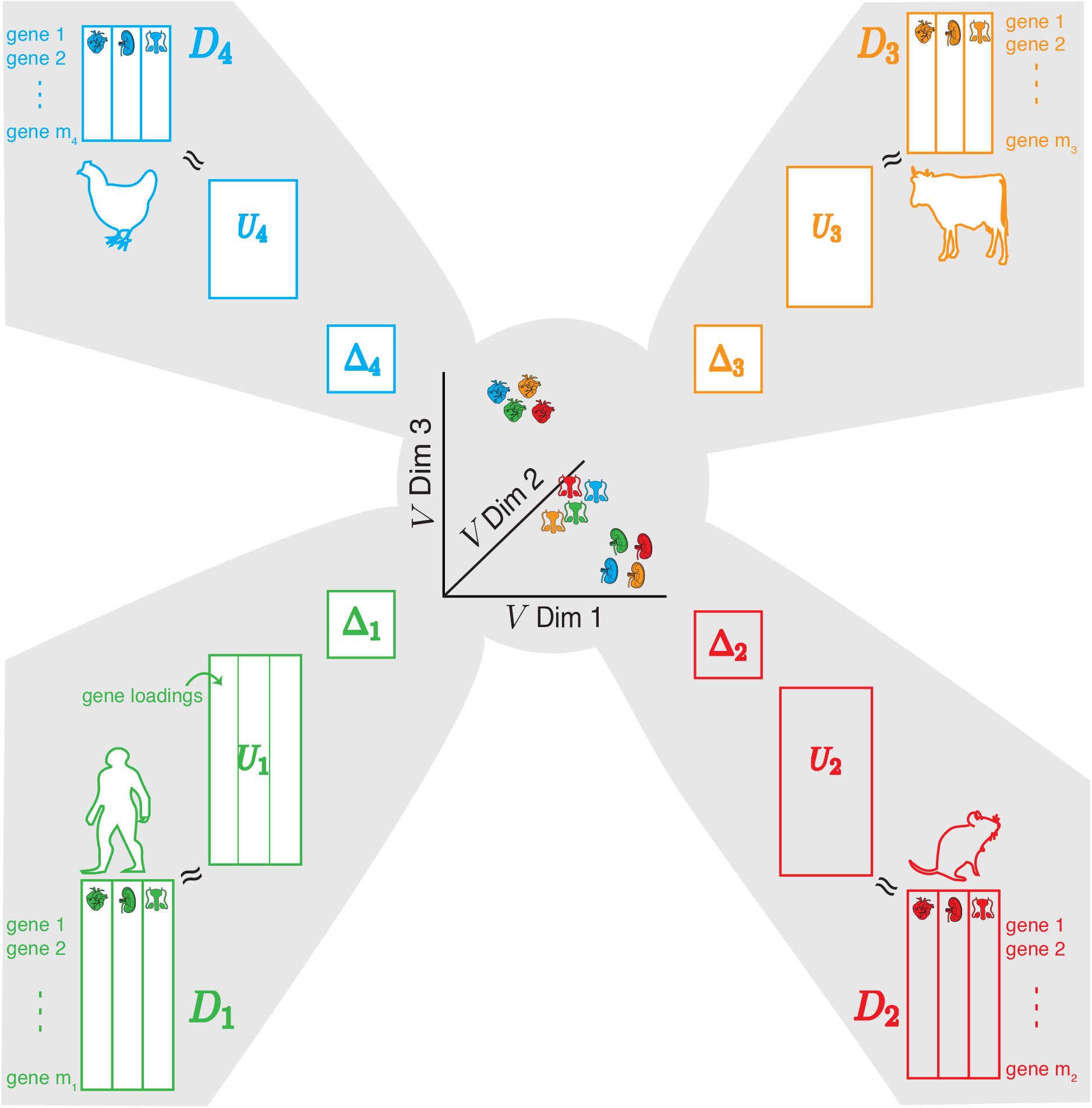
Graphical summary of OSBF: OSBF is a joint matrix factorization of *k* 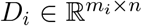 matrices. Each *D*_*i*_ matrix is decomposed into a species-specific loading matrix 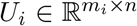, a species-specific diagonal matrix Δ_*i*_ ∈ ℝ^*n*×*n*^, and a shared orthogonal right basis matrix *V*. The columns of *U*_*i*_ (loading vectors) are orthonormal, and Δ_*i*_ has positive entries. The cartoon shows the joint factorization of four (*k* = 4) *D*_*I*_ matrices each with *n* = 3 columns. The columns of the *V* matrix represent the different dimensions of the common subspace shared by all species.

### 2.2 Estimating species-independent common expression subspace

In OSBF, the initial estimate of the common expression subspace *V* is computed based on the inter-tissue correlations within each species. Let 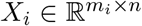 be the standardized gene expression matrix for species *I* (see Methods). Within species *i*, the correlation between expression profiles for the *n* tissues is

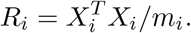

Each *R*_*i*_ matrix has a dimension of *n* × *n* independent of the number of genes annotated in the species. Since the corresponding column of each *D*_*i*_ matrix represents a similar phenotype, we define an expected correlation matrix across the species:

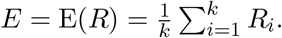

The eigenvalue decomposition of *E* is determined, where *E* = *V* Λ*V* ^*T*^ and the eigenvectors of the *E* matrix form the initial estimates for the different dimensions of the common expression subspace. The right basis matrix *V* is identical in all the *k* species, and the dimensions of *V* represent the correlation relationship between the phenotypes independent of the species. The steps in estimating the common expression subspace are shown in Figure 2. Given the initial estimate of the common expression subspace, we find the best configuration of *V, U*_*i*_, and Δ_*i*_ that minimizes the total reconstruction error 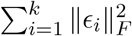 (Additional file 2). We show that our update strategy reduces the factorization error by maintaining the properties of the factors and the optimized *V* represents correlation relationship between the phenotypes (Additional file 2). The columns of the *U*_*i*_ matrix (loading vectors) store the loadings for independent linear combination of genes that transforms gene expression profiles to the *V* space. The genes with significant contributions to the individual dimensions of the common expression subspace can be identified using coefficients (loadings) values in the *U*_*i*_ matrix. The diagonal Δ_*i*_ matrix indicates the amount of correlation information represented by different dimensions of the common expression subspace. In summary, our methodology has easily interpretable factors and employs a systematic strategy to estimate the expression subspace shared by all species.

**Figure 2:**
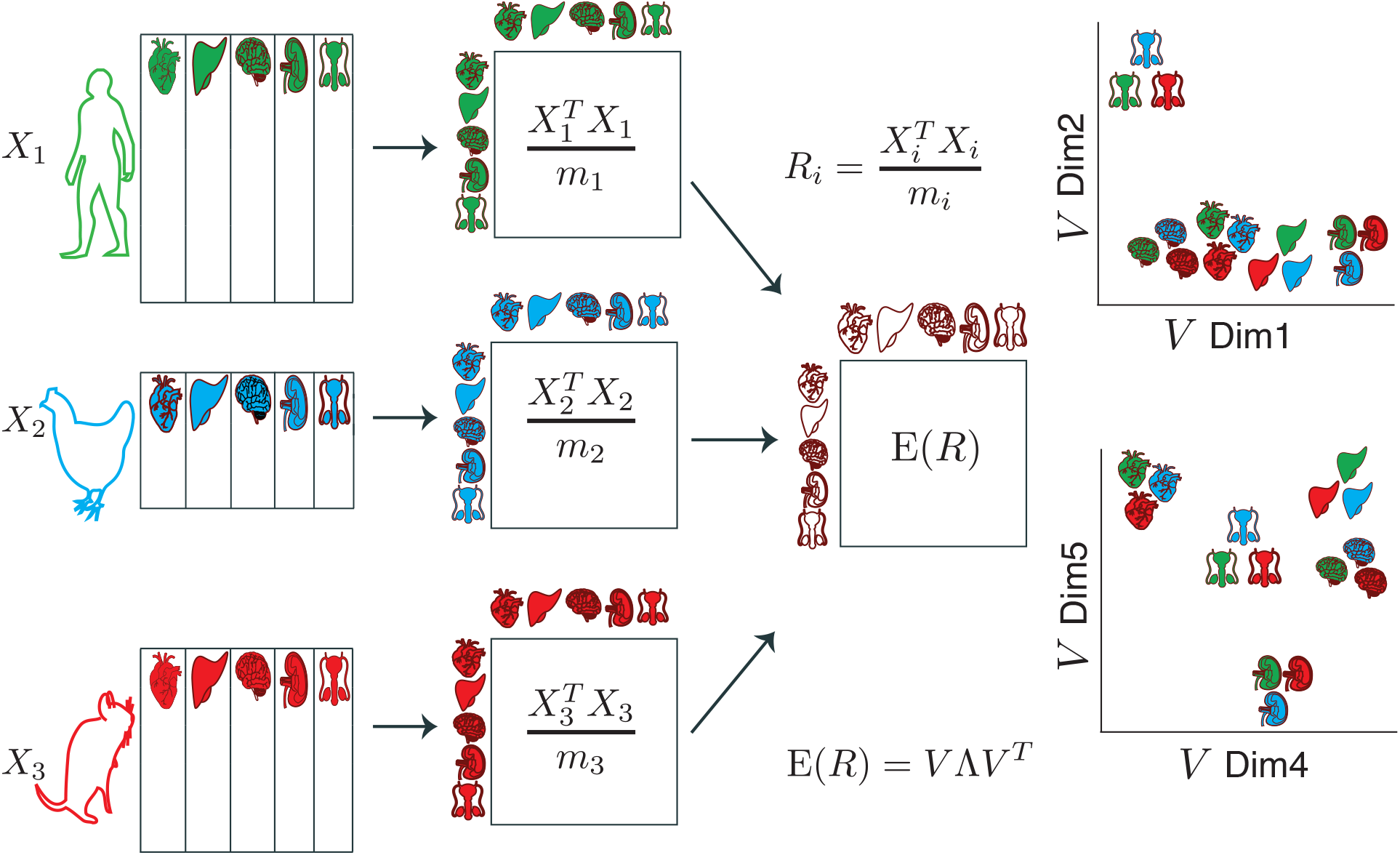
Steps involved in the estimation of common expression subspace: Estimating common expression subspace (*V*) based on gene expression profiles of *n* = 5 tissue types (columns) for three (*k* = 3) species. The *D*_*i*_ matrices are first standardized to obtain the *X*_*i*_ matrices. The inter-tissue correlation within a species (*R*_*i*_) and the expected *R*_*i*_ matrix *E* is then computed. The eigenvectors of the *E* matrix span different dimensions of the expression subspace shared by all species.

### 2.3 Inter-tissue correlation relationship across species

We first examine the inter-tissue correlation within a species (*R*_*i*_ for species *i*) and their relationship across species. We curated a total of 1,018 bulk RNA-Seq profiles from public databases consisting of data from eight vertebrate species, including nine different tissue types (Additional file 3). All the eight species have profiles of six different tissue types (brain, heart, kidney, liver, lung, and testis) in common. Using public RNA-Seq datasets, we compared the *R*_*i*_ relationship across pairs of species (see Methods). The pairwise *R*_*i*_ relationship among a set of species is calculated based on the expression profiles of tissue types present in all the species. When comparing expression profiles of six tissues present in all eight different species, *R*_*i*_ within a species were highly correlated across species (mean *ρ* = 0.90; Additional file 1: Figure S2). Similarly, a correlated pattern (mean *ρ* = 0.83) is observed for the pairwise analysis of five species with nine common tissue types (Additional file 1: Figure S3). The correlated relationship is not observed when the RNA-Seq counts are permuted (Additional file 1: Figure S4). These observations show that the inter-tissue gene expression relationship is similar across species.

### 2.4 Species-independent phenotypic subspace for joint analysis

We applied our joint matrix factorization approach to analyze the RNA-Seq expression profiles of the eight (*k* = 8) species: human, chimp, macaque, mouse, rat, cow, pig, and chicken. The *V* space common to the eight species is estimated using the expression profiles of six tissues (*n* = 6). The estimated *V* space is six-dimensional. The columns of *V*, the right basis vectors (eigenvectors), represent the different dimensions of the common subspace, providing a shared summary of the expression of genes in the six tissues of the eight species. Once the species-specific *U*_*i*_ and Δ_*i*_ matrices are estimated, the joint analysis of gene expression profiles from the eight species is performed by projecting individual expression profiles into the *V* space. A gene expression profile 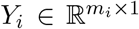 from species *i* is projected to the shared space by computing 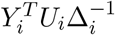. The gene expression libraries from different species projected into the common subspace have identical dimensions (*n* × 1), independent of the number of genes annotated in individual species. Also, the projected profiles in the *V* space cluster by tissue type, independent of species of origin and species-divergence time (Figure 3 a). The species independence and the identical dimension of gene expression profiles in the *V* space enable efficient comparisons across species.

**Figure 3:**
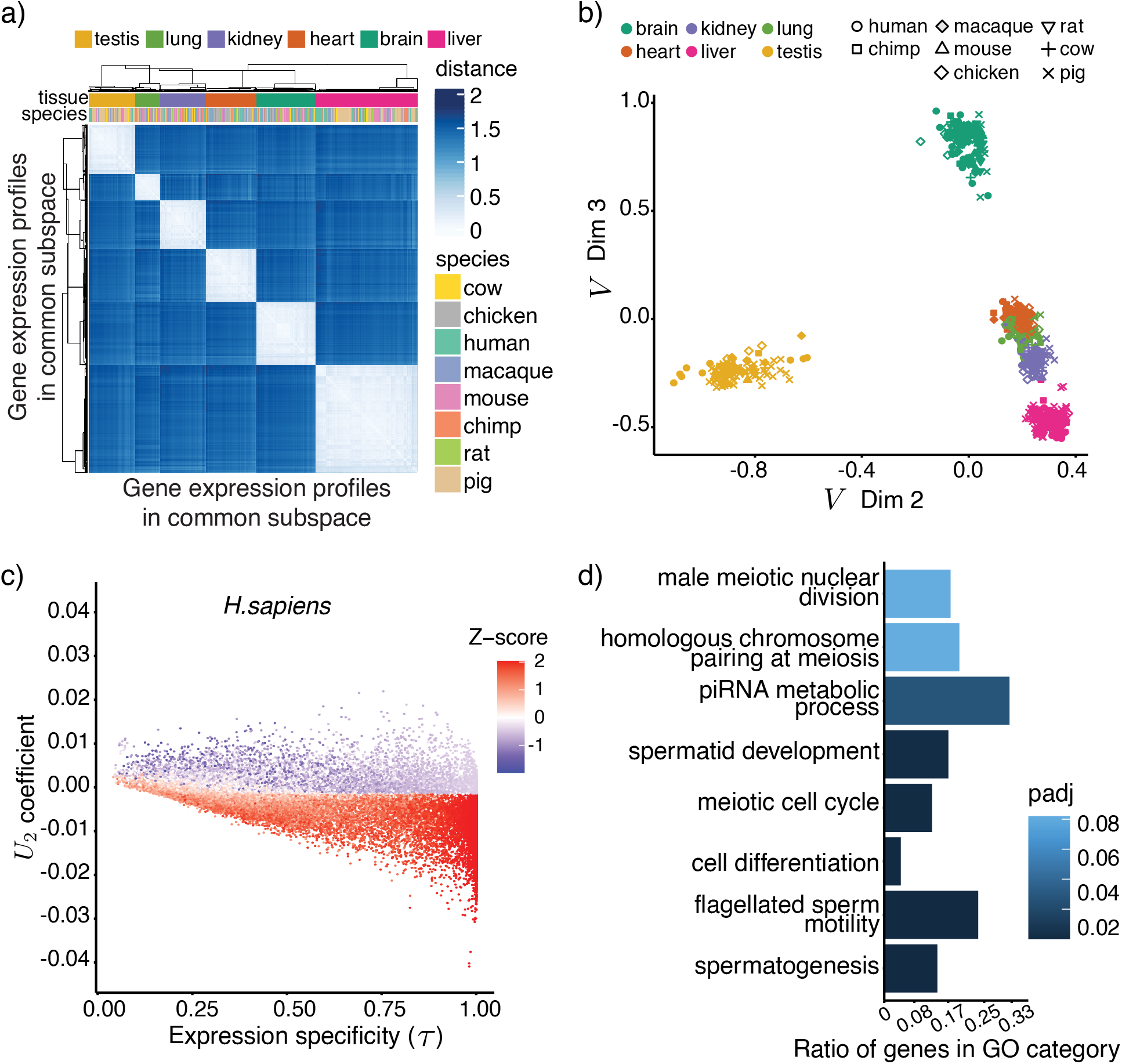
Analysis in the OSBF common subspace: a) Heatmap showing clustering of projected expression profiles in the common subspace. b) The projected expression profiles along dimensions 2 and 3 of the common subspace. c) Scatter plot showing the relationship between gene loadings of *U*_2_ and gene expression specificity (*τ*) for human. Each gene is colored using its testis z-score expression value. d) GO enrichment analysis for top human genes in dimension 2.

#### 2.4.1 Properties of common expression subspace dimensions

We next investigated the properties of different dimensions of the *V* space and the genes contributing to these dimensions. Along dimension 1, the inter-tissue correlation is the highest, resulting in close positioning of the projected expression profiles of different tissue types. (Additional file 1: Table S1, Figure S5 a). The genes with high coefficients in the corresponding column of the loading matrix (loading vector 1) are ubiquitously expressed in all tissues and have low tissue specificity scores (expression specificity: *τ* ≤ 0.25, see Methods, Additional file 1: Figures S5 b-d). The ontology analysis of genes with high loadings in the loading vector 1 shows enrichment for basic cellular functions for different species (Additional file 1: Tables S2, S3). Also, a significant fraction of human housekeeping genes is observed among the human genes with high loadings in loading vector 1 (hypergeometric p-value = 7.29e-12, see Methods). These observations confirm that dimension 1 of the common expression subspace is dominated by housekeeping genes and thus, represents essential cellular processes related to translation, cellular transport, and cell cycle shared in all species. In dimension 2, the testis profiles separate from the rest of the tissues (Figure 3 b). The centroid of the testis profiles is located in the negative axis of the dimension 2, and the genes that show testis-specific expression tend to have high negative loadings in the corresponding loading vector of each species (Figure 3 c, Additional file 1: Figures S6 a-b). The gene ontology (GO) analysis confirms enrichment of similar testis related functions for the top loading vector 2 genes in different species (see Methods, Figure 3 d, Additional file 1: Figures S6 c-d, Table S4).

We similarly identified genes contributing to brain, heart, and kidney phenotype from dimensions 3, 5, and 6, respectively (Additional file 1: Figures S7 a-c, and S8 a-d). Along dimension 4, we observed a weak signature for lung-related genes (Additional file 1: Figures S7 a, d and Table S5). These observations suggest that individual dimensions among the 2-6 of the *V* space represents a similar inter-tissue relationship in all species. Taken together, for all species, each dimension of the common expression subspace represents one or more biological processes related to a column phenotype of the gene expression matrices. The contribution of each gene to the phenotype can be measured using the coefficients in the loading vectors. A summary of cross-species gene expression analysis steps using the OSBF is shown in Additional file 1: Figure S9.

### 2.5 Tissue-relevant genes identified by the OSBF

Using the loadings in the species-specific *U*_*i*_ matrix, we next identified genes with a significant contribution to the tissue phenotype for each species: tissue-relevant genes (TRGs) (see Methods). We focused on the testis, brain, heart, and kidney-specific genes identified from the OSBF’s four dimensions (2, 3, 5, and 6). We first examined the annotation classification of the TRGs, and how many of these genes were characterized by previous studies. The identified genes include protein-coding, non-coding genes, and pseudogenes (Additional file 4). For all the four tissues, nearly 90% of the TRGs are protein-coding in all eight species (Additional file 1: Tables S6, S7). Compared to the fraction of protein-coding genes annotated in different species, the protein-coding genes are over-represented in our TRG list (min hypergeometric p-value ≤ 1.34e-80, Additional file 1: Tables S6, S7). Among the non-coding TRGs, long non-coding RNAs (lncRNAs), microRNAs (miRNAs), and other non-coding RNAs consist of the majority in the brain, heart, and kidney. In contrast, testis non-coding TRGs are mainly lncRNAs and pseudogenes (Additional file 1: Figure S10). Our analysis identified several non-coding genes previously characterized as tissue-specific (Additional file 1: Table S8). The testis TRGs include 56 genes out of the 62 testis-specific genes identified using deep sequencing and immunohistochemistry-based analysis in the Djureinovic et al. (2014) study (hypergeometric p-value = 2.82e-129). Similarly, a significant number of organ-specific genes reported in the human protein atlas (Uhlén et al., 2015) are identified among our brain, heart, and kidney TRGs (Additional file 1: Table S9). In summary, OSBF-based analysis identifies coding as well as non-coding TRGs in different species, including many organ-specific genes identified by previous studies.

#### 2.5.1 Homology-independent identification of tissue-specific expressed genes

The estimation of the OSBF common subspace does not rely on gene orthology mapping between species. Thus, the OSBF-based analysis should be able to identify genes with prominent tissue-specific expression patterns independent of their orthology classification. We first investigated whether our method can identify homologous tissue-specific expressed genes that can be detected with routine cross-species analysis using ortholog mapping information between genes. The OSBF TRGs include species-specific genes as well as orthologs conserved across different species (Figure 4a, Additional file 1: Figure S11). We observed a higher number of orthologs that show tissue-specific expression shared between close species, and chicken has fewer orthologs identified as TRGs (Additional file 1: Table S10). We identified several homologous protein-coding genes that show tissue-specific expression in different species clades and in all eight species (Additional file 1: Tables S11 and S12). Among the well-studied non-coding genes, the homologs of the host gene of *miR-124a* are identified in both human and mouse brain. MicroRNA *miR-124a* is known to be highly expressed in the brain and plays a vital role in hippocampal axonogenesis (Sanuki et al., 2011). The cardiac myocytes and skeletal muscle-specific *miR-133* (Liu et al., 2008) is identified in rhesus, chimpanzee, pig, and chicken heart. MicroRNA-196, a known miRNA enriched in kidney (Meng et al., 2016) is also identified in mouse, pig, and cow. These examples show that the OSBF can detect the homologous TRGs without using orthology-mapping information between species.

**Figure 4:**
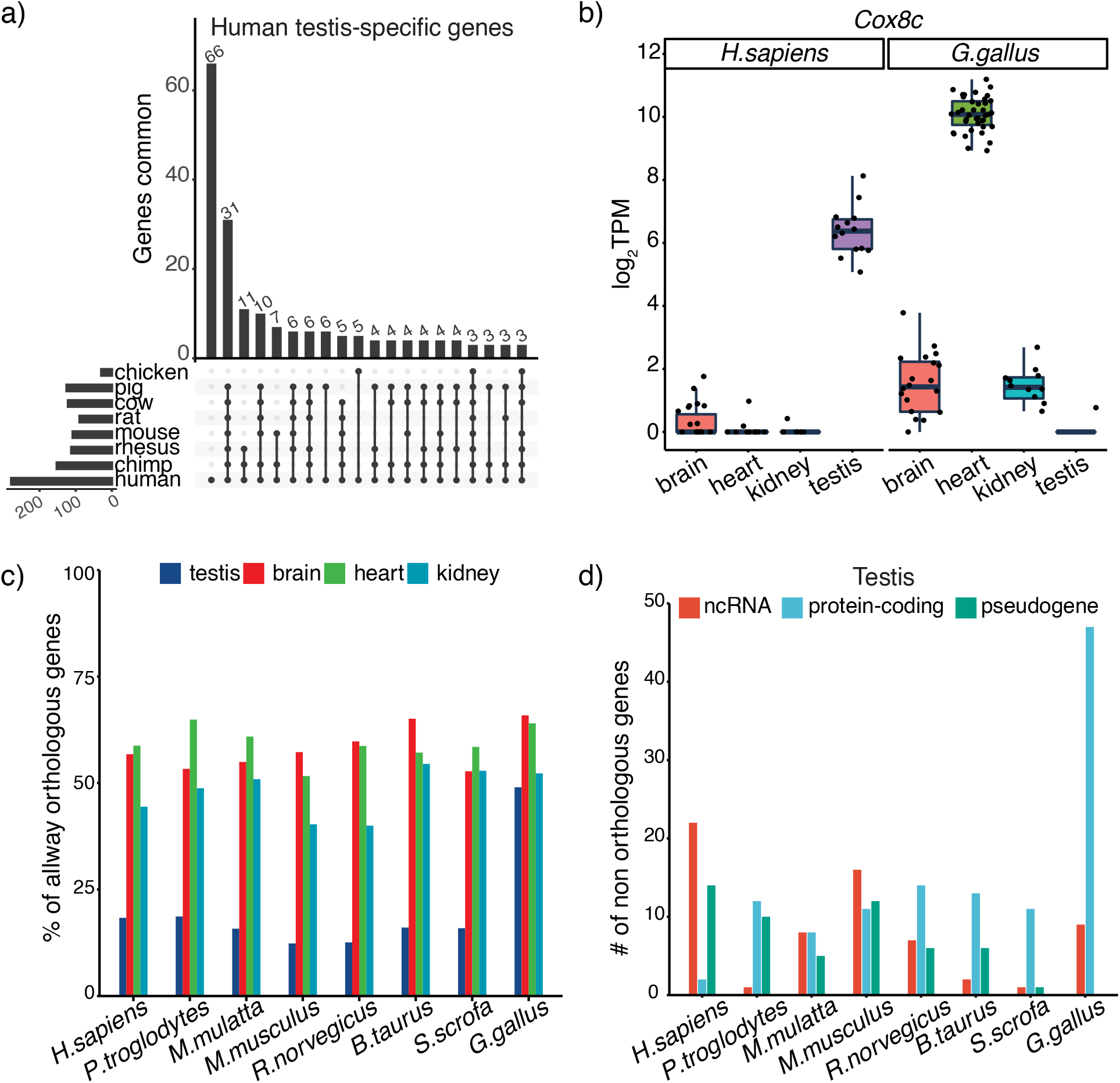
Tissue-relevant genes identified by the OSBF: a) Distribution of testis TRGs unique to human and shared with other species. A human testis-TRG is considered to be shared with a species if the corresponding ortholog is also identified as testis-specific. Only the top 20 intersection sets are shown. b) Among the four tissues, the *Cox8c* gene is highly expressed in the testis in human, while the corresponding ortholog is chicken is over-expressed in the heart. c) The percentage of the OSBF identified TRGs that are shared in all eight species (allway orthologs) in the Ensembl database (v94). More than 40% (on an average testis: 80.15%, brain 41.78%, heart 40.68%, and kidney 51.99%) of the TRGs are not allway orthologous. d) Gene annotation classification of TRGs that do not have any pairwise orthologs in the other seven species in the Ensembl database.

#### 2.5.2 Expression divergence of orthologs

*We examined whether OSBF-based analysis could detect expression divergence of orthologs. We explored those cases where orthologs are identified as TRGs by OSBF but in different organs. In our analysis, Cox8c* gene coding for one of the isoforms of the cytochrome c oxidase subunit is found to be a TRG in the heart of mouse, rat, cow, pig, and chicken. The human ortholog of the *Cox8c* is identified as a TRG in testis but not in the heart (Figure 4b). Hallmann et al. (2016) observed that *Cox8c* transcript is expressed in human testis and three other tissues but not in the heart or muscle. Similarly, *C1qtnf9* is identified to be heart-specific in mouse (Su et al., 2013), while the ortholog in chicken is kidney-specific. We identified three other cases of orthologs with different expression patterns (Additional file 1: Table S13). We also examined the cases where a gene is identified as TRG in one species but not the ortholog in other species. When comparing each species with human, we found that on an average 52% of the brain, 44% of the testis, 36% of the kidney, and 22% of the heart homologous TRGs in human belong to this category (Additional file 1: Figure S12). Similarly, when comparing each species with the mouse, a lower percentage of heart homologous TRGs seems to show expression divergence (Additional file 1: Table S14).

#### 2.5.3 OSBF facilitates discovery of non-homologs contributing to the phenotype

Next, we investigated the non-homologous genes identified by the OSBF that cannot be detected using homology-based analysis. Notably, more than 40% of the TRGs are not one-to-one orthologous (allway orthologous) genes shared in all eight species (Figure 4c). Thus any joint cross-species analysis based on the allway orthologous genes would not be able to identify these genes. Next, we examined whether the identified TRGs have at least one ortholog in the other seven species in the Ensembl database. More than 89% of the TRGs have at least one ortholog in the other seven species. In the category of genes with no reported orthologs with the other seven species, we identified a total of 427 TRGs in eight species (testis 238, brain 55, heart 45, and kidney 88) (Additional file 1: Figure S13 a). Most of these genes are less studied, and any pairwise analysis between species based on orthology mapping would not be able to detect these genes. These genes belong to protein-coding, non-coding RNAs as well as pseudogenes and to be experimentally confirmed annotation classification (Figure 4d, Additional file 1: Figures S13 b-d). Among these genes include previously characterized human testis-specific lncRNA *SPATA42* (Xie et al., 2020) and miRNA *miR-202* (Ro et al., 2007). In mouse testis, genes with no ortholog in other seven species include protein-coding gene *Cypt4* (Kato & Nozaki, 2012), lncRNAs *Rbakdn* (Liu et al., 2021), and pseudogene *1700019M22Rik* (Kato & Nozaki, 2012). The brain-specific genes include scRNA *BCYRN1* (Mu et al., 2020) and lncRNA *MIR124-1HG* (Liu et al., 2016) in human, lncRNAs *Meg3* (Tan et al., 2017), *Miat* (He *et al*., 2020), and *Mir124a-1hg* (Sanuki et al., 2011) in mouse brain. Protein-coding *CEND1* involved in neuron differentiation (Gaitanou et al., 2019) is identified in the chicken brain. Chicken *CEND1* does not have reported ortholog in primates, rodents, and mammals. We identified lncRNAs *LINC00881* (Li et al., 2015), *TRDN-AS1* (Zhang et al., 2018), and pseudogene *NPY6R* (Burkhoff et al., 1998) in human heart. The mouse heart-specific genes with no ortholog in the other seven species include lncRNAs *Mhrt* (Han et al., 2014) and *C130080G10Rik* (Ponnusamy et al., 2019). The kidney-specific non-orthologs include lncRNAs *LINC01187* (Skala et al., 2020), *EMX2OS* (Jiang et al., 2020), and pseudogene *RPS24P17* (Tomaszewski et al., 2015) in the human and mouse lncRNA *D630029K05Rik* (Sanchez-Martin et al., 2021). In addition, 61 organ-specific genes were identified by the OSBF (Ensembl release 94) that did not have any pairwise ortholog in the other seven species have at least one ortholog in the latest annotation release 105 (Additional file 1: Table S15). Taken together, these findings show that OSBF can identify genes contributing to the phenotype independent of their annotation and homology classification.

#### 2.5.4 OSBF identifies tissue-relevant differentially expressed genes

Further, we investigated how TRGs identified by the OSBF can complement within-species differential expression analysis. For each tissue in a species, pairwise differential expression analyses with other tissues were performed, and common differentially expressed (cDE) genes were identified (log fold change ≥ 2, FDR ≤ 0.01, see methods). On average, 93% (brain 97%, heart 94%, testis 94%, kidney 88%) of the TRGs identified by OSBF are cDE genes across eight species (Additional file 1: Table S16). These are the DE genes with relatively high expression, tissue specificity, and gene loadings (Additional file 1: Figure S14 a-c). The remaining TRGs that are not cDE have high expression in two tissues, thus not appearing in all pairwise DE analyses. Interestingly, the DE genes that are also identified as TRGs only constitute a small portion (13%) of the DE genes (Additional file 1: Table S16). So we compared the properties of the DE genes that are also TRGs with those DE genes that are not TRGs and have low gene loadings. For all four tissues, the ontology analysis of non-TRG DE genes with low loadings shows enrichment of biological processes not related to the tissue phenotype (Additional file 1: Table S17). On the contrary, the DE genes that are also TRGs show tissue-related functions (Additional file 1: Table S17). This confirms that *OSBF’s U*_*i*_ loadings help to identify DE genes with significant contribution to the phenotype.

## 3 Discussion

Cross-species analysis of gene expression is challenging as different species have a varying number of genes, and complex relationships exist between genes of two species – a problem dramatically exacerbated when co-analyzing more than two species. The usual strategy to compare gene expression profiles across species is to rely on orthology relationships between genes, assuming functional equivalence of genes across species. The accumulation of evolutionary events such as gene expansions, losses, and other genomic rearrangements complicate the homology relationships between the genes (Lynch & Conery, 2003; Koonin, 2005; Kuzniar et al., 2008) and restricts orthology-based comparisons to evolutionary close species. Furthermore, independent strategies exist in nature to achieve the same molecular function. In this study, we developed a novel mathematical approach called the orthogonal shared basis factorization (OSBF) to analyze gene expression profiles across species. Our method estimates a shared biological space capturing the similar correlation structure across species and enables cross-species comparisons of expression profiles without relying on gene mapping between species.

The conventional cross-species gene expression analysis aims to identify genes important for the phenotypes using a “genes selection - compare phenotype - identify genes” strategy. These approaches first select a subset of shared genes across species and then compares the expression profiles to identify genes essential for the phenotype. In contrast, the OSBF cross-species analysis employs a “phenotype to genes” approach. Our joint matrix factorization starts with gene expression profiles of functionally matching phenotypes (columns) from different species. The correlation relationships between the phenotypes in individual species and their similarity across species are leveraged to estimate a species-independent expression subspace. The common expression subspace can be considered as a subspace representing correlation relationships between functionally similar gene expression profiles in a pseudo-ancestor species. Cross-species analysis based on OSBF uses information from all genes and does not require prior selection of shared genes. Importantly, no prior assumption of functional equivalence of genes across species is made in our analysis. In our estimation procedure, the relative weights of genes in the species-specific loading matrix are distributed such that the genes with high loadings have similar biological functions in all species. Thus, the genes with significant contributions to the different dimensions of the common subspace, and hence to the phenotype can be easily identified.

Using OSBF, we performed a joint analysis of gene expression profiles of six tissues from eight species, including primates, rodents, mammals, and bird. Without relying on orthology mapping between species, we estimated a species-independent six-dimensional expression subspace common for all species. OSBF offers a common coordinate system for cross-species comparisons as gene expression profiles from different species projected into the shared subspace have identical dimensions. We showed that similar biological property is represented along individual dimensions of the common expression subspace, independent of the species and species-divergence time (Figures 3c-d, S5-S8). Genes relevant to organ functions were determined using the gene loadings in species-specific loading matrix. The identified genes include well-studied genes as well as less-studied genes from diverse annotation classifications. Apart from the genes that can be identified using routine cross-species analysis based on orthologs, we identified several tissue-relevant genes that do not have orthologs. In summary, cross-species analysis using OSBF identifies a comprehensive list of genes driving the phenotype independent of their annotation and homology classification.

The cross-species gene expression analysis using the OSBF requires the availability of functionally similar profiles in different species. Finding similar functional profiles from multiple species is challenging as majority of the existing publicly available expression datasets are from a few model organisms. Recently single-cell genomics has made complete transcriptomic profiles of different species available (Fincher et al., 2018; Farrell et al., 2018). This provides an unprecedented opportunity to curate a rich collection of matching cell types across a wide array of species. Extending our approach from tissue to cell type resolution will increase the number of functionally similar profiles and allows cross-species analysis of diverse species using the OSBF.

In OSBF, the common expression subspace is estimated based on the assumption that the inter-tissue relationship within a species is similar across species. In the future, incorporating the phylogenetic effect on the inter-tissue relationships across species could be a potential improvement. Our study uses previously published RNA-Seq datasets from eight species. Our cross-species gene expression analysis is limited due to the differences in tissue extraction and profiling methodology used by different laboratories; confounding effects due to age, sex, and differences in cell population between species.

## 4 Conclusion

Our novel matrix factorization approach enables joint analysis of cross-species gene expression profiles without relying on orthology mapping between species. Genes/transcripts of multiple species driving the phenotype can be identified using OSBF independent of their annotation and homology classification. Homology-independent cross-species comparisons will become very useful as new cell types and tissue types of less-studied species are characterized. We believe that OSBF will be a valuable tool for researchers for cross-species gene expression comparisons.

## 5 Methods

## 5.1 Minimizing factorization error

We devised an iterative algorithm to minimize the total decomposition error. We show that the optimal update is made in each iteration, and the factorization error decreases (Additional file 2). Since the objective function value is bounded below by zero, our procedure will converge to a stable local optimum. Steps for the iterative updates of the factorization are shown below:

### Algorithm

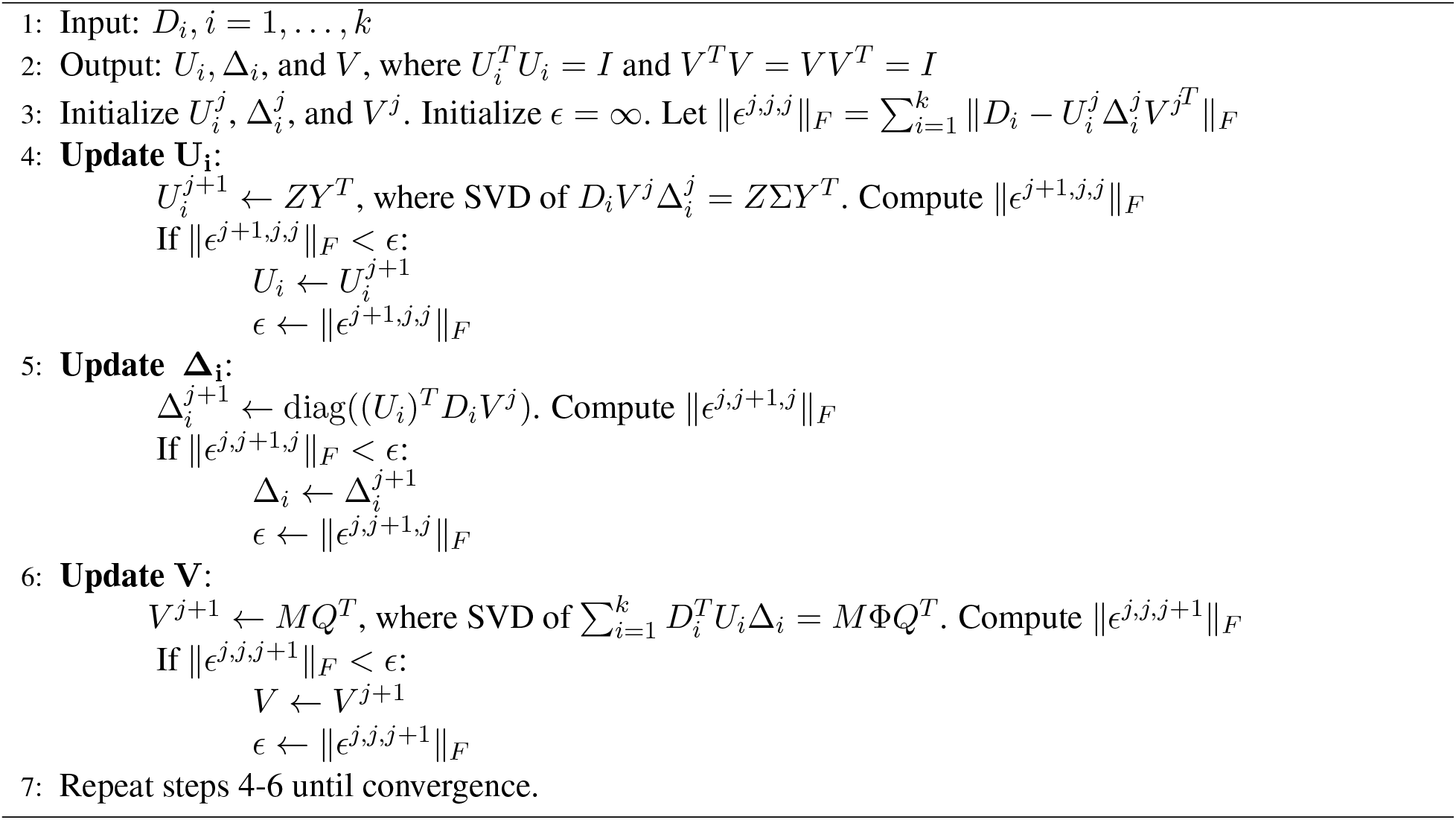

### 5.2 Inter-tissue correlation relationship

The gene expression profiles are first standardized to compute the inter-tissue correlation relationship (*R*_*i*_). Let *X*_*i*_ be the gene expression profiles of species *i* with mean-centered columns and unit variance. We can write

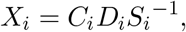

Where

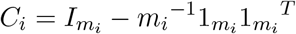

is a centering matrix and *S*_*i*_ = diag(*s*_1_, …, *s*_*n*_) is a diagonal scaling matrix. The standard deviation of *j*-th column of *D*_*i*_ is represented by *s*_*j*_. The profiles from functionally similar tissue types present in all the species are used to calculate the pairwise *R*_*i*_ relationship between species. Inter-tissue gene expression correlation within a species is calculated based on the mean expression of tissue types, and only genes that are expressed in both tissues (mean log_2_(TPM) ≥ 1) is used to calculate the correlation between two tissues in a species. The scatter plot between two species is generated based on corresponding *R*_*i*_ entries (e.g., human brain-heart correlation with mouse brain-heart correlation) from each species. The symmetric and diagonal entries of *R*_*i*_ are excluded. The percentage of correlation information (*p*_*ij*_) represented by a common subspace dimension is defined as 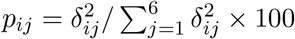,where Δ*i* = diag (*δ*_*i*1_, … *δ*_*i*6_)

## 5.3 Gene annotations and RNA-seq processing

The genome assembly (FASTA file), GTF files, and orthology annotation were obtained from Ensembl v94 (Cunningham et al., 2019). Orthology mapping information was obtained using Ensembl biomaRt (v2.48.3, Durinck et al., 2009). The raw RNA-Seq FASTQ files for eight species were collected from NCBI GEO and EBI ArrayExpress (Additional file 3). Reads were mapped to the corresponding genome using STAR (v2.7.1a) (Dobin et al., 2013) after trimming adapters using Cutadapt (v1.16) (Martin, 2011). The library preparation protocol strand type of each library was inferred using infer experiment.py module in RSeQC (v4.0, Wang et al., 2012). Genes annotated in the GTF files were quantified using HTSeq-count (v0.13.5, Anders et al., 2015). Project-level and species-level quality checks were performed to detect outlier libraries and fix mislabeling if any (Additional file 2).

## 5.4 OSBF gene expression analysis

The OSBF common subspace is estimated using the mean expression profiles of six tissues (brain, heart, kidney, liver, lung, and testis) present in all eight species. The joint-matrix factorization is computed using the ‘SBF’ function call in our R package with parameters ‘orthogonal = TRUE’ and ‘transform matrix = TRUE’ (see R package for details). After projecting the profiles in the common subspace, the heatmap is generated based on clustering (centroid linkage) using the ComplexHeatmap package (v2.8, Gu et al., 2016). Euclidean distance between the projected profiles is used to score the similarity. GO enrichment of top genes is performed using the goseq Bioconductor package (v 1.44 Young et al., 2010) with shared orthologous genes in all eight species as the background. For each species, custom mapping of Ensembl genes to Ensembl GO terms were used for GO enrichment analysis. The human housekeeping gene list is obtained from Eisenberg & Levanon (2013) and Hounkpe et al. (2021). The p-value for overlap between gene sets is calculated using the hypergeometric test. For dimension 1, the genes are ranked based on a score given by the absolute value of the *U*_*i*_ coefficient. For dimensions 2-6, the genes in the corresponding *U*_*i*_ are ranked based on a score (*S*) computed using the *U*_*i*_ coefficient and expression specificity (*τ*) (Yanai et al., 2005). The score for a gene *g* in dimension *i* is defined as 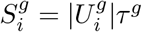, where the *τ* ^*g*^ is the tissue expression specificity index. *τ* ^*g*^ is computed based on the mean expression profile of tissue types. The rank is calculated separately for the genes with positive and negative *U*_*i*_ coefficients. Since the *D*_*i*_ values are non-negative, genes contributing to the tissue of interest will have the same sign in the *U*_*i*_ coefficients as the tissue centroid in the *V* space. The location of the tissue centroids in the common subspace is dependent on the dataset and the estimation parameters. The GO enrichment and overlap with housekeeping genes are performed based on the top 100 ranked genes. The tissue-relevant genes are identified as follows. Using the null distribution of score generated using the OSBF factorization of permuted expression profiles, an empirical p-value is calculated for genes in different dimensions. The genes with significant p-value (≤ 0.001) are identified as genes. The intersection plots are generated using UpSetR package (v1.4, Lex et al., 2014). Tissue-enriched genes from The human protein atlas (https://www.proteinatlas.org) are used for the previously reported tissue-specific markers (Uhlén et al., 2015). In each species, pairwise differential analysis between tissues (testis, brain, heart, and kidney) is performed using DESeq2 (v1.36.0, (Love et al., 2014)) after controlling for the project variable. For each tissue, differentially expressed (DE) genes were identified (log fold change ≥ 2, FDR ≤ 0.01) with respect to the other tissue and those genes identified as DE in all three analyses were used as the common DE genes. All analysis are performed using R (v 4.1.3, R Core Team, 2022). Using different cross-species RNA-Seq datasets, the analysis steps are demonstrated in the package vignettes.

## 6 Code Availability

The OSBF factorization algorithm is implemented as an R package and is available at https://github.com/amalthomas111/SBF under GNU General Public License. The package includes several vignettes demonstrating examples of cross-species gene expression analysis using OSBF. Using public gene expression datasets from different species, the properties of factors before and after applying the optimization procedure, and analysis comparisons of cross-species expression profiles with other matrix factorization approaches are also shown.

## 7 Data Availability

The processed RNA-Seq counts datasets are available at https://doi.org/10.6084/m9.figshare.20216858

## 8 Acknowledgements

The author acknowledges Saket Choudhary, Andrew D. Smith, Guilherme De Sena, Rishvanth Prabakar, Benjamin Decato, Nithin C, and Masaru Nakajima for their helpful discussions and comments on the software and manuscript. The author acknowledges support from the USC Graduate Research Fellowship.

## References

Alter O, Brown PO, Botstein D (2003) Generalized singular value decomposition for comparative analysis of genome-scale expression data sets of two different organisms. Proceedings of the National Academy of Sciences 100:3351–3356.

Anders S, Pyl PT, Huber W (2015) HTSeq a Python framework to work with high-throughput sequencing data. Bioinformatics 31:166–169.

Bergmann S, Ihmels J, Barkai N, Eisen M (2004) Similarities and differences in genome-wide expression data of six organisms. PLoS Biology 2:e9.

Brawand D, Soumillon M, Necsulea A, Julien P, Csárdi G, Harrigan P, Weier M, Liechti A, Aximu-Petri A, Kircher M et al. (2011) The evolution of gene expression levels in mammalian organs. Nature 478:343.

Burkhoff AM, Linemeyer DL, Salon JA (1998) Distribution of a novel hypothalamic neuropeptide Y receptor gene and its absence in rat. Molecular Brain Research 53:311–316.

Cardoso-Moreira M, Halbert J, Valloton D, Velten B, Chen C, Shao Y, Liechti A, Ascenção K, Rummel C, Ovchinnikova S et al. (2019) Gene expression across mammalian organ development. Nature 571:505–509.

Carelli FN, Liechti A, Halbert J, Warnefors M, Kaessmann H (2018) Repurposing of promoters and enhancers during mammalian evolution. Nature Communications 9:4066.

Chen S, Zhang YE, Long M (2010) New genes in Drosophila quickly become essential. Science 330:1682–1685.

Cunningham F, Achuthan P, Akanni W, Allen J, Amode MR, Armean IM, Bennett R, Bhai J, Billis K, Boddu S et al. (2019) Ensembl 2019. Nucleic Acids Research 47:D745–D751.

De Magalhães JP, Curado J, Church GM (2009) Meta-analysis of age-related gene expression profiles identifies common signatures of aging. Bioinformatics 25:875–881.

Djureinovic D, Fagerberg L, Hallström B, Danielsson A, Lindskog C, Uhlén M, Pontén F (2014) The human testis-specific proteome defined by transcriptomics and antibody-based profiling. Molecular Human Reproduction 20:476–488.

Dobin A, Davis CA, Schlesinger F, Drenkow J, Zaleski C, Jha S, Batut P, Chaisson M, Gingeras TR (2013) STAR: ultrafast universal RNA-seq aligner. Bioinformatics 29:15–21.

Durinck S, Spellman PT, Birney E, Huber W (2009) Mapping identifiers for the integration of genomic datasets with the R/Bioconductor package biomaRt. Nature Protocols 4:1184.

Eisenberg E, Levanon EY (2013) Human housekeeping genes, revisited. Trends in Genetics 29:569–574.

Farrell JA, Wang Y, Riesenfeld SJ, Shekhar K, Regev A, Schier AF (2018) Single-cell reconstruction of developmental trajectories during zebrafish embryogenesis. Science 360:eaar3131.

Fincher CT, Wurtzel O, de Hoog T, Kravarik KM, Reddien PW (2018) Cell type transcriptome atlas for the planarian Schmidtea mediterranea. Science 360.

Gaitanou M, Segklia K, Matsas R (2019) CEND1, a story with many tales: from regulation of cell cycle progression/exit of neural stem cells to brain structure and function. Stem Cells International 2019.

Galperin MY, Koonin EV (2012) Divergence and convergence in enzyme evolution. Journal of Biological Chemistry 287:21–28.

Gerstein MB, Rozowsky J, Yan KK, Wang D, Cheng C, Brown JB, Davis CA, Hillier L, Sisu C, Li JJ et al. (2014) Comparative analysis of the transcriptome across distant species. Nature 512:445–448.

Ginis I, Luo Y, Miura T, Thies S, Brandenberger R, Gerecht-Nir S, Amit M, Hoke A, Carpenter MK, Itskovitz-Eldor J et al. (2004) Differences between human and mouse embryonic stem cells. Developmental Biology 269:360–380.

Gu Z, Eils R, Schlesner M (2016) Complex heatmaps reveal patterns and correlations in multidimensional genomic data. Bioinformatics 32:2847–2849.

Hallmann K, Kudin AP, Zsurka G, Kornblum C, Reimann J, Stüve B, Waltz S, Hattingen E, Thiele H, Nürnberg P et al. (2016) Loss of the smallest subunit of cytochrome c oxidase, COX8A, causes Leigh-like syndrome and epilepsy. Brain 139:338–345.

Han P, Li W, Lin CH, Yang J, Shang C, Nurnberg ST, Jin KK, Xu W, Lin CY, Lin CJ et al. (2014) A long noncoding RNA protects the heart from pathological hypertrophy. Nature 514:102–106.

He J, Xue Y, Wang Q, Zhou X, Liu L, Zhang T, Shang C, Ma J, Ma T (2020) Long non-coding RNA MIAT regulates blood tumor barrier permeability by functioning as a competing endogenous RNA. Cell Death & Disease 11:1–18.

Hirano R, Interthal H, Huang C, Nakamura T, Deguchi K, Choi K, Bhattacharjee MB, Arimura K, Umehara F, Izumo S et al. (2007) Spinocerebellar ataxia with axonal neuropathy: consequence of a Tdp1 recessive neomorphic mutation? The EMBO Journal 26:4732–4743.

Hounkpe BW, Chenou F, de Lima F, De Paula EV (2021) HRT Atlas v1. 0 database: redefining human and mouse housekeeping genes and candidate reference transcripts by mining massive RNA-seq datasets. Nucleic Acids Research 49:D947–D955.

Jiang H, Chen H, Wan P, Song S, Chen N (2020) Downregulation of enhancer RNA EMX2OS is associated with poor prognosis in kidney renal clear cell carcinoma. Aging (Albany NY) 12:25865.

Kajala K, Gouran M, Shaar-Moshe L, Mason GA, Rodriguez-Medina J, Kawa D, Pauluzzi G, Reynoso M, Canto-Pastor A, Manzano C et al. (2021) Innovation, conservation, and repurposing of gene function in root cell type development. Cell 184:3333–3348.

Kato Y, Nozaki M (2012) Distinct DNA Methylation Dynamics of Spermatogenic Cell-Specific Intronless Genes Is Associated with CpG Content. PLOS ONE 7:1–10.

Kim SK, Lund J, Kiraly M, Duke K, Jiang M, Stuart JM, Eizinger A, Wylie BN, Davidson GS (2001) A gene expression map for Caenorhabditis elegans. Science 293:2087–2092.

Koonin EV (2005) Orthologs, paralogs, and evolutionary genomics. Annual Review of Genetics 39:309–338.

Koonin EV, Mushegian AR, Bork P (1996) Non-orthologous gene displacement. Trends in Genetics 12:334–336.

Kuzniar A, van Ham RC, Pongor S, Leunissen JA (2008) The quest for orthologs: finding the corresponding gene across genomes. Trends in Genetics 24:539–551.

Lefebvre C, Aude JC, Glémet E, Néri C (2005) Balancing protein similarity and gene co-expression reveals new links between genetic conservation and developmental diversity in invertebrates. Bioinformatics 21:1550–1558.

Lex A, Gehlenborg N, Strobelt H, Vuillemot R, Pfister H (2014) UpSet: visualization of intersecting sets. IEEE Transactions on Visualization and Computer Graphics 20:1983–1992.

Li Y, Lin B, Yang L (2015) Comparative transcriptomic analysis of multiple cardiovascular fates from embryonic stem cells predicts novel regulators in human cardiogenesis. Scientific Reports 5:1–16.

Liu N, Bezprozvannaya S, Williams AH, Qi X, Richardson JA, Bassel-Duby R, Olson EN (2008) microRNA-133a regulates cardiomyocyte proliferation and suppresses smooth muscle gene expression in the heart. Genes & Development 22:3242–3254.

Liu SJ, Nowakowski TJ, Pollen AA, Lui JH, Horlbeck MA, Attenello FJ, He D, Weissman JS, Kriegstein AR, Diaz AA et al. (2016) Single-cell analysis of long non-coding RNAs in the developing human neocortex. Genome Biology 17:1–17.

Liu W, Zhao Y, Liu X, Zhang X, Ding J, Li Y, Tian Y, Wang H, Liu W, Lu Z (2021) A Novel Meiosis-Related lncRNA, Rbakdn, Contributes to Spermatogenesis by Stabilizing Ptbp2. Frontiers in Genetics p. 1963.

Liu Z, Miner JJ, Yago T, Yao L, Lupu F, Xia L, McEver RP (2010) Differential regulation of human and murine p-selectin expression and function in vivo. Journal of Experimental Medicine 207:2975–2987.

Love MI, Huber W, Anders S (2014) Moderated estimation of fold change and dispersion for RNA-seq data with DESeq2. Genome Biology 15:550.

Lynch M, Conery JS (2003) The evolutionary demography of duplicate genes. Genome Evolution pp. 35–44.

Martin M (2011) Cutadapt removes adapter sequences from high-throughput sequencing reads. EMB-net.journal 17:10–12.

Meng J, Li L, Zhao Y, Zhou Z, Zhang M, Li D, Zhang CY, Zen K, Liu Z (2016) MicroRNA-196a/b mitigate renal fibrosis by targeting TGF-β receptor 2. Journal of the American Society of Nephrology 27:3006–3021.

Merkin J, Russell C, Chen P, Burge CB (2012) Evolutionary dynamics of gene and isoform regulation in Mammalian tissues. Science 338:1593–1599.

Mu M, Niu W, Zhang X, Hu S, Niu C (2020) LncRNA BCYRN1 inhibits glioma tumorigenesis by competitively binding with miR-619-5p to regulate CUEDC2 expression and the PTEN/AKT/p21 pathway. Oncogene 39:6879–6892.

Mueller AJ, Canty-Laird EG, Clegg PD, Tew SR (2017) Cross-species gene modules emerge from a systems biology approach to osteoarthritis. NPJ Systems Biology and Applications 3:1–15.

Obayashi T, Kagaya Y, Aoki Y, Tadaka S, Kinoshita K (2019) Coxpresdb v7: a gene coexpression database for 11 animal species supported by 23 coexpression platforms for technical evaluation and evolutionary inference. Nucleic Acids Research 47:D55–D62.

Omelchenko MV, Galperin MY, Wolf YI, Koonin EV (2010) Non-homologous isofunctional enzymes: a systematic analysis of alternative solutions in enzyme evolution. Biology direct 5:1–20.

Ponnapalli SP, Saunders MA, Van Loan CF, Alter O (2011) A higher-order generalized singular value decomposition for comparison of global mRNA expression from multiple organisms. PLOS ONE 6:e28072.

Ponnusamy M, Liu F, Zhang YH, Li RB, Zhai M, Liu F, Zhou LY, Liu CY, Yan KW, Dong YH et al. (2019) Long noncoding RNA CPR (cardiomyocyte proliferation regulator) regulates cardiomyocyte proliferation and cardiac repair. Circulation 139:2668–2684.

R Core Team (2022) R: A Language and Environment for Statistical Computing R Foundation for Statistical Computing, Vienna, Austria.

Ro S, Park C, Sanders KM, McCarrey JR, Yan W (2007) Cloning and expression profiling of testis-expressed microRNAs. Developmental Biology 311:592–602.

Sanchez-Martin I, Magalhães P, Ranjzad P, Fatmi A, Richard F, Manh TPV, Saurin AJ, Feuillet G, Denis C, Woolf AS et al. (2021) Haploinsufficiency of the mouse Tshz3 gene leads to kidney defects. Human Molecular Genetics.

Sanuki R, Onishi A, Koike C, Muramatsu R, Watanabe S, Muranishi Y, Irie S, Uneo S, Koyasu T, Matsui R et al. (2011) miR-124a is required for hippocampal axogenesis and retinal cone survival through Lhx2 suppression. Nature Neuroscience 14:1125–1134.

Segal E, Shapira M, Regev A, Pe’er D, Botstein D, Koller D, Friedman N (2003) Module networks: identifying regulatory modules and their condition-specific regulators from gene expression data. Nature Genetics 34:166–176.

Skala SL, Wang X, Zhang Y, Mannan R, Wang L, Narayanan SP, Vats P, Su F, Chen J, Cao X et al. (2020) Next-generation RNA sequencing–based biomarker characterization of chromophobe renal cell carcinoma and related oncocytic neoplasms. European Urology 78:63–74.

Soucy SM, Huang J, Gogarten JP (2015) Horizontal gene transfer: building the web of life. Nature Reviews Genetics 16:472–482.

Stuart JM, Segal E, Koller D, Kim SK (2003) A gene-coexpression network for global discovery of conserved genetic modules. Science 302:249–255.

Su H, Yuan Y, Wang XM, Lau WB, Wang Y, Wang X, Gao E, Koch WJ, Ma XL (2013) Inhibition of CTRP9, a novel and cardiac-abundantly expressed cell survival molecule, by TNFα-initiated oxidative signaling contributes to exacerbated cardiac injury in diabetic mice. Basic Research in Cardiology 108:1–12.

Sweet-Cordero A, Mukherjee S, Subramanian A, You H, Roix JJ, Ladd-Acosta C, Mesirov J, Golub TR, Jacks T (2005) An oncogenic KRAS2 expression signature identified by cross-species gene-expression analysis. Nature Genetics 37:48–55.

Tan MC, Widagdo J, Chau YQ, Zhu T, Wong JJL, Cheung A, Anggono V (2017) The activity-induced long non-coding RNA Meg3 modulates AMPA receptor surface expression in primary cortical neurons. Frontiers in Cellular Neuroscience 11:124.

Tautz D, Domazet-Lošo T (2011) The evolutionary origin of orphan genes. Nature Reviews Genetics 12:692–702.

Toll-Riera M, Bosch N, Bellora N, Castelo R, Armengol L, Estivill X, Mar Alba M (2009) Origin of primate orphan genes: a comparative genomics approach. Molecular Biology and Evolution 26:603–612.

Tomaszewski M, Eales J, Denniff M, Myers S, Chew GS, Nelson CP, Christofidou P, Desai A, Büsst C, Wojnar L et al. (2015) Renal mechanisms of association between fibroblast growth factor 1 and blood pressure. Journal of the American Society of Nephrology 26:3151–3160.

Uhlén M, Fagerberg L, Hallström BM, Lindskog C, Oksvold P, Mardinoglu A, Sivertsson Å, Kampf C, Sjöstedt E, Asplund A et al. (2015) Tissue-based map of the human proteome. Science 347:1260419.

van Dam S, Craig T, de Magalhães JP (2015) Genefriends: a human rna-seq-based gene and transcript co-expression database. Nucleic Acids Research 43:D1124–D1132.

Van Loan CF (1976) Generalizing the singular value decomposition. SIAM Journal on Numerical Analysis 13:76–83.

Wang L, Wang S, Li W (2012) RSeQC: quality control of RNA-seq experiments. Bioinformatics 28:2184–2185.

Xie Y, Yao J, Zhang X, Chen J, Gao Y, Zhang C, Chen H, Wang Z, Zhao Z, Chen W et al. (2020) A panel of extracellular vesicle long noncoding RNAs in seminal plasma for predicting testicular spermatozoa in nonobstructive azoospermia patients. Human Reproduction 35:2413–2427.

Yanai I, Benjamin H, Shmoish M, Chalifa-Caspi V, Shklar M, Ophir R, Bar-Even A, Horn-Saban S, Safran M, Domany E et al. (2005) Genome-wide midrange transcription profiles reveal expression level relationships in human tissue specification. Bioinformatics 21:650–659.

Yashiro K, Saijoh Y, Sakuma R, Tada M, Tomita N, Amano K, Matsuda Y, Monden M, Okada S, Hamada H (2000) Distinct transcriptional regulation and phylogenetic divergence of human LEFTY genes. Genes to Cells 5:343–357.

Young MD, Wakefield MJ, Smyth GK, Oshlack A (2010) Gene ontology analysis for RNA-seq: accounting for selection bias. Genome Biology 11:R14.

Zhang J, Dean AM, Brunet F, Long M (2004) Evolving protein functional diversity in new genes of Drosophila. Proceedings of the National Academy of Sciences 101:16246–16250.

Zhang L, Salgado-Somoza A, Vausort M, Leszek P, Devaux Y et al. (2018) A heart-enriched antisense long non-coding RNA regulates the balance between cardiac and skeletal muscle triadin. Biochimica et Biophysica Acta (BBA)-Molecular Cell Research 1865:247–258.

