## Additional File 1 for "Orthogonal Shared Basis Factorization: Cross-species gene expression analysis using a common expression subspace"

### Additional File 1: Supplementary figures and tables: Orthogonal Shared Basis Factorization

Amal Thomas

#### Supplementary Figures

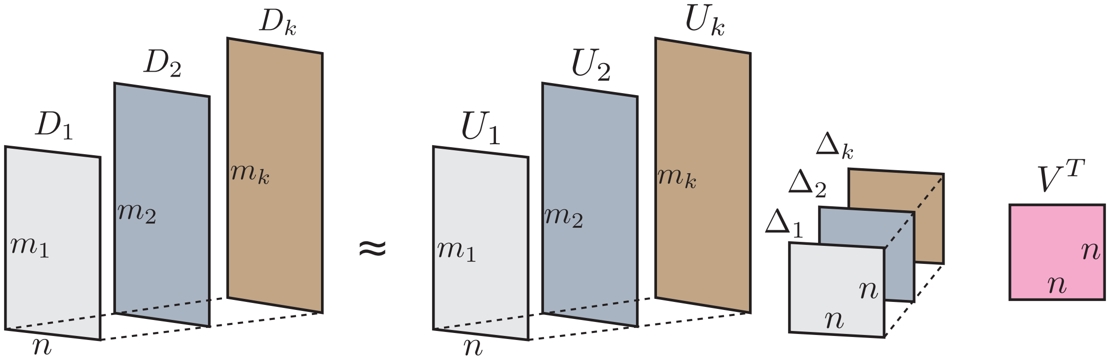


Figure S1: **Individual components of OSBF:** A cartoon summarizing the joint matrix factorization using the OSBF. The dimensions of different matrices are shown using smaller fonts.

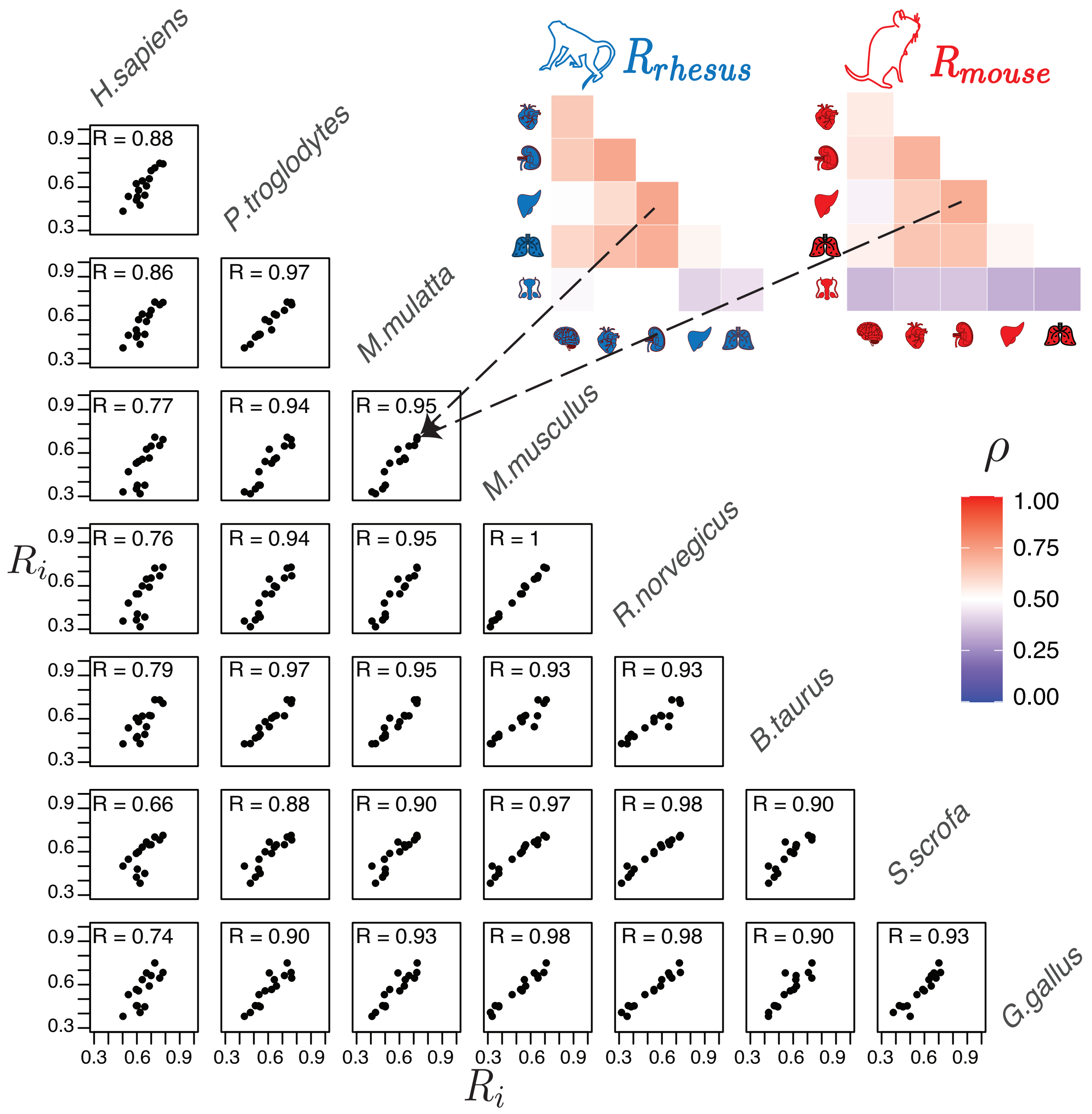


Figure S2: **Inter-tissue gene expression correlation** ($R_i$) **relationship across species:** Pairwise scatter plot shows $R_i$ relationship between eight species: human, chimpanzee, macaque, mouse, rat, cow, pig, and chicken. For each species, $R_i$ is calculated based on six tissues: brain, heart, kidney, liver, lung, and testis. A data point in the scatter plot represents the correlation between the same tissues within a species for two species.

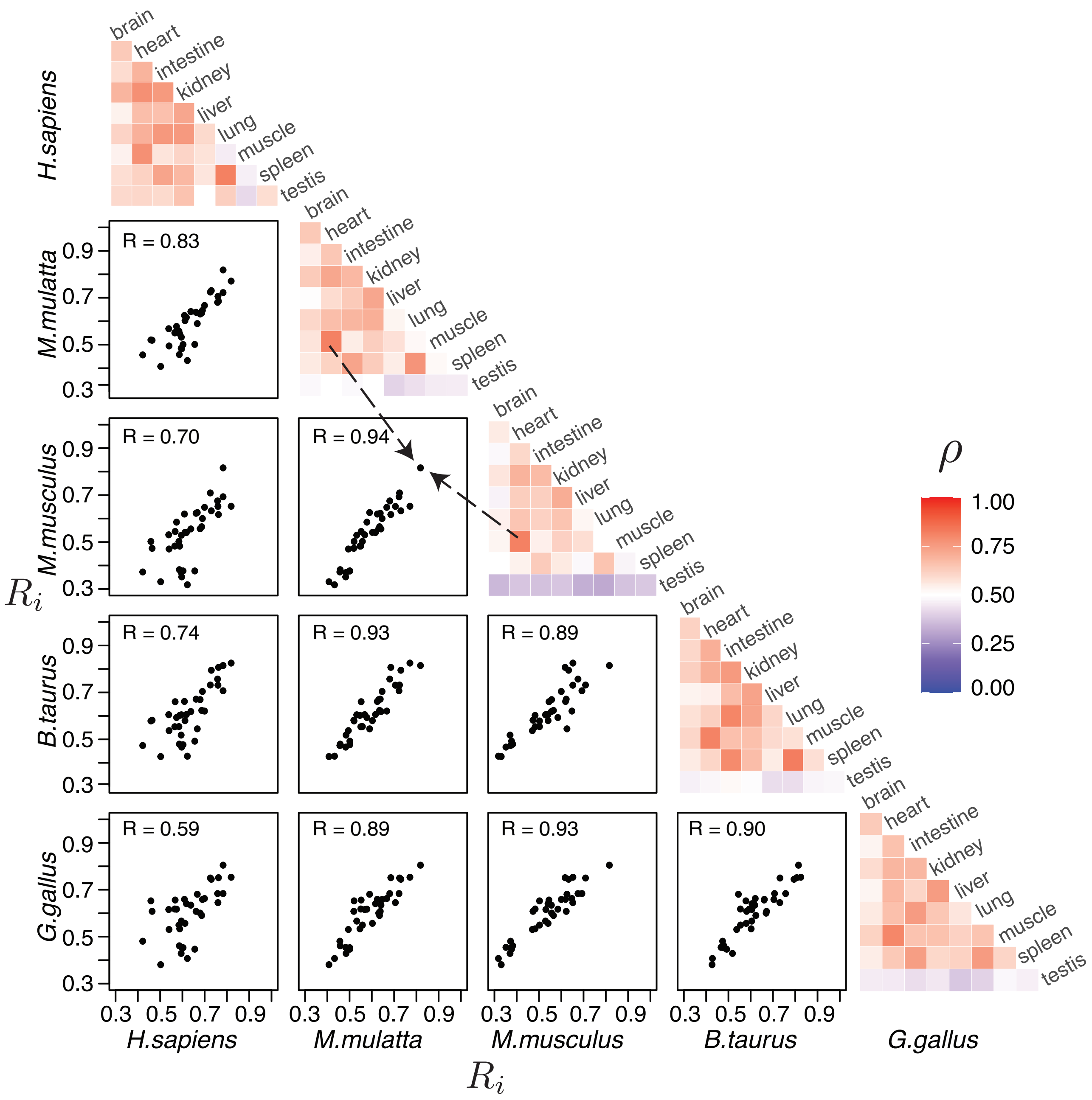


Figure S3: **Inter-tissue gene expression correlation ($R_i$) relationship across species:** Heatmap shows gene expression inter-tissue correlation ($R_i$) within a species for nine different tissue types. Pairwise scatter plot shows $R_i$ relationship between five species: human, macaque, mouse, cow, and chicken. A data point in the scatter plot represents the correlation between the same tissues within a species for two species.

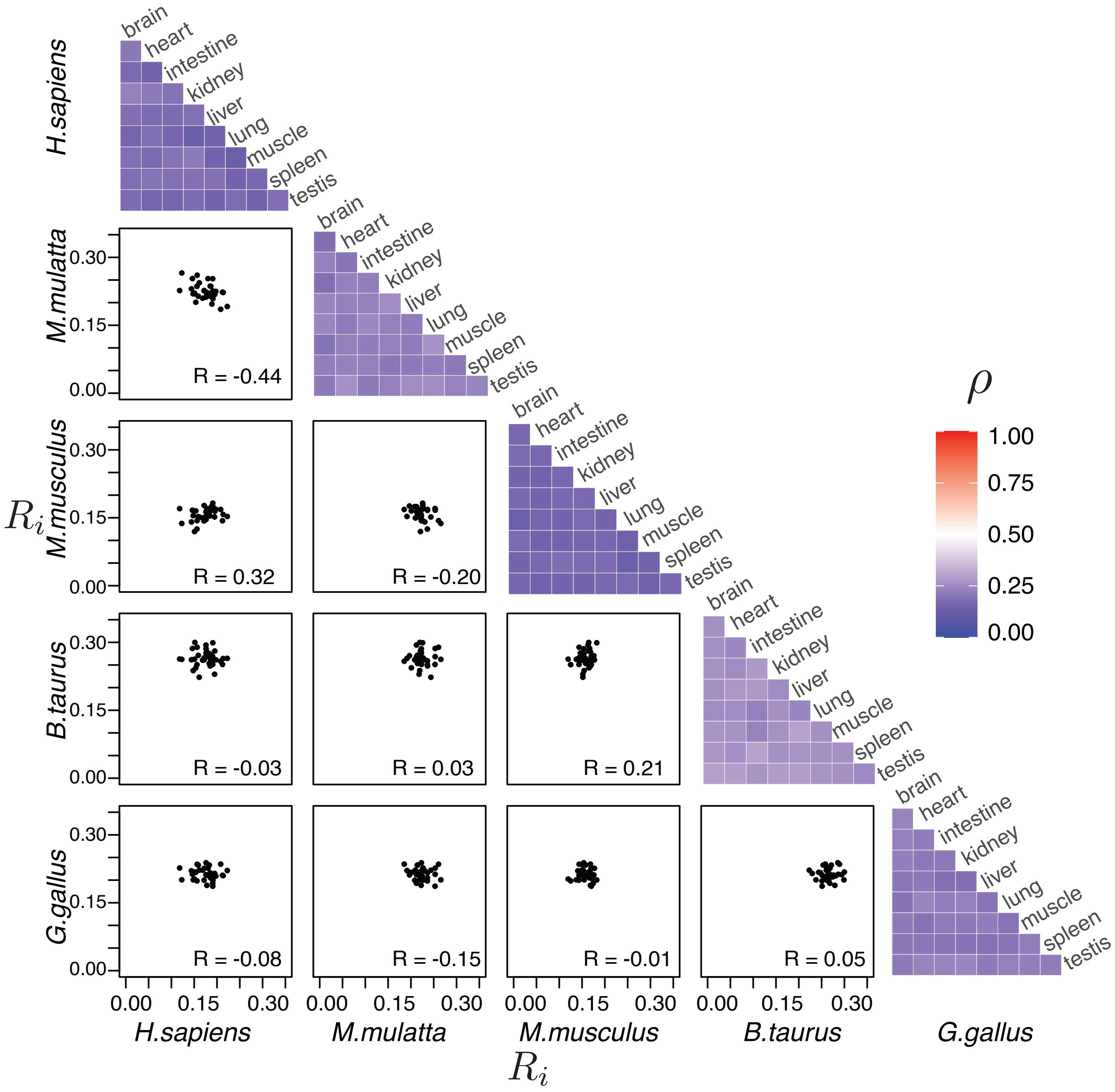


Figure S4: **Inter-tissue gene expression correlation ($R_i$) relationship across species for shuffled counts:** Pairwise inter-tissue gene expression correlation relationship ($R_i$) between five species human, macaque, mouse, cow, and chicken for nine different tissue types when counts of the tissues are permuted to create random expression profiles.

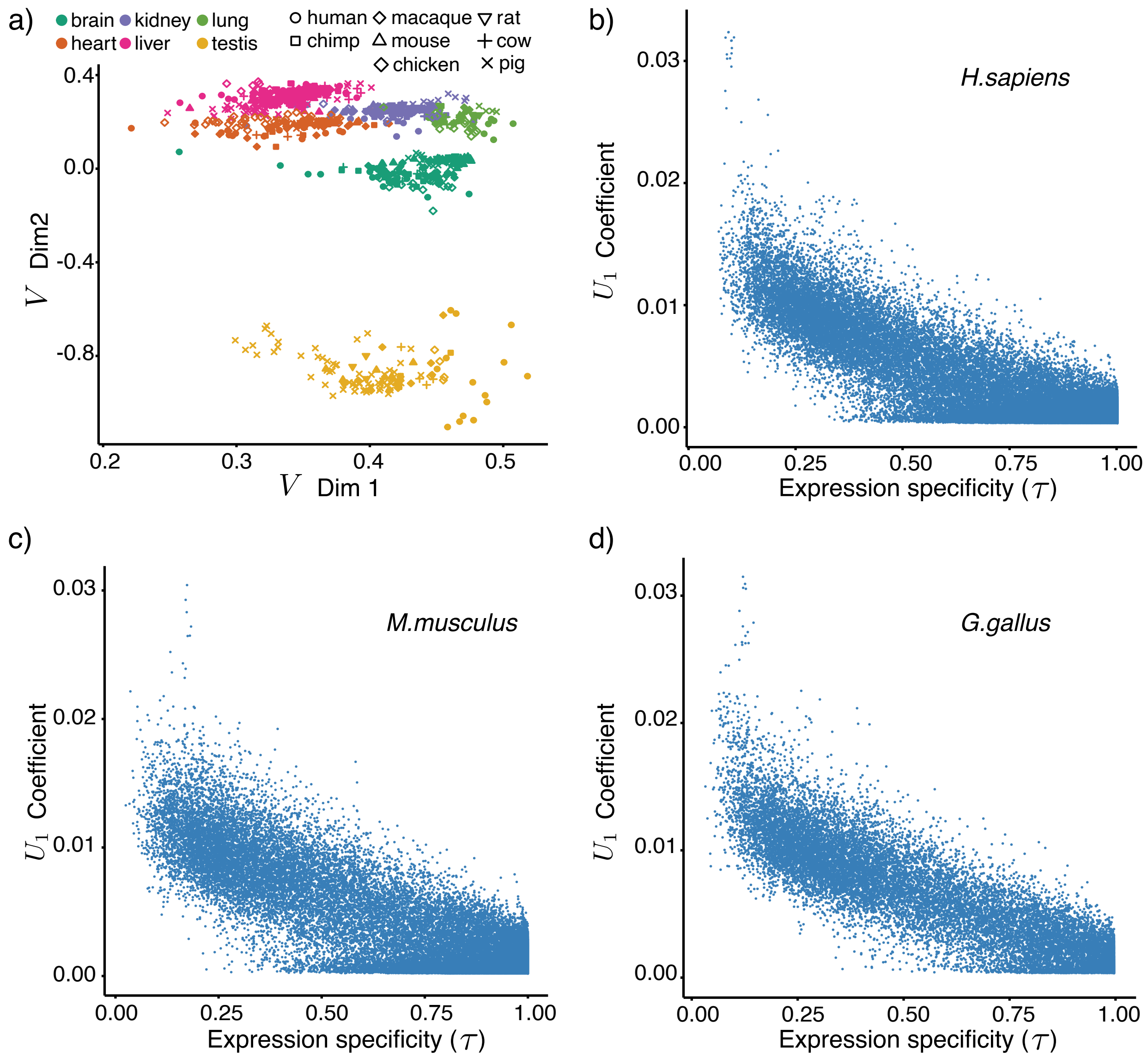


Figure S5: **Properties of the dimension 1 of the common subspace:** a) Gene expression profiles of different tissue types lie close to each other along dimension 1 of the common subspace. b-d) Scatter plot showing the relationship between gene loadings in loading vector 1 ($U_1$) and gene expression specificity ($\tau$) for three different species. All the gene coefficients along dimension 1 are positive values as the centroid of all the tissue types from different species lie on the positive coordinates along dimension 1 of the common subspace (a).

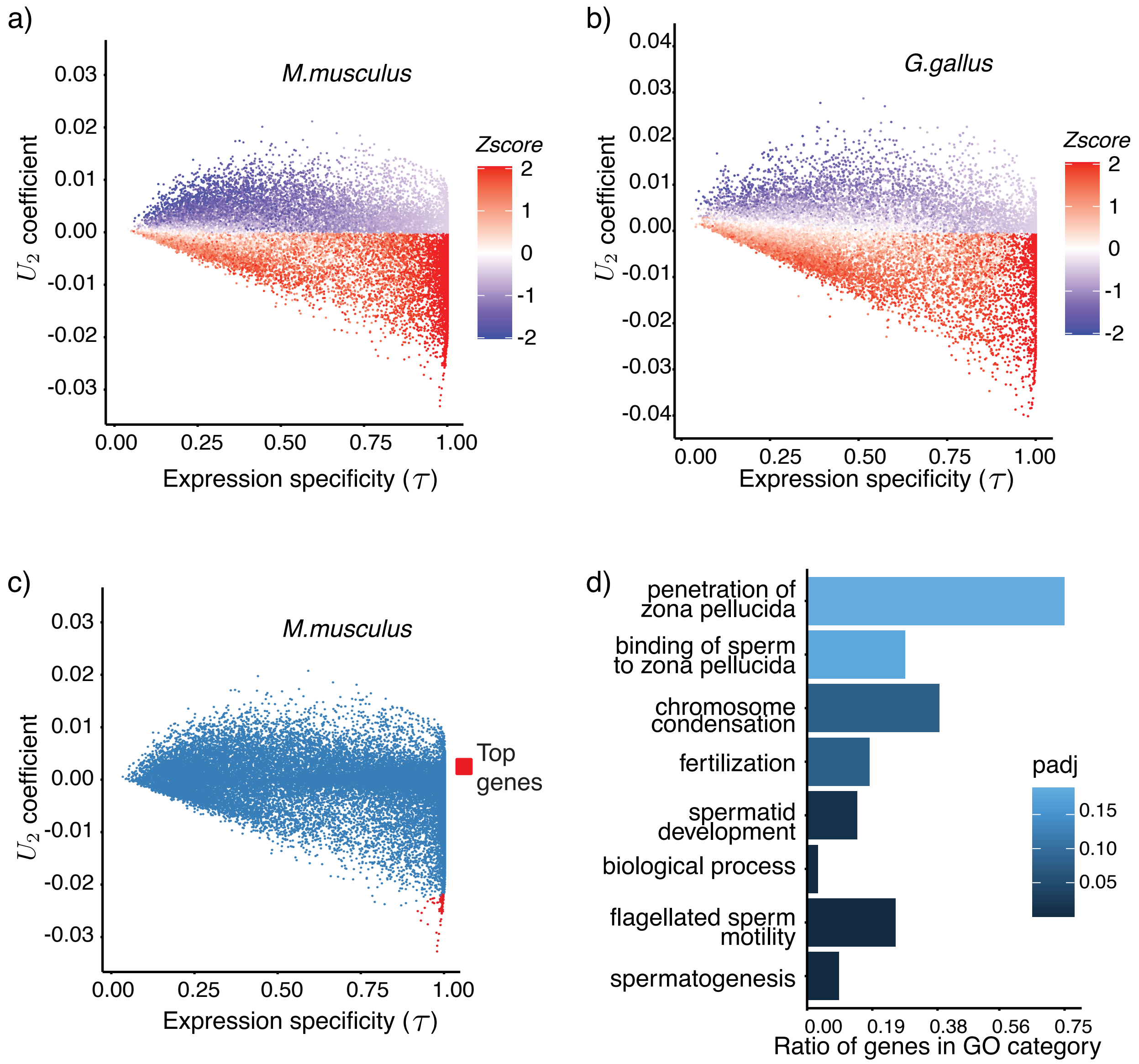


Figure S6: **Properties of the dimension 2 of the common subspace:** a-c) Scatter plot showing the relationship between gene loadings in loading vector 2 ($U_2$) and gene expression specificity ($\tau$) for mouse and chicken. The normalized testis gene expression value for each gene is shown using the z-score values. In (c), the top-ranked genes (see methods) selected for GO analysis (d) for the mouse are highlighted in red. d) The GO ontology analysis of the top mouse genes in dimension 2 shows enrichment for testis-related functions.

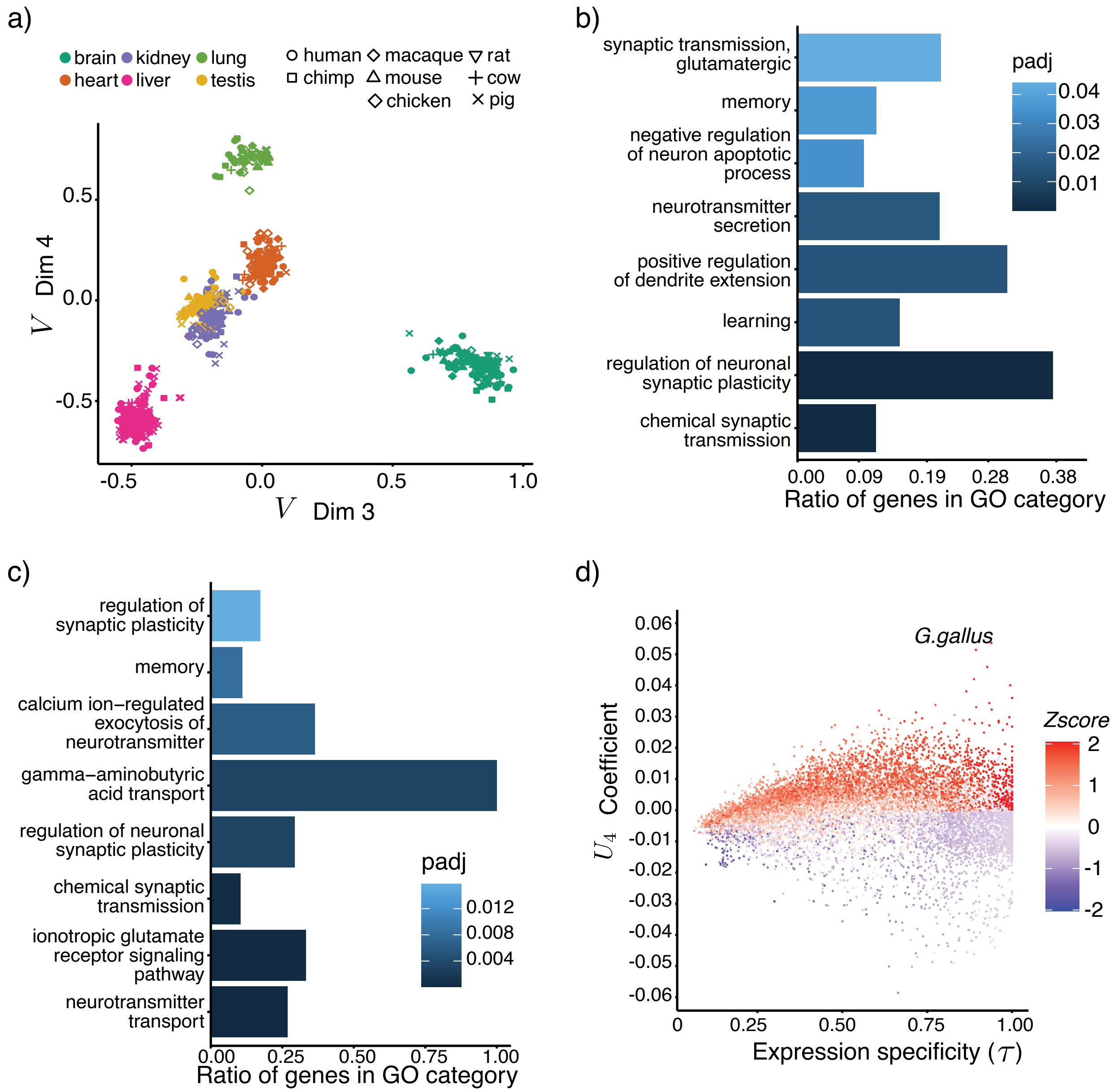


Figure S7: **Properties of the dimensions 3 and 4 of the common subspace:** a) The projected expression profiles along dimensions 3 and 4 of the common subspace. Along dimension 3, a clear separation of the brain tissue from the rest of the tissues is observed. Along dimension 4, a weak separation of the lung tissue is observed. The GO ontology analysis of the top human genes (b) and top mouse genes (c) in dimension three show enrichment for brain-related functions. d) Scatter plot showing the relationship between gene loadings in loading vector 4 ($U_4$) and gene expression specificity ($\tau$) for chicken. The normalized lung gene expression value for each gene is shown using the z-score values.

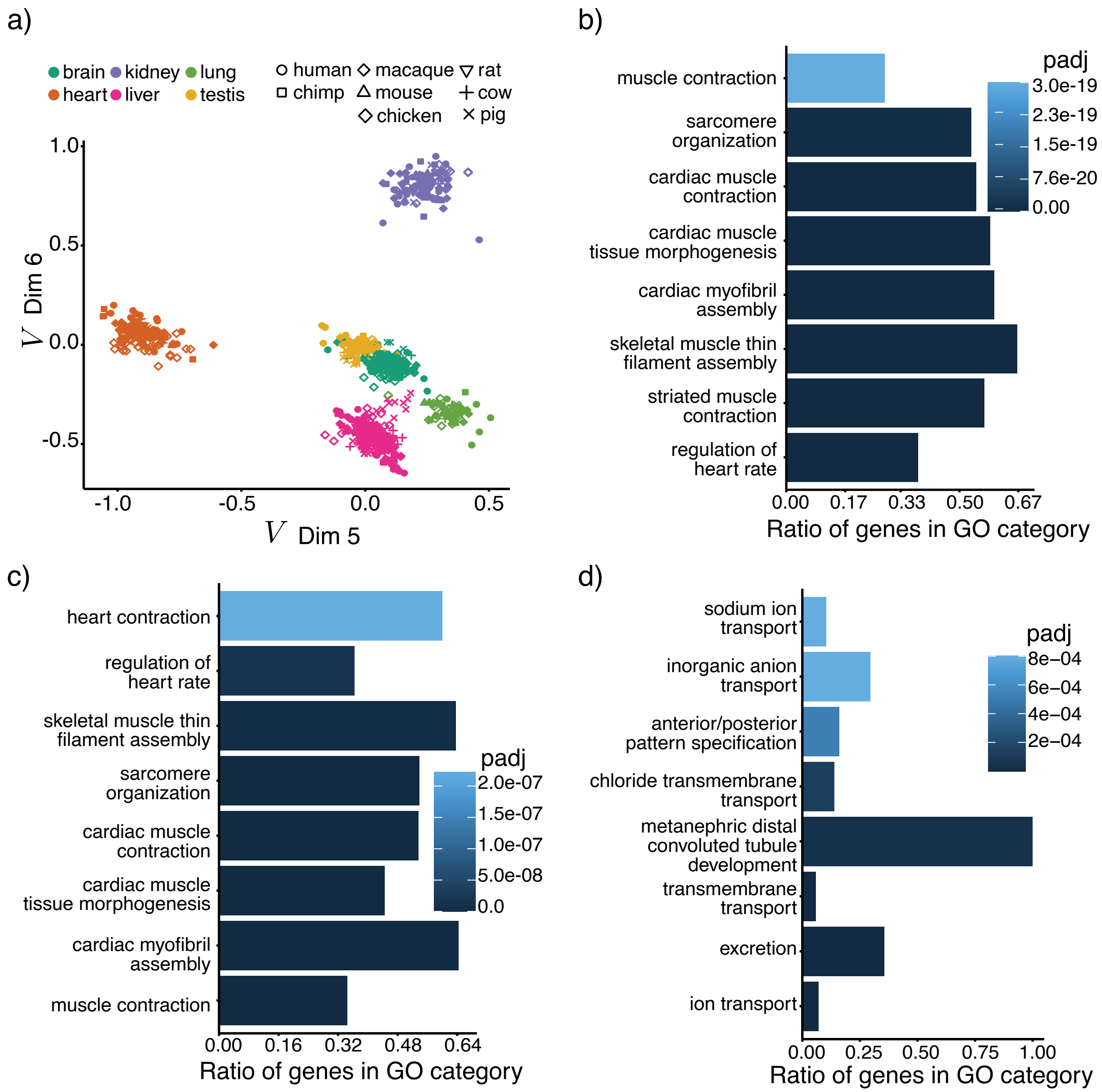


Figure S8: **Properties of the dimensions 5 and 6 of the common subspace:** a) The projected expression profiles along dimensions 4 and 5 of the common subspace. The GO ontology analysis of the top human genes (b) and top mouse genes (c) identified from dimension five show enrichment for heart-related functions. d) The GO ontology analysis of the top human genes in dimension six shows enrichment for kidney-related functions.

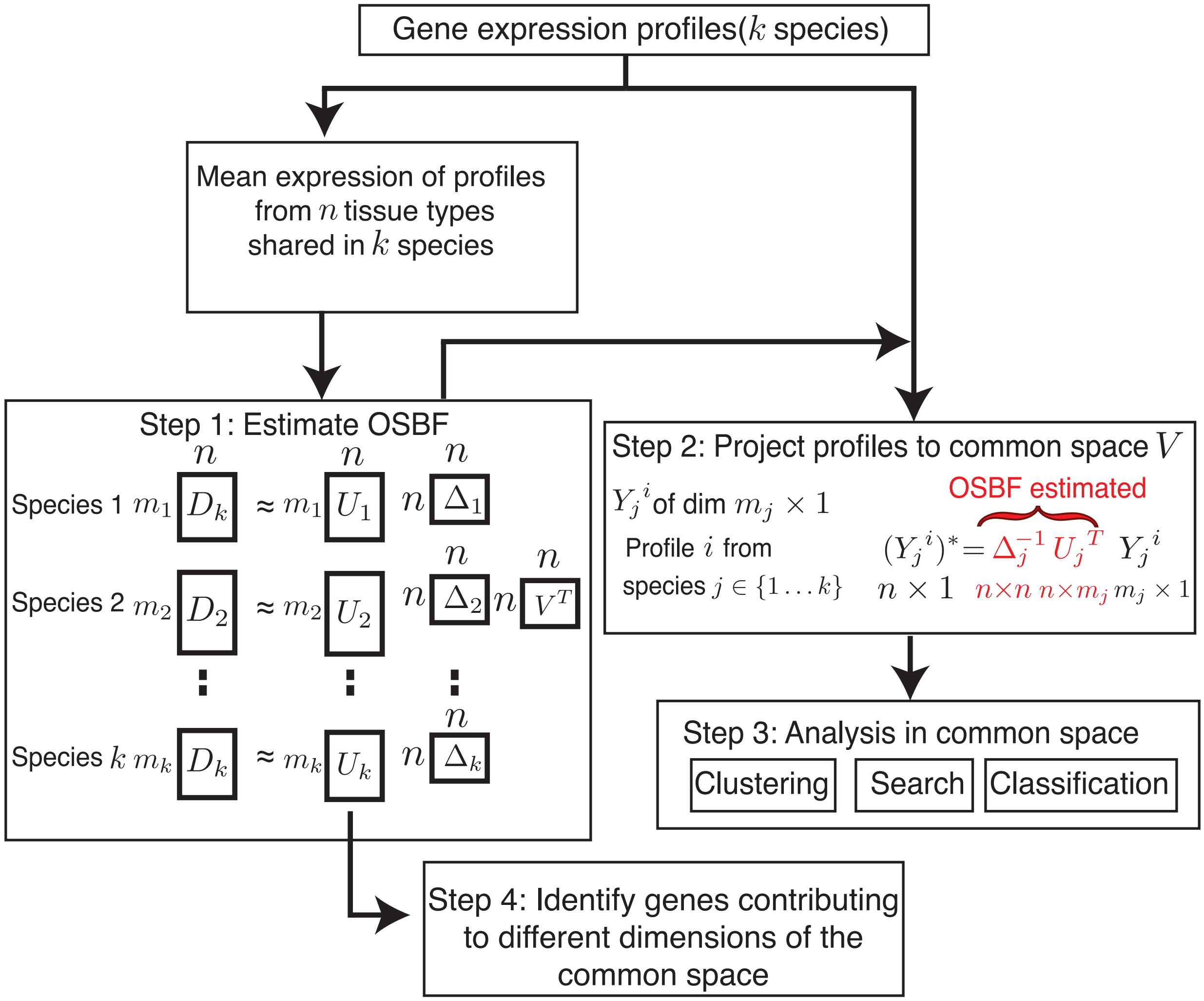


Figure S9: **Steps involved in the estimation of the OSBF common subspace and analysis:** The OSBF matrix factorization is estimated using the mean expression profile of functionally similar tissues present in all $k$ species. All $k$ factorizations share a common basis $V$, a conserved subspace representing the column phenotypes. Individual gene expression libraries from $k$ species are projected on the common subspace for joint analysis where each projected library has a dimension of $n \times 1$.

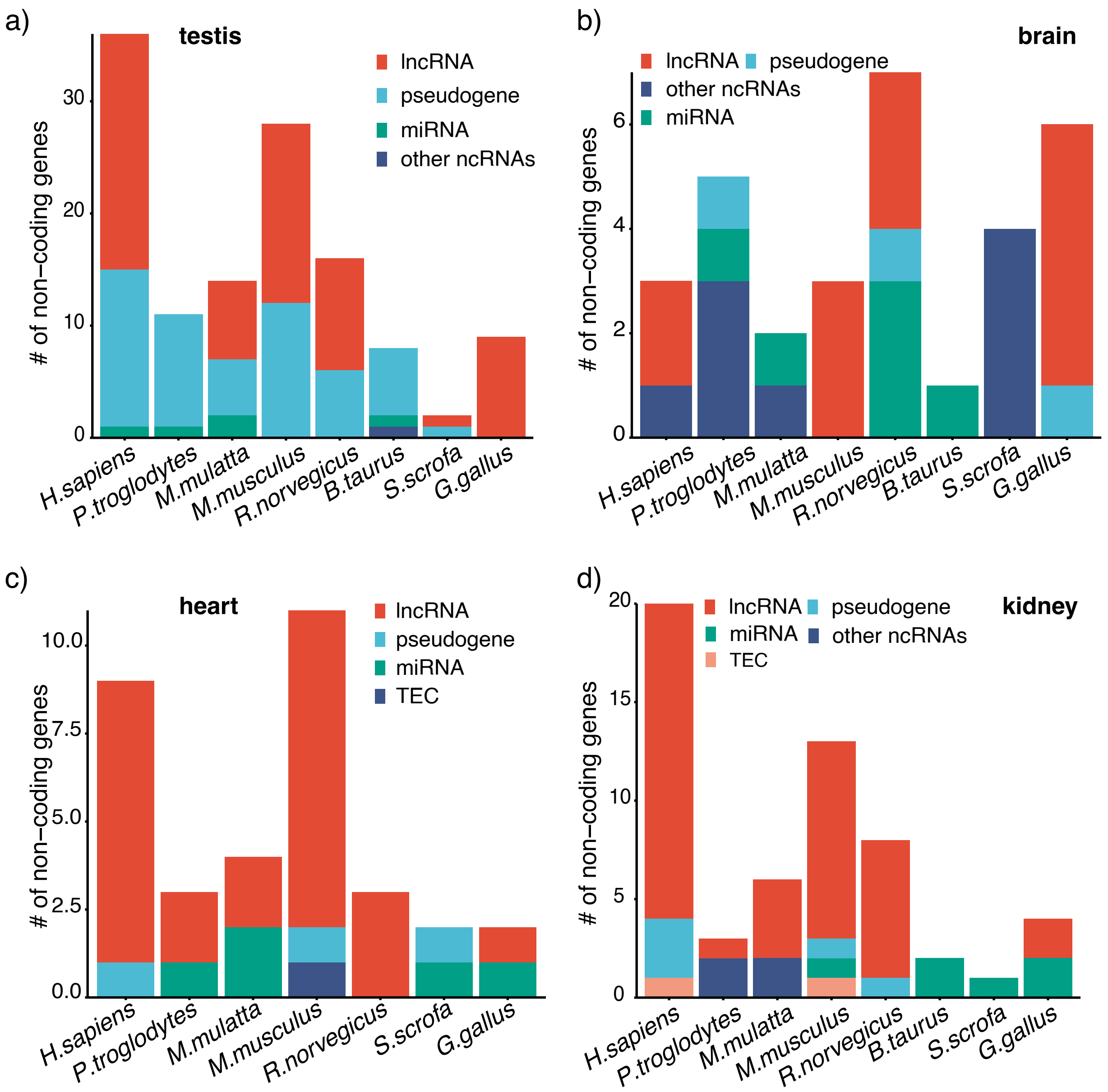


Figure S10: **Gene annotation classification of non-coding genes:** a-d) The number of different non-coding TRGs identified by the OSBF.

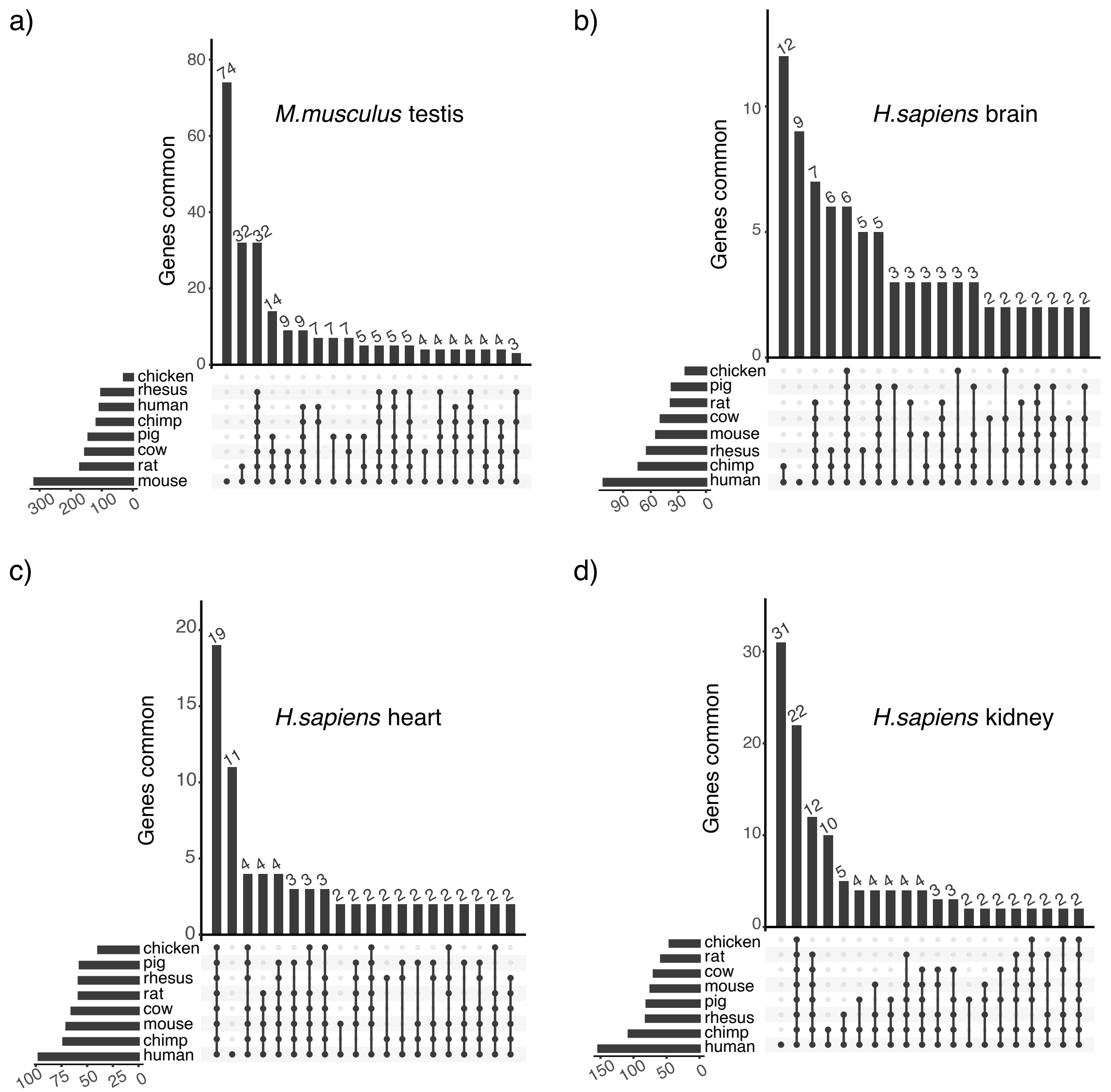


Figure S11: **Distribution of species-specific and shared tissue-relevant genes:** a) Distribution of testis-specific genes that are unique to the mouse and shared with other species. A mouse gene is shared with a species if the corresponding ortholog is identified as testis-specific in the other species. b-d) Human organ-specific genes for brain, heart, and kidney. Similar to a), orthology relationship between human is considered for b-d. Only the top 20 intersection sets are shown for a-d.

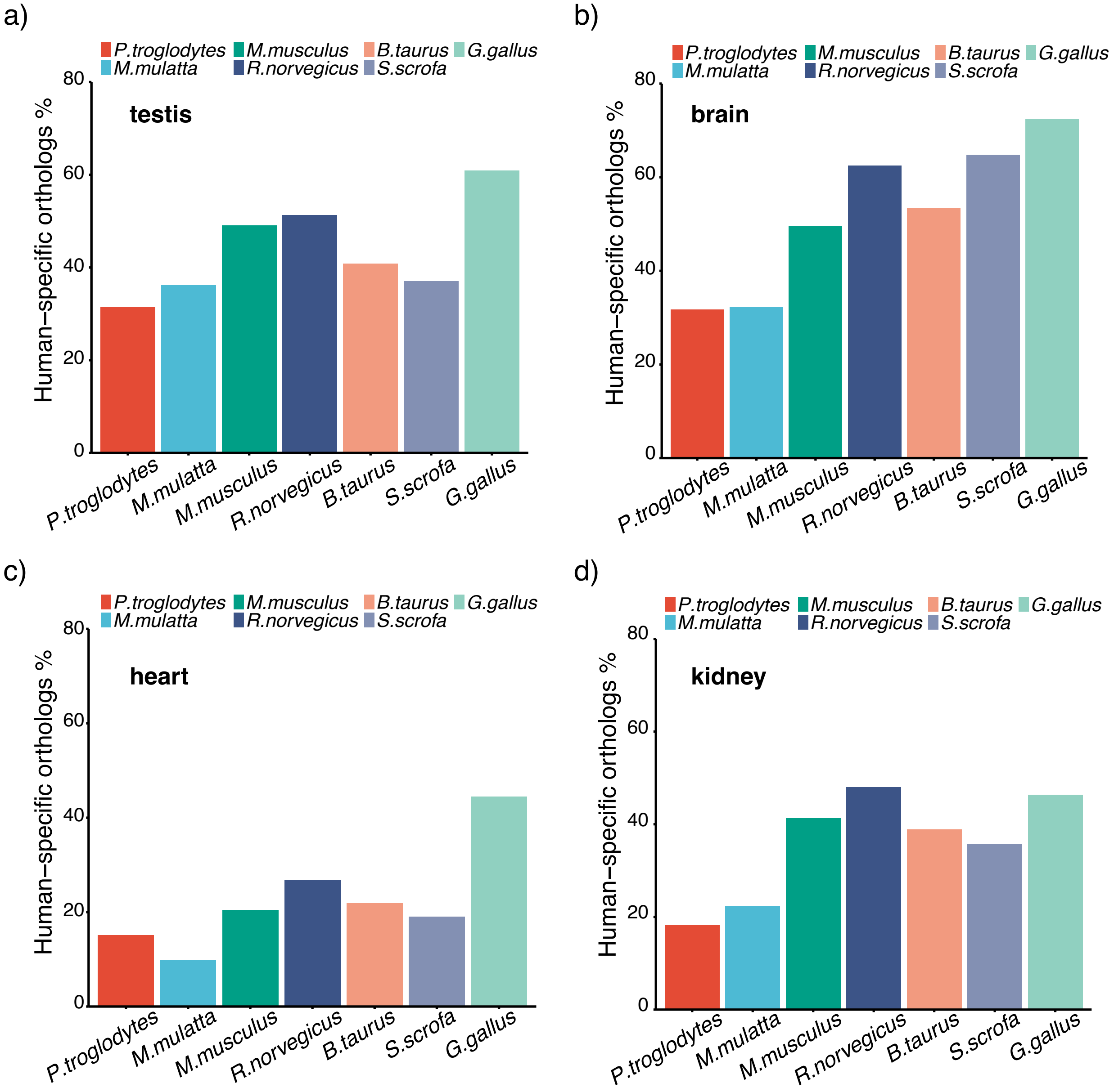


Figure S12: **Human TRGs genes with orthologs not being tissue-specific expressed:** a-d)The percentage of human TRGs identified by OSBF that have orthologs, and the orthologs are not identified as TRGs in the other species.

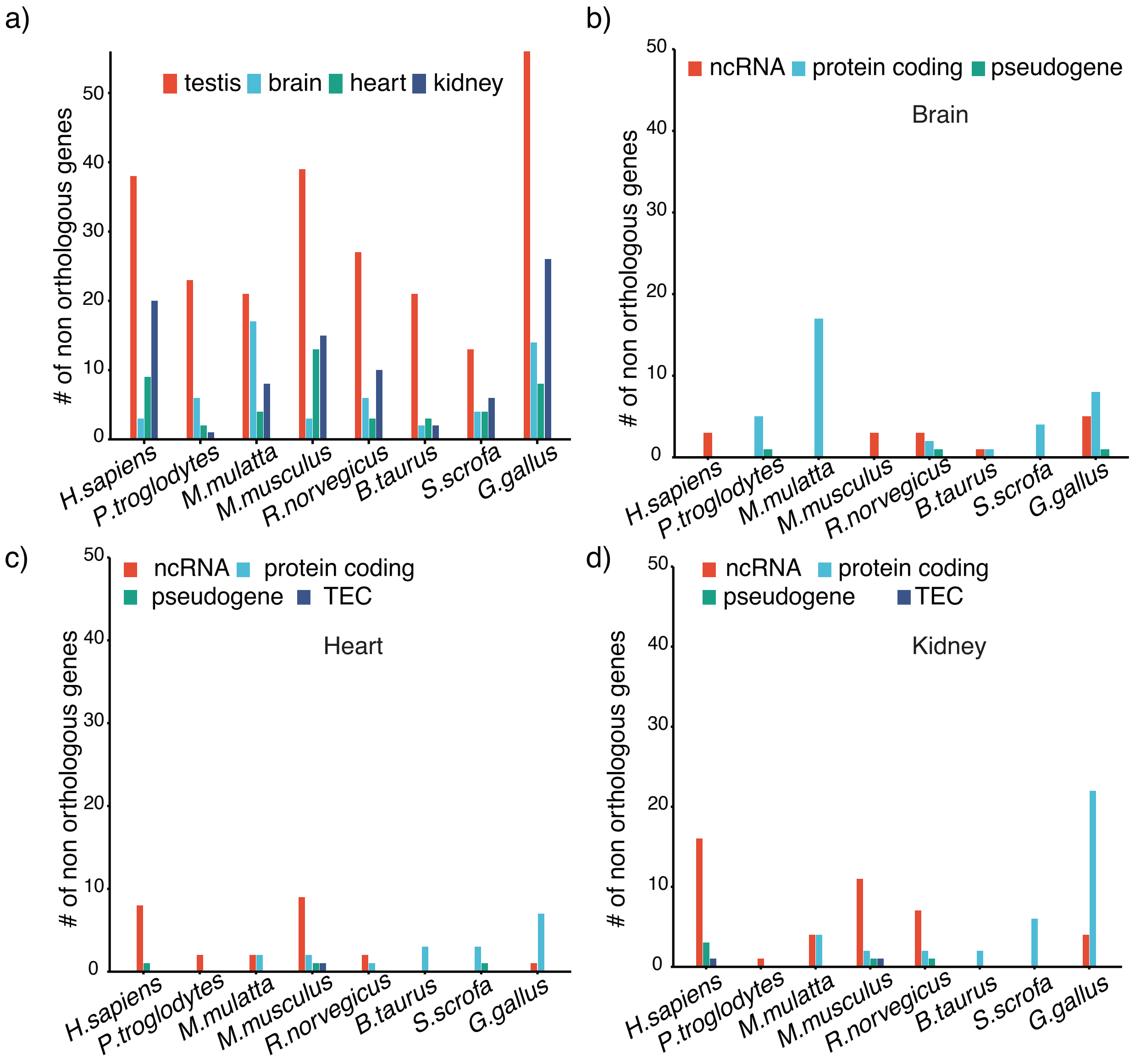


Figure S13: **Tissue-relevant genes with no pairwise orthologs:** a) The number of identified TRGs for different species with no pairwise orthologs with the other seven species in the Ensemble database b-d) Gene annotation classifcation of TRGs with no pairwise orthologs for the brain, heart, and kidney.

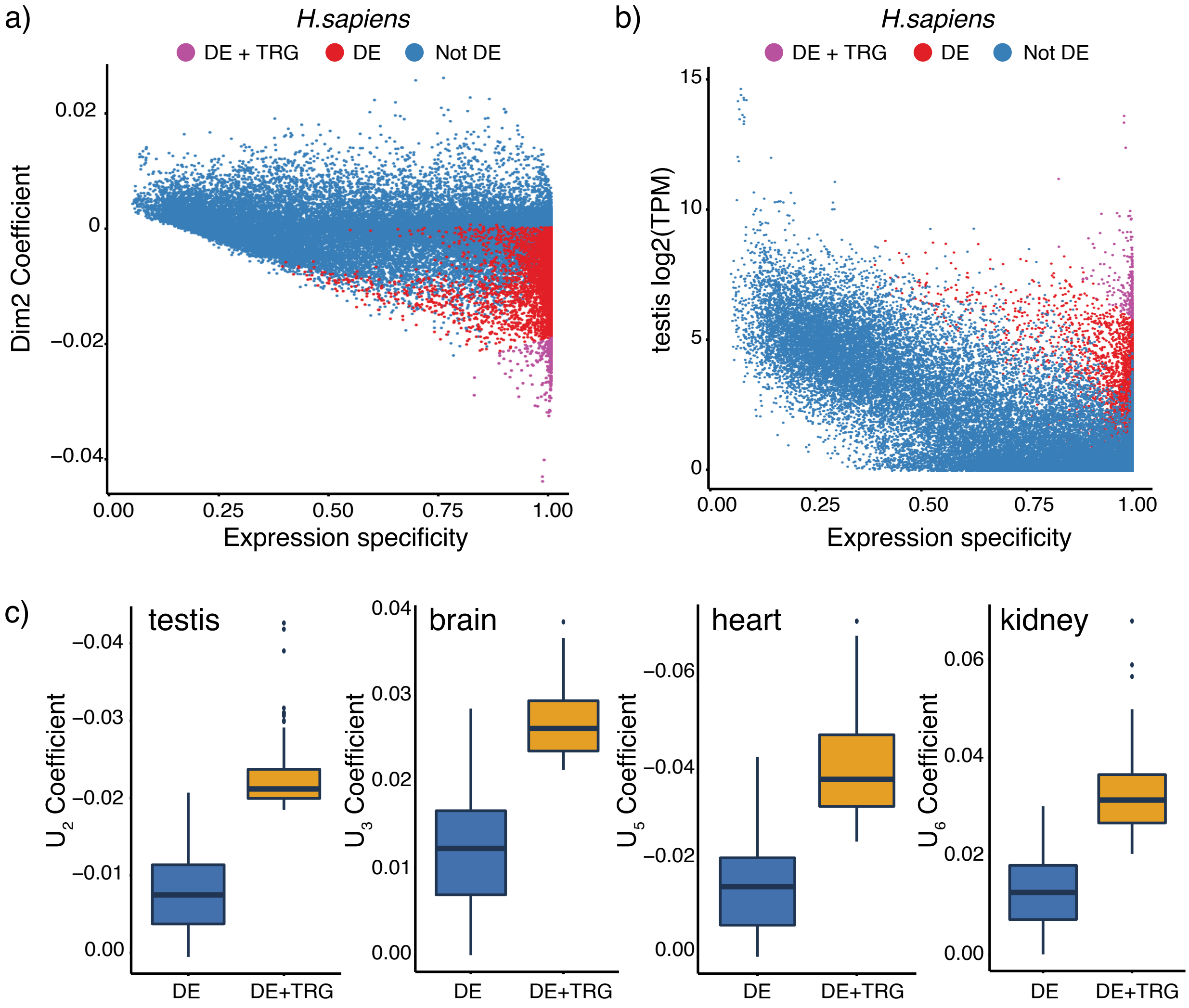


Figure S14: **Differentially expressed and tissue-relevant genes:** Pairwise DE analysis of human testis with other four human tissues is performed (LFC $\geq$ 2, FDR $\leq$ 0.01), and common DE genes were identified. a) Scatter plot showing the relationship between gene loadings of $U_2$ and gene expression specificity ($\tau$) in human. The genes are color based on the DE and TRG status. b) Same as (a) with the x-axis representing actual gene expression values in human testis. c) Distribution of corresponding gene coefficient values of DE and DE + TRGs in human testis, brain, heart, and kidney, respectively.

### Supplementary Tables

Table S1: The percentage of correlation information ($p_{ij}$) represented by the common subspace dimensions.

| | $p_{ij}$ | | | | | | | |
|---|---|---|---|---|---|---|---|---|
| **Dimension** | ***H.sapiens*** | ***P.troglodytes*** | ***M.mulatta*** | ***M.musculus*** | ***R.norvegicus*** | ***B.taurus*** | ***S.scrofa*** | ***G.gallus*** |
| 1 | 87.22 | 88.86 | 87.77 | 81.64 | 84.09 | 87.37 | 85.53 | 88.21 |
| 2 | 5.05 | 4.64 | 4.87 | 7.99 | 6.48 | 4.94 | 5.39 | 4.15 |
| 3 | 2.99 | 2.7 | 2.98 | 4.43 | 3.86 | 2.96 | 3.38 | 2.98 |
| 4 | 2.14 | 1.59 | 1.8 | 2.66 | 2.51 | 1.78 | 2.42 | 1.86 |
| 5 | 1.41 | 1.24 | 1.36 | 1.68 | 1.62 | 1.71 | 1.81 | 1.55 |
| 6 | 1.18 | 0.96 | 1.22 | 1.59 | 1.43 | 1.23 | 1.47 | 1.26 |

$p_{ij} = \delta_{ij}^2 / \sum_{j=1}^{6} \delta_{ij}^2 \times 100,$ where $\Delta_i = \mathrm{diag}(\delta_{i1}, \ldots, \delta_{i6})$

Table S2: Top gene ontology (GO) biological process terms enriched for genes with high $U_1$ coefficient in human.

| **Category** | **Term** | **InCat** | **Total** | **p-adj** |
|---|---|---|---|---|
| GO:0006614 | SRP-dependent cotranslational protein targeting to membrane | 19 | 27 | 9.86E-16 |
| GO:0000184 | mRNA catabolism, nonsense-mediated | 19 | 39 | 8.54E-15 |
| GO:0006413 | translational initiation | 20 | 51 | 9.33E-15 |
| GO:0006412 | translation | 22 | 130 | 1.17E-11 |
| GO:0002181 | cytoplasmic translation | 6 | 9 | 1.19E-3 |

InCat: number of genes of interest in the GO category
Total: total number of genes annotated in the GO category
p-adj: FDR adjusted p-value for over-representation

Table S3: Top gene ontology (GO) biological process terms enriched for genes with high $U_1$ coefficient in mouse.

| **Category** | **Term** | **InCat** | **Total** | **p-adj** |
|---|---|---|---|---|
| GO:0006412 | translation | 14 | 107 | 2.15E-06 |
| GO:0002181 | cytoplasmic translation | 6 | 10 | 3.99E-05 |
| GO:0022900 | electron transport chain | 7 | 26 | 9.79E-05 |
| GO:1902600 | proton transmembrane transport | 7 | 34 | 4.97E-04 |
| GO:0006457 | protein folding | 8 | 49 | 4.97E-04 |

InCat: number of genes of interest in the GO category
Total: total number of genes annotated in the GO category
p-adj: FDR adjusted p-value for over-representation

Table S4: Gene ontology (GO) biological process terms enriched for top genes with high $U_2$ coefficient in different species.

| Category | Term |
|---|---|
| ***Macaca Mulatta*** | |
| GO:0007286 | spermatid development |
| GO:0030317 | flagellated sperm motility |
| GO:0007283 | spermatogenesis |
| GO:0051983 | regulation of chromosome segregation |
| GO:0007341 | penetration of zona pellucida |
| ***Sus scrofa*** | |
| GO:0007283 | spermatogenesis |
| GO:0007286 | spermatid development |
| GO:0030317 | flagellated sperm motility |
| GO:0007342 | fusion of sperm to egg plasma membrane involved in single fertilization |
| ***Gallus gallus*** | |
| GO:0060294 | cilium movement involved in cell motility |
| GO:0030317 | flagellated sperm motility |
| GO:0003341 | cilium movement |
| GO:0007343 | egg activation |
| GO:0007286 | spermatid development |

Table S5: Top gene ontology (GO) biological process terms enriched for genes with high $U_4$ coefficient in humans.

| Category | Term | InCat | Total | p-adj |
|---|---|---|---|---|
| GO:0044267 | cellular protein metabolic process | 11 | 100 | 0.0005 |
| GO:0007585 | respiratory gaseous exchange by respiratory system | 5 | 19 | 0.0184 |
| GO:0043129 | surfactant homeostasis | 4 | 10 | 0.0184 |
| GO:0010811 | positive regulation of cell-substrate adhesion | 5 | 25 | 0.0361 |

InCat: number of genes of interest in the GO category
Total: total number of genes annotated in the GO category
p-adj: FDR adjusted p-value for over-representation

Table S6: Annotation distribution of TRGs identified for testis and brain

| | Testis | | | | Brain | | | |
|---|---|---|---|---|---|---|---|---|
| species | n | protein coding% | non-coding RNAs% | pseudogene% | n | protein coding% | non-coding RNAs% | pseudogene% |
| *H.sapiens* | 278 | 87.05 (7.90e-37) | 7.91 | 5.04 | 111 | 97.30 (2.58e-45) | 2.70 | NA |
| *P.troglodytes* | 343 | 96.79 (1.11e-39) | 0.29 | 2.92 | 135 | 96.30 (1.28e-15) | 2.96 | 0.74 |
| *M.mulatta* | 291 | 95.19 (1.88e-35) | 3.09 | 1.72 | 162 | 98.77 (2.22e-27) | 1.23 | NA |
| *M.musculus* | 316 | 91.14 (1.34e-80) | 5.06 | 3.80 | 138 | 97.83 (6.65e-49) | 2.17 | NA |
| *R.norvegicus* | 302 | 94.70 (4.70e-31) | 3.31 | 1.99 | 82 | 91.46 (3.18e-07) | 7.32 | 1.22 |
| *B.taurus* | 336 | 97.62 (1.01e-20) | 0.60 | 1.79 | 106 | 99.06 (6.49e-09) | 0.94 | NA |
| *S.scrofa* | 358 | 99.44 (1.61e-20) | 0.28 | 0.28 | 127 | 96.85 (6.09e-05) | 3.15 | NA |
| *G.gallus* | 253 | 96.44 (2.45e-22) | 3.56 | NA | 85 | 92.94 (6.15e-06) | 5.88 | 1.18 |

n: Total number of genes identified.
Enrichment p-value (hyper-geometric) shown in brackets for protein-coding genes.

Table S11: Table showing a few ($n = 40$) well-studied protein-coding marker genes identified by the OSBF

| **Gene** | **Biotype** | **Species** | **Reference** |
|---|---|---|---|
| | | **Testis** | |
| ODF1 | protein coding | 7 mammals + chicken | (61) |
| KLHL10 | protein coding | 7 mammals + chicken | (60) |
| TPPP2 | protein coding | 7 mammals + chicken | (39) |
| PRM2 | protein coding | 7 mammals | (58) |
| TNP1 | protein coding | 7 mammals | (35) |
| AKAP4 | protein coding | 7 mammals | (3) |
| SMCP | protein coding | 7 mammals | (21) |
| TCP11 | protein coding | 7 mammals | (34) |
| UBQLN3 | protein coding | 7 mammals | (63) |
| CETN1 | protein coding | 7 mammals | (19) |
| | | **Brain** | |
| GFAP | protein coding | 7 mammals + chicken | (62) |
| SNAP25 | protein coding | 7 mammals + chicken | (52) |
| GAP43 | protein coding | 7 mammals + chicken | (41) |
| GRIN1 | protein coding | 7 mammals + chicken | (43) |
| SYT1 | protein coding | 7 mammals + chicken | (4) |
| PLP1 | protein coding | 7 mammals + chicken | (5) |
| STMN4 | protein coding | 7 mammals | (64) |
| GRIA2 | protein coding | 7 mammals | (6) |
| NEFL | protein coding | 7 mammals | (9) |
| NRSN1 | protein coding | 7 mammals | (27) |
| | | **Heart** | |
| MYL2 | protein coding | 7 mammals + chicken | (48) |
| CSRP3 | protein coding | 7 mammals + chicken | (2) |
| ACTC1 | protein coding | 7 mammals + chicken | (16) |
| MYBPC3 | protein coding | 7 mammals + chicken | (37) |
| MYOZ2 | protein coding | 7 mammals + chicken | (42) |
| TECRL | protein coding | 7 mammals + chicken | (10) |
| MYL3 | protein coding | 7 mammals + chicken | (1) |
| SMYD1 | protein coding | 7 mammals + chicken | (57) |
| TNNC1 | protein coding | 7 mammals + chicken | (29) |
| CASQ2 | protein coding | 7 mammals + chicken | (14) |
| | | **Kidney** | |
| AQP2 | protein coding | 7 mammals + chicken | (51) |
| SLC12A1 | protein coding | 7 mammals + chicken | (36) |
| KCNJ1 | protein coding | 7 mammals + chicken | (28) |
| NPHS2 | protein coding | 7 mammals + chicken | (12) |
| ATP6V0A4 | protein coding | 7 mammals + chicken | (50) |

| | | | |
|---|---|---|---|
| SLC5A12 | protein coding | 7 mammals + chicken | (26) |
| SLC34A1 | protein coding | 7 mammals + chicken | (15) |
| RHCG | protein coding | 7 mammals + chicken | (7) |
| KCNJ16 | protein coding | 7 mammals + chicken | (33) |
| TMEM174 | protein coding | 7 mammals + chicken | (56) |

Table S7: Annotation distribution of TRGs identified for heart and kidney

| | Heart | | | | | Kidney | | | | |
|---|---|---|---|---|---|---|---|---|---|---|
| | n | protein coding% | non-coding RNAs% | pseudogene% | TEC% | n | protein coding% | non-coding RNAs% | pseudogene% | TEC% |
| *H.sapiens* | 97 | 90.72 (1.89e-30) | 8.25 | 1.03 | NA | 153 | 86.93 (1.69e-40) | 10.46 | 1.96 | 0.65 |
| *P.troglodytes* | 94 | 96.81 (9.86e-12) | 3.19 | NA | NA | 164 | 98.17 (2.63e-22) | 1.83 | NA | NA |
| *M.mulatta* | 110 | 96.36 (1.59e-15) | 3.64 | NA | NA | 167 | 96.41 (4.81e-23) | 3.59 | NA | NA |
| *M.musculus* | 122 | 90.98 (1.07e-31) | 7.38 | 0.82 | 0.82 | 191 | 93.19 (4.73e-54) | 5.76 | 0.52 | 0.52 |
| *R.norvegicus* | 92 | 96.74 (3.46e-12) | 3.26 | NA | NA | 140 | 94.29 (1.50e-14) | 5 | 0.71 | NA |
| *B.taurus* | 98 | 100 (1.35e-09) | NA | NA | NA | 156 | 98.72 (5.03e-12) | 1.28 | NA | NA |
| *S.scrofa* | 106 | 98.11 (2.62e-05) | 0.94 | 0.94 | NA | 157 | 99.36 (2.29e-09) | 0.64 | NA | NA |
| *G.gallus* | 75 | 97.33 (4.38e-08) | 2.67 | NA | NA | 155 | 97.42 (1.08e-15) | 2.58 | NA | NA |

n: Total number of genes identified.
TEC: To be experimentally confirmed annotation biotype. Enrichment p-value (hyper-geometric) shown in brackets for protein-coding genes.

Table S8: Table showing a few ($n = 25$) previously studied tissue-specific expressed non-coding genes identified by the OSBF

| Gene | Biotype | Species | Reference |
|---|---|---|---|
| **Testis** | | | |
| SPATA8 | lncRNA | human | (11) |
| MIR202 | miRNA | human | (45) |
| SPATA42 | lncRNA | human | (59) |
| 1700019M22Rik | pseudogene | mouse | (25) |
| Rbakdn | lncRNA | mouse | (32) |
| 4930571K23Rik | lncRNA | mouse | (25) |
| 1700008K24Rik | lncRNA | mouse | (25) |
| **Brain** | | | |
| MIR124-1HG | lncRNA | human | (31) |
| BCYRN1 | scRNA | human | (40) |
| Meg3 | lncRNA | mouse | (53) |
| Miat | lncRNA | mouse | (20) |
| Mir124a-1hg | lncRNA | mouse | (47) |
| **Heart** | | | |
| LINC00881 | lncRNA | human | (30) |
| NPY6R | pseudogene | human | (8) |
| TRDN-AS1 | lncRNA | human | (65) |
| Mhrt | lncRNA | mouse | (18) |
| C130080G10Rik | lncRNA | mouse | (44) |
| mml-mir-133a | miRNA | rhesus | (23) |
| **Kidney** | | | |
| RPS24P17 | pseudogene | human | (54) |
| LINC01187 | lncRNA | human | (49) |
| EMX2OS | lncRNA | human | (24) |
| LINC01874 | lncRNA | human | (13) |
| AP005432.2 | lncRNA | human | (17) |
| D630029K05Rik | lncRNA | mouse | (46) |
| Mir196a-1 | miRNA | mouse | (38) |

Table S9: The number of previously characterized tissue-specific expressed human marker genes identified by the OSBF

| Tissue | OSBF # of genes | Study | Study # of genes | # genes common | p-value |
|---|---|---|---|---|---|
| Testis | 278 | Djureinovic et. al. 2014 | 62 | 56 | 2.82E-129 |
| Testis | 278 | Human protein atlas | 911 | 216 | 0 |
| Brain | 111 | Human protein atlas | 517 | 46 | 1.07E-67 |
| Heart | 97 | Human protein atlas | 34 | 17 | 2.17E-40 |
| Kidney | 153 | Human protein atlas | 56 | 38 | 3.30E-89 |

Human protein atlas: (55) (22)
p-value: Overlap hyper-geometric p-value

Table S10: Percentage of tissue-specific genes with at least one ortholog in other seven species in the Ensembl database identified as TRGs by the OSBF

| | testis | brain | heart | kidney |
|---|---|---|---|---|
| *H.sapiens* | 75.90 | 90.99 | 88.66 | 80.39 |
| *P.troglodytes* | 79.01 | 82.96 | 91.49 | 85.37 |
| *M.mulatta* | 84.54 | 74.07 | 86.36 | 76.05 |
| *M.musculus* | 79.11 | 78.99 | 80.33 | 67.54 |
| *R.norvegicus* | 79.47 | 89.02 | 91.30 | 81.43 |
| *B.taurus* | 82.74 | 88.68 | 87.76 | 77.56 |
| *S.scrofa* | 81.28 | 66.93 | 83.96 | 85.35 |
| *G.gallus* | 35.57 | 57.65 | 70.67 | 48.39 |

Table S12: The number of homologous TRGs genes shared across different species clades

| | primates | rodents | mammals | all 8 species |
|---|---|---|---|---|
| **testis** | 81 | 147 | 32 | 3 |
| **brain** | 47 | 57 | 11 | 7 |
| **heart** | 49 | 67 | 23 | 17 |
| **kidney** | 73 | 83 | 30 | 20 |

Table S13: Expression divergence of orthologs identified by OSBF analysis

| Gene | Species | Tissue | Orthology | Gene | Species | Tissue |
|---|---|---|---|---|---|---|
| ENSG00000187581(COX8C) | *H. Sapiens* | testis | 1 to 1 | ENSMUSG00000025488 | *M.musculus* | heart |
| | | | | ENSRNOG00000014656 | *R.norvegicus* | heart |
| | | | | ENSBTAG00000002046 | *B.taurus* | heart |
| | | | | ENSSSCG00000014560 | *S.scrofa* | heart |
| | | | 1 to many | ENSGALG00000045362 | *G.gallus* | heart |
| ENSMMUG00000019818(CRISP3) | *M.mulatta* | testis | 1 to many | ENSGALG00000034474 | *G.gallus* | kidney |
| ENSPTRG00000023956(SULT1C3) | *P.troglodytes* | kidney | 1 to 1 | ENSSSCG00000028691 | *S.scrofa* | testis |
| ENSSSCG00000011277(CCK) | *S.scrofa* | heart | 1 to 1 | ENSPTRG00000014791 | *P.troglodytes* | brain |
| | | | | ENSMUSG00000032532 | *M.musculus* | brain |
| | | | | ENSBTAG00000013027 | *B.taurus* | brain |
| ENSMUSG00000071347(C1qtnf9) | *M.musculus* | heart | 1 to 1 | ENSGALG00000017117 | *G.gallus* | kidney |

Table S14: Mouse homologous TRGs with corresponding orthologs not identified in the other species

| **tissue** | **reference #total** | **reference #orthologs** | **species** | **# common** | **reference-specific ortholog %** |
|---|---|---|---|---|---|
| testis | 316 | 235 | *H.sapiens* | 108 | 54.04 |
| | | 221 | *P.troglodytes* | 121 | 45.25 |
| | | 209 | *M.mulatta* | 123 | 41.15 |
| | | 259 | *R.norvegicus* | 182 | 29.73 |
| | | 230 | *B.taurus* | 159 | 30.87 |
| | | 240 | *S.scrofa* | 155 | 35.42 |
| | | 84 | *G.gallus* | 34 | 59.52 |
| brain | 138 | 134 | *H.sapiens* | 54 | 59.70 |
| | | 126 | *P.troglodytes* | 55 | 56.35 |
| | | 126 | *M.mulatta* | 62 | 50.79 |
| | | 129 | *R.norvegicus* | 58 | 55.04 |
| | | 129 | *B.taurus* | 61 | 52.71 |
| | | 133 | *S.scrofa* | 49 | 63.16 |
| | | 109 | *G.gallus* | 23 | 78.90 |
| heart | 122 | 108 | *H.sapiens* | 70 | 35.19 |
| | | 108 | *P.troglodytes* | 69 | 36.11 |
| | | 101 | *M.mulatta* | 73 | 27.72 |
| | | 105 | *R.norvegicus* | 74 | 29.52 |
| | | 106 | *B.taurus* | 69 | 34.91 |
| | | 105 | *S.scrofa* | 75 | 28.57 |
| | | 91 | *G.gallus* | 48 | 47.25 |
| kidney | 191 | 160 | *H.sapiens* | 73 | 54.38 |
| | | 156 | *P.troglodytes* | 80 | 48.72 |
| | | 147 | *M.mulatta* | 83 | 43.54 |
| | | 165 | *R.norvegicus* | 91 | 44.85 |
| | | 155 | *B.taurus* | 74 | 52.26 |
| | | 163 | *S.scrofa* | 81 | 50.31 |
| | | 118 | *G.gallus* | 51 | 56.78 |

reference #total: # of mouse TRGs
reference #orthologs: # of mouse homologous TRGs with an ortholog in other species
### common: # of orthologs identified as TSGEs in both mouse and other species
reference-specific ortholog%: % of mouse-specific homologous TRGs

Table S15: The number of OSBF TRGs (Ensembl v94) that do not have any pairwise orthologs in the other seven species but have at least one in Ensembl v105

| **Tissue** | **species** | **#updated in v105** | **Tissue** | **species** | **#updated in v105** |
|---|---|---|---|---|---|
| testis | *H.sapiens* | 3 | heart | *H.sapiens* | 0 |
| testis | *P.troglodytes* | 1 | heart | *P.troglodytes* | 0 |
| testis | *M.mulatta* | 7 | heart | *M.mulatta* | 2 |
| testis | *M.musculus* | 4 | heart | *M.musculus* | 1 |
| testis | *R.norvegicus* | 10 | heart | *R.norvegicus* | 1 |
| testis | *B.taurus* | 5 | heart | *B.taurus* | 1 |
| testis | *S.scrofa* | 4 | heart | *S.scrofa* | 0 |
| testis | *G.gallus* | 6 | heart | *G.gallus* | 1 |
| **Tissue** | **species** | **#updated in v105** | **Tissue** | **species** | **#updated in v105** |
| brain | *H.sapiens* | 0 | kidney | *H.sapiens* | 1 |
| brain | *P.troglodytes* | 2 | kidney | *P.troglodytes* | 0 |
| brain | *M.mulatta* | 0 | kidney | *M.mulatta* | 1 |
| brain | *M.musculus* | 0 | kidney | *M.musculus* | 1 |
| brain | *R.norvegicus* | 2 | kidney | *R.norvegicus* | 1 |
| brain | *B.taurus* | 0 | kidney | *B.taurus* | 0 |
| brain | *S.scrofa* | 2 | kidney | *S.scrofa* | 0 |
| brain | *G.gallus* | 1 | kidney | *G.gallus* | 4 |

Table S16: TRGs and differential expression

| **species** | **tissue** | **nDE** | **nTRG** | **nDE+TRG** | **% TRG** | **% DE** |
|---|---|---|---|---|---|---|
| *H.sapiens* | brain | 1853 | 111 | 110 | 99.10 | 5.94 |
| *H.sapiens* | heart | 567 | 97 | 95 | 97.94 | 16.75 |
| *H.sapiens* | kidney | 1006 | 153 | 138 | 90.20 | 13.72 |
| *H.sapiens* | testis | 6578 | 278 | 278 | 100.00 | 4.23 |
| *P.troglodytes* | brain | 1219 | 135 | 130 | 96.30 | 10.66 |
| *P.troglodytes* | heart | 375 | 94 | 84 | 89.36 | 22.40 |
| *P.troglodytes* | kidney | 561 | 164 | 126 | 76.83 | 22.46 |
| *P.troglodytes* | testis | 2310 | 343 | 279 | 81.34 | 12.08 |
| *M.mulatta* | brain | 1395 | 162 | 157 | 96.91 | 11.25 |
| *M.mulatta* | heart | 542 | 110 | 106 | 96.36 | 19.56 |
| *M.mulatta* | kidney | 864 | 167 | 153 | 91.62 | 17.71 |
| *M.mulatta* | testis | 3346 | 291 | 290 | 99.66 | 8.67 |
| *M.musculus* | brain | 2357 | 138 | 137 | 99.28 | 5.81 |
| *M.musculus* | heart | 890 | 122 | 118 | 96.72 | 13.26 |
| *M.musculus* | kidney | 1224 | 191 | 176 | 92.15 | 14.38 |
| *M.musculus* | testis | 6207 | 316 | 273 | 86.39 | 4.40 |
| *R.norvegicus* | brain | 1571 | 82 | 79 | 96.34 | 5.03 |
| *R.norvegicus* | heart | 630 | 92 | 85 | 92.39 | 13.49 |
| *R.norvegicus* | kidney | 970 | 140 | 132 | 94.29 | 13.61 |
| *R.norvegicus* | testis | 3449 | 302 | 296 | 98.01 | 8.58 |
| *B.taurus* | brain | 1136 | 106 | 100 | 94.34 | 8.80 |
| *B.taurus* | heart | 694 | 98 | 91 | 92.86 | 13.11 |
| *B.taurus* | kidney | 752 | 156 | 142 | 91.03 | 18.88 |
| *B.taurus* | testis | 2200 | 336 | 312 | 92.86 | 14.18 |
| *S.scrofa* | brain | 1332 | 127 | 120 | 94.49 | 9.01 |
| *S.scrofa* | heart | 486 | 106 | 90 | 84.91 | 18.52 |
| *S.scrofa* | kidney | 730 | 157 | 127 | 80.89 | 17.40 |
| *S.scrofa* | testis | 2647 | 358 | 356 | 99.44 | 13.45 |
| *G.gallus* | brain | 1379 | 85 | 84 | 98.82 | 6.09 |
| *G.gallus* | heart | 543 | 75 | 74 | 98.67 | 13.63 |
| *G.gallus* | kidney | 777 | 155 | 140 | 90.32 | 18.02 |
| *G.gallus* | testis | 1840 | 253 | 235 | 92.89 | 12.77 |
| | | | | mean | 93.21 | 12.75 |

nDE: common DE identified for each tissue in a species
% TRG: % of TRGs that are also DE
% DE: % of non-TRG DE genes

Table S17: GO enrichment analysis for human DE genes with low gene loadings and DE + TRGs

| GO category | term | GO category | term |
|---|---|---|---|
| **testis** | | | |
| **DE with low coefficients** | | **DE + TRGs** | |
| GO:0007608 | sensory perception of smell | GO:0007283 | spermatogenesis |
| GO:0050896 | response to stimulus | GO:0030317 | flagellated sperm motility |
| GO:0050911 | detection of chemical stimulus involved in sensory perception of smell | GO:0051321 | meiotic cell cycle |
| GO:0070268 | cornification | GO:0030154 | cell differentiation |
| GO:0031424 | keratinization | GO:0007338 | single fertilization |
| **brain** | | | |
| **DE with low coefficients** | | **DE + TRGs** | |
| GO:0060041 | retina development in camera-type eye | GO:0007268 | chemical synaptic transmission |
| GO:0034765 | regulation of ion transmembrane transport | GO:0048168 | regulation of neuronal synaptic plasticity |
| GO:0050896 | response to stimulus | GO:0007612 | learning |
| GO:0001523 | retinoid metabolic process | GO:1903861 | positive regulation of dendrite extension |
| GO:0009416 | response to light stimulus | GO:0007269 | neurotransmitter secretion |
| **heart** | | | |
| **DE with low coefficients** | | **DE + TRGs** | |
| GO:0086069 | bundle of His cell to Purkinje myocyte communication | GO:0030240 | skeletal muscle thin filament assembly |
| GO:0033034 | positive regulation of myeloid cell apoptotic process | GO:0055003 | cardiac myofibril assembly |
| GO:0070543 | response to linoleic acid | GO:0055008 | cardiac muscle tissue morphogenesis |
| GO:0070994 | detection of oxidative stress | GO:0060048 | cardiac muscle contraction |
| GO:0090317 | negative regulation of intracellular protein transport | GO:0030049 | muscle filament sliding |
| **kidney** | | | |
| **DE with low coefficients** | | **DE + TRGs** | |
| GO:0001658 | branching involved in ureteric bud morphogenesis | GO:0006811 | ion transport |
| GO:0015730 | propanoate transport | GO:0055085 | transmembrane transport |
| GO:0015913 | short-chain fatty acid import | GO:0007588 | excretion |
| GO:0015802 | basic amino acid transport | GO:0006814 | sodium ion transport |
| GO:0003339 | regulation of mesenchymal to epithelial transition involved in metanephros morphogenesis | GO:0016338 | calcium-independent cell-cell adhesion via plasma membrane cell-adhesion molecules |

### References

Paal Skytt Andersen, Paula Louise Hedley, Stephen P Page, Petros Syrris, Johanna Catharina Moolman-Smook, William John McKenna, Perry Mark Elliott, and Michael Christiansen. A novel myosin essential light chain mutation causes hypertrophic cardiomyopathy with late onset and low expressivity. *Biochemistry Research International*, 2012, 2012.

Silvia Arber, John J Hunter, John Ross Jr, Minoru Hongo, Gilles Sansig, Jacques Borg, Jean-Claude Perriard, Kenneth R Chien, and Pico Caroni. Mlp-deficient mice exhibit a disruption of cardiac cytoarchitectural organization, dilated cardiomyopathy, and heart failure. *Cell*, 88(3):393–403, 1997.

Ferial Aslani, Mohammad Hossein Modarresi, Haleh Soltanghoraee, Mohammad Mehdi Akhondi, Ashraf Shabani, Niknam Lakpour, and Mohammad Reza Sadeghi. Seminal molecular markers as a non-invasive diagnostic tool for the evaluation of spermatogenesis in non-obstructive azoospermia. *Systems Biology in Reproductive Medicine*, 57(4):190–196, 2011.

Kate Baker, Sarah L Gordon, Holly Melland, Fabian Bumbak, Daniel J Scott, Tess J Jiang, David Owen, Bradley J Turner, Stewart G Boyd, Mari Rossi, et al. Syt1-associated neurodevelopmental disorder: a case series. *Brain*, 141(9):2576–2591, 2018.

Detlev Boison, H Bussow, Donatella D'Urso, HW Muller, and Wilhelm Stoffel. Adhesive properties of proteolipid protein are responsible for the compaction of cns myelin sheaths. *Journal of Neuroscience*, 15(8):5502–5513, 1995.

Karin Borges and Raymond Dingledine. Ampa receptors: molecular and functional diversity. *Progress in Brain Research*, 116:153–170, 1998.

Alice CN Brown, Dalila Hallouane, William J Mawby, Fiona E Karet, Moin A Saleem, Alexander J Howie, and Ashley M Toye. RhCG is the major putative ammonia transporter expressed in the human kidney, and RhBG is not expressed at detectable levels. *American Journal of Physiology-Renal Physiology*, 296(6):F1279–F1290, 2009.

Amanda Milgram Burkhoff, David L Linemeyer, and John A Salon. Distribution of a novel hypothalamic neuropeptide Y receptor gene and its absence in rat. *Molecular Brain Research*, 53(1-2):311–316, 1998.

Lauren M Byrne, Filipe B Rodrigues, Kaj Blennow, Alexandra Durr, Blair R Leavitt, Raymund AC Roos, Rachael I Scahill, Sarah J Tabrizi, Henrik Zetterberg, Douglas Langbehn, et al. Neurofilament light protein in blood as a potential biomarker of neurodegeneration in Huntington's disease: a retrospective cohort analysis. *The Lancet Neurology*, 16(8):601–609, 2017.

Harsha D Devalla, Roselle Gélinas, Elhadi H Aburawi, Abdelaziz Beqqali, Philippe Goyette, Christian Freund, Marie-A Chaix, Rafik Tadros, Hui Jiang, Antony Le Béchec, et al. TECRL, a new life-threatening inherited arrhythmia gene associated with overlapping clinical features of both LQTS and CPVT. *EMBO Molecular Medicine*, 8(12):1390–1408, 2016.

Dijana Djureinovic, Linn Fagerberg, Björn Hallström, A Danielsson, C Lindskog, Mathias Uhlén, and Fredrik Pontén. The human testis-specific proteome defined by transcriptomics and antibody-based profiling. *Molecular Human Reproduction*, 20(6):476–488, 2014.

Lihua Dong, Stefan Pietsch, Zenglai Tan, Birgit Perner, Ralph Sierig, Dagmar Kruspe, Marco Groth, Ralph Witzgall, Hermann-Josef Gröne, Matthias Platzer, et al. Integration of cistromic and transcriptomic analyses identifies nphs2, mafb, and magi2 as wilms' tumor 1 target genes in podocyte differentiation and maintenance. *Journal of the American Society of Nephrology*, 26(9):2118–2128, 2015.

Linn Fagerberg, Björn M Hallström, Per Oksvold, Caroline Kampf, Dijana Djureinovic, Jacob Odeberg, Masato Habuka, Simin Tahmasebpoor, Angelika Danielsson, Karolina Edlund, et al. Analysis of the human tissue-specific expression by genome-wide integration of transcriptomics and antibody-based proteomics. *Molecular & Cellular Proteomics*, 13(2):397–406, 2014.

Michela Faggioni and Björn C Knollmann. Calsequestrin 2 and arrhythmias. *American Journal of Physiology-Heart and Circulatory Physiology*, 302(6):H1250–H1260, 2012.

Amy Fearn, Benjamin Allison, Sarah J Rice, Noel Edwards, Jan Halbritter, Soline Bourgeois, Eva M Pastor-Arroyo, Friedhelm Hildebrandt, Velibor Tasic, Carsten A Wagner, et al. Clinical, biochemical, and pathophysiological analysis of slc 34a1 mutations. *Physiological Reports*, 6(12):e13715, 2018.

Derk Frank, Ashraf Yusuf Rangrez, Corinna Friedrich, Sven Dittmann, Birgit Stallmeyer, Pankaj Yadav, Alexander Bernt, Ellen Schulze-Bahr, Ankush Borlepawar, Wolfram-Hubertus Zimmermann, et al. Cardiac $\alpha$-actin (ACTC1) gene mutation causes atrial-septal defects associated with late-onset dilated cardiomyopathy. *Circulation: Genomic and Precision Medicine*, 12(8):e002491, 2019.

Ke Gong, Ting Xie, Yong Luo, Hui Guo, Jinlan Chen, Zhiping Tan, Yifeng Yang, and Li Xie. Comprehensive analysis of lncRNA biomarkers in kidney renal clear cell carcinoma by lncRNA-mediated ceRNA network. *PLOS ONE*, 16(6):e0252452, 2021.

Pei Han, Wei Li, Chiou-Hong Lin, Jin Yang, Ching Shang, Sylvia T Nurnberg, Kevin Kai Jin, Weihong Xu, Chieh-Yu Lin, Chien-Jung Lin, et al. A long noncoding RNA protects the heart from pathological hypertrophy. *Nature*, 514(7520):102–106, 2014.

Peter E Hart, Janel N Glantz, James D Orth, Gregory M Poynter, and Jeffrey L Salisbury. Testis-specific murine centrin, cetn1: genomic characterization and evidence for retroposition of a gene encoding a centrosome protein. *Genomics*, 60(2):111–120, 1999.

Jiayuan He, Yixue Xue, Qingyuan Wang, Xinxin Zhou, Libo Liu, Tianyuan Zhang, Chao Shang, Jun Ma, and Teng Ma. Long non-coding RNA MIAT regulates blood tumor barrier permeability by functioning as a competing endogenous RNA. *Cell Death & Disease*, 11(10):1–18, 2020.

John C Herr, David Thomas, Leigh Ann Bush, Scott Coonrod, Vrinda Khole, Stuart S Howards, and Charles J Flickinger. Sperm mitochondria-associated cysteine-rich protein (SMCP) is an autoantigen in Lewis rats. *Biology of Reproduction*, 61(2):428–435, 1999.

HPA. The Human Protein Atlas, v21.0.

Takuma Iguchi, Noriyo Niino, Satoshi Tamai, Ken Sakurai, and Kazuhiko Mori. Comprehensive analysis of circulating microRNA specific to the liver, heart, and skeletal muscle of cynomolgus monkeys. *International Journal of Toxicology*, 36(3):220–228, 2017.

Huiming Jiang, Haibin Chen, Pei Wan, Shengda Song, and Nanhui Chen. Downregulation of enhancer RNA EMX2OS is associated with poor prognosis in kidney renal clear cell carcinoma. *Aging (Albany NY)*, 12(24):25865, 2020.

Yuzuru Kato and Masami Nozaki. Distinct DNA Methylation Dynamics of Spermatogenic Cell-Specific Intronless Genes Is Associated with CpG Content. *PLOS ONE*, 7(8):1–10, 08 2012.

Yuhei Kirita, Haojia Wu, Kohei Uchimura, Parker C Wilson, and Benjamin D Humphreys. Cell profiling of mouse acute kidney injury reveals conserved cellular responses to injury. *Proceedings of the National Academy of Sciences*, 117(27):15874–15883, 2020.

Keiko Kiyonaga-Endou, Manabu Oshima, Kazuya Sugimoto, Mervyn Thomas, Shigeru Taketani, and Masasuke Araki. Localization of neurensin1 in cerebellar purkinje cells of the developing chick and its possible function in dendrite formation. *Brain Research*, 1635:113–120, 2016.

Martin Konrad, Tom Nijenhuis, Gema Ariceta, Aurelia Bertholet-Thomas, Lorenzo A Calo, Giovambattista Capasso, Francesco Emma, Karl P Schlingmann, Mandeep Singh, Francesco Trepiccione, et al. Diagnosis and management of bartter syndrome: executive summary of the consensus and recommendations from the european rare kidney disease reference network working group for tubular disorders. *Kidney International*, 99(2):324–335, 2021.

Monica X Li and Peter M Hwang. Structure and function of cardiac troponin C (TNNC1): Implications for heart failure, cardiomyopathies, and troponin modulating drugs. *Gene*, 571(2):153–166, 2015.

Yang Li, Bo Lin, and Lei Yang. Comparative transcriptomic analysis of multiple cardiovascular fates from embryonic stem cells predicts novel regulators in human cardiogenesis. *Scientific Reports*, 5(1):1–16, 2015.

Siyuan John Liu, Tomasz J Nowakowski, Alex A Pollen, Jan H Lui, Max A Horlbeck, Frank J Attenello, Daniel He, Jonathan S Weissman, Arnold R Kriegstein, Aaron A Diaz, et al. Single-cell analysis of long non-coding RNAs in the developing human neocortex. *Genome Biology*, 17(1):1–17, 2016.

Wensheng Liu, Yinan Zhao, Xiaohua Liu, Xiaoya Zhang, Jiancheng Ding, Yang Li, Yingpu Tian, Haibin Wang, Wen Liu, and Zhongxian Lu. A Novel Meiosis-Related lncRNA, Rbakdn, Contributes to Spermatogenesis by Stabilizing Ptbp2. *Frontiers in Genetics*, page 1963, 2021.

Y Liu, E McKenna, DJ Figueroa, R Blevins, CP Austin, PB Bennett, and R Swanson. The human inward rectifier K+ channel subunit kir5. 1 (KCNJ16) maps to chromosome 17q25 and is expressed in kidney and pancreas. *Cytogenetic and Genome Research*, 90(1-2):60–63, 2000.

Yanyan Liu, Min Jiang, Chao Li, Ping Yang, Huaqin Sun, Dachang Tao, Sizhong Zhang, and Yongxin Ma. Human t-complex protein 11 (TCP11), a testis-specific gene product, is a potential determinant of the sperm morphology. *The Tohoku Journal of Experimental Medicine*, 224(2):111–117, 2011.

P Mali, A Kaipia, M Kangasniemi, J Toppari, M Sandberg, NB Hecht, and M Parvinen. Stage-specific expression of nucleoprotein mRNAs during rat and mouse spermiogenesis. *Reproduction, Fertility and Development*, 1(4):369–382, 1989.

Nicolas Markadieu and Eric Delpire. Physiology and pathophysiology of SLC12A1/2 transporters. *Pflügers Archiv-European Journal of Physiology*, 466(1):91–105, 2014.

Giulia Mearini, Doreen Stimpel, Birgit Geertz, Florian Weinberger, Elisabeth Krämer, Saskia Schlossarek, Julia Mourot-Filiatre, Andrea Stoehr, Alexander Dutsch, Paul JM Wijnker, et al. Mybpc3 gene therapy for neonatal cardiomyopathy enables long-term disease prevention in mice. *Nature Communications*, 5(1):1–10, 2014.

Jiao Meng, Limin Li, Yue Zhao, Zhen Zhou, Mingchao Zhang, Donghai Li, Chen-Yu Zhang, Ke Zen, and Zhihong Liu. MicroRNA-196a/b mitigate renal fibrosis by targeting TGF-$\beta$ receptor 2. *Journal of the American Society of Nephrology*, 27(10):3006–3021, 2016.

Domenico Milardi, Giuseppe Grande, Federica Vincenzoni, Francesco Pierconti, and Alfredo Pontecorvi. Proteomics for the identification of biomarkers in testicular cancer–review. *Frontiers in Endocrinology*, 10:462, 2019.

Maolin Mu, Wanxiang Niu, Xiaoming Zhang, Shanshan Hu, and Chaoshi Niu. LncRNA BCYRN1 inhibits glioma tumorigenesis by competitively binding with miR-619-5p to regulate CUEDC2 expression and the PTEN/AKT/p21 pathway. *Oncogene*, 39(45):6879–6892, 2020.

Ashley D Nemes, Katayoun Ayasoufi, Zhong Ying, Qi-Gang Zhou, Hoonkyo Suh, and Imad M Najm. Growth associated protein 43 (GAP-43) as a novel target for the diagnosis, treatment and prevention of Epileptogenesis. *Scientific Reports*, 7(1):1–13, 2017.

Adriana Osio, Lily Tan, Suet N Chen, Raffaella Lombardi, Sherif F Nagueh, Sanjay Shete, Robert Roberts, James T Willerson, and Ali J Marian. Myozenin 2 is a novel gene for human hypertrophic cardiomyopathy. *Circulation Research*, 100(6):766–768, 2007.

Konrad Platzer and Johannes R Lemke. Grin1-related neurodevelopmental disorder, 2019.

Murugavel Ponnusamy, Fang Liu, Yu-Hui Zhang, Rui-Bei Li, Mei Zhai, Fei Liu, Lu-Yu Zhou, Cui-Yun Liu, Kao-Wen Yan, Yan-Han Dong, et al. Long noncoding RNA CPR (cardiomyocyte proliferation regulator) regulates cardiomyocyte proliferation and cardiac repair. *Circulation*, 139(23):2668–2684, 2019.

Seungil Ro, Chanjae Park, Kenton M Sanders, John R McCarrey, and Wei Yan. Cloning and expression profiling of testis-expressed microRNAs. *Developmental Biology*, 311(2):592–602, 2007.

Irene Sanchez-Martin, Pedro Magalhães, Parisa Ranjzad, Ahmed Fatmi, Fabrice Richard, Thien Phong Vu Manh, Andrew J Saurin, Guylène Feuillet, Colette Denis, Adrian S Woolf, et al. Haploinsufficiency of the mouse Tshz3 gene leads to kidney defects. *Human Molecular Genetics*, 2021.

Rikako Sanuki, Akishi Onishi, Chieko Koike, Rieko Muramatsu, Satoshi Watanabe, Yuki Muranishi, Shoichi Irie, Shinji Uneo, Toshiyuki Koyasu, Ryosuke Matsui, et al. miR-124a is required for hippocampal axogenesis and retinal cone survival through Lhx2 suppression. *Nature Neuroscience*, 14(9):1125–1134, 2011.

Farah Sheikh, Robert C Lyon, and Ju Chen. Functions of myosin light chain-2 (myl2) in cardiac muscle and disease. *Gene*, 569(1):14–20, 2015.

Stephanie L Skala, Xiaoming Wang, Yuping Zhang, Rahul Mannan, Lisha Wang, Sathiya P Narayanan, Pankaj Vats, Fengyun Su, Jin Chen, Xuhong Cao, et al. Next-generation RNA sequencing–based biomarker characterization of chromophobe renal cell carcinoma and related oncocytic neoplasms. *European Urology*, 78(1):63–74, 2020.

EH Stover, KJ Borthwick, C Bavalia, N Eady, DM Fritz, N Rungroj, ABS Giersch, CC Morton, PR Axon, I Akil, et al. Novel ATP6V1B1 and ATP6V0A4 mutations in autosomal recessive distal renal tubular acidosis with new evidence for hearing loss. *Journal of Medical Genetics*, 39(11):796–803, 2002.

Wen Su, Rong Cao, Xiao-yan Zhang, and Youfei Guan. Aquaporins in the kidney: physiology and pathophysiology. *American Journal of Physiology-Renal Physiology*, 318(1):F193–F203, 2020.

Lawrence CR Tafoya, Manuel Mameli, Teiko Miyashita, John F Guzowski, C Fernando Valenzuela, and Michael C Wilson. Expression and function of snap-25 as a universal snare component in gabaergic neurons. *Journal of Neuroscience*, 26(30):7826–7838, 2006.

Men C Tan, Jocelyn Widagdo, Yu Q Chau, Tianyi Zhu, Justin J-L Wong, Allen Cheung, and Victor Anggono. The activity-induced long non-coding RNA Meg3 modulates AMPA receptor surface expression in primary cortical neurons. *Frontiers in Cellular Neuroscience*, 11:124, 2017.

Maciej Tomaszewski, James Eales, Matthew Denniff, Stephen Myers, Guat Siew Chew, Christopher P Nelson, Paraskevi Christofidou, Aishwarya Desai, Cara Büsst, Lukasz Wojnar, et al. Renal mechanisms of association between fibroblast growth factor 1 and blood pressure. *Journal of the American Society of Nephrology*, 26(12):3151–3160, 2015.

Mathias Uhlén, Linn Fagerberg, Björn M Hallström, Cecilia Lindskog, Per Oksvold, Adil Mardinoglu, Åsa Sivertsson, Caroline Kampf, Evelina Sjöstedt, Anna Asplund, et al. Tissue-based map of the human proteome. *Science*, 347(6220):1260419, 2015.

Pingzhang Wang, Bo Sun, Dongxia Hao, Xiujun Zhang, Taiping Shi, and Dalong Ma. Human TMEM174 that is highly expressed in kidney tissue activates AP-1 and promotes cell proliferation. *Biochemical and Biophysical Research Communications*, 394(4):993–999, 2010.

Junco S Warren, Christopher M Tracy, Mickey R Miller, Aman Makaju, Marta W Szulik, Shin-ichi Oka, Tatiana N Yuzyuk, James E Cox, Anil Kumar, Bucky K Lozier, et al. Histone methyltransferase smyd1 regulates mitochondrial energetics in the heart. *Proceedings of the National Academy of Sciences*, 115(33):E7871–E7880, 2018.

Susan M Wykes, James E Nelson, Daniel W Visscher, Daniel Djakiew, and Stephen A Krawetz. Coordinate expression of the PRM1, PRM2, and TNP2 multigene locus in human testis. *DNA and Cell Biology*, 14(2):155–161, 1995.

Yun Xie, Jiahui Yao, Xinzong Zhang, Jun Chen, Yong Gao, Chi Zhang, Haicheng Chen, Zelin Wang, Zhiying Zhao, Wenqiu Chen, et al. A panel of extracellular vesicle long noncoding RNAs in seminal plasma for predicting testicular spermatozoa in nonobstructive azoospermia patients. *Human Reproduction*, 35(11):2413–2427, 2020.

Wei Yan, Lang Ma, Kathleen H Burns, and Martin M Matzuk. Haploinsufficiency of kelch-like protein homolog 10 causes infertility in male mice. *Proceedings of the National Academy of Sciences*, 101(20):7793–7798, 2004.

Kefei Yang, Andreas Meinhardt, Bing Zhang, Pawel Grzmil, Ibrahim M Adham, and Sigrid Hoyer-Fender. The small heat shock protein ODF1/HSPB10 is essential for tight linkage of sperm head to tail and male fertility in mice. *Molecular and Cellular Biology*, 32(1):216–225, 2012.

Zhihui Yang and Kevin KW Wang. Glial fibrillary acidic protein: from intermediate filament assembly and gliosis to neurobiomarker. *Trends in Neurosciences*, 38(6):364–374, 2015.

Shuiqiao Yuan, Weibing Qin, Connor R Riordan, Hayden Mcswiggin, Huili Zheng, and Wei Yan. Ubqln3, a testis-specific gene, is dispensable for embryonic development and spermatogenesis in mice. *Molecular Reproduction and Development*, 82(4):266, 2015.

Scott A Yuzwa, Michael J Borrett, Brendan T Innes, Anastassia Voronova, Troy Ketela, David R Kaplan, Gary D Bader, and Freda D Miller. Developmental emergence of adult neural stem cells as revealed by single-cell transcriptional profiling. *Cell Reports*, 21(13):3970–3986, 2017.

Lu Zhang, Antonio Salgado-Somoza, Melanie Vausort, Przemyslaw Leszek, Yvan Devaux, et al. A heart-enriched antisense long non-coding RNA regulates the balance between cardiac and skeletal muscle triadin. *Biochimica et Biophysica Acta (BBA)-Molecular Cell Research*, 1865(2):247–258, 2018.
