## Additional File 2 for "Orthogonal Shared Basis Factorization: Cross-species gene expression analysis using a common expression subspace"

### Additional File 2: Supplementary Methods: Orthogonal Shared Basis Factorization

Amal Thomas

Quantitative and Computational Biology, University of Southern California, Los Angeles, CA, 90007 USA

In cross-species gene expression analysis using OSBF, we define our shared factor as a subspace representing similar inter-sample correlation relationships. We also require the independence of the columns of the species-specific factors as it helps to differentiate the contribution of genes/features across different dimensions of the shared factor. The diagonal matrices represent the amount of information represented by individual dimensions. Given these conditions, an exact factorization is not always possible. This introduces errors in individual factorizations, and we want to minimize the total factorization error.

We estimate the shared factor $V$ from the expected inter-sample correlation matrix across species. The different dimensions of $V$ represent the correlation relationship between the samples independent of the species. An orthonormal basis of this shared subspace is used as the axes of the common space. Given $V$, we compute initial estimates of $U_i$ and $\Delta_i$ by solving the linear systems $D_iV = U_i\Delta_i$. This gives us an exact solution for all $k$ factorizations. The initial estimates of $U_i$'s do not have independent columns. Orthonormalizing $U_i$'s introduce errors, and depending upon the $D_i$ matrices, an exact factorization is not possible. Our goal in the optimization is that, given our initial estimate of meaningful common expression subspace, we want to find the closest configuration of $U_i$, and $\Delta_i$, that minimize the total factorization error.

In the following sections, we will discuss how to achieve the best configuration of $V$, $U_i$, and $\Delta_i$ that minimizes the total reconstruction error $\sum_{i=1}^{k} \|\epsilon_i\|_F^2$. First, we will discuss our objective function, the constraints, and the general strategy to minimize the error.

#### 1 Minimizing error in orthogonal shared basis factorization

Here we provide details on minimizing the decomposition error of the OSBF method. In OSBF, we want to solve the following optimization problem:

$$\begin{aligned}
\text{minimize:} \quad & \textstyle\sum_{i=1}^{k} \|D_i - U_i\Delta_iV^T\|_F^2, \\
\text{subject to:} \quad & U_i^TU_i = I, \ \text{for } 1 \le i \le k, \\
& V^TV = VV^T = I, \\
& \Delta_i = \operatorname{diag}(\delta_{i,1}, \ldots, \delta_{i,n}), \\
& \delta_{i,j} \ge 0, \ \text{for } 1 \le i \le k.
\end{aligned}$$

The objective function can be reformulated as

$$\textstyle\sum_{i=1}^{k} \operatorname{tr}(D_i^TD_i) - 2\operatorname{tr}\left(D_i^TU_i\Delta_iV^T\right) + \operatorname{tr}(\Delta_i^2)$$

Let $\Phi$, $\Psi_i$, and $\Theta_i$ be matrices with Lagrange multipliers for constraints $V^TV - I = 0$, $U_i^TU_i - I = 0$ and $\delta_{i,j} \geq 0$. The Lagrange $\mathcal{L}$ is

$$\mathcal{L}(U, \Delta, V) = \sum_{i=1}^{k} \mathrm{tr}(D_i^TD_i - 2D_i^TU_i\Delta_iV^T + \Delta_i^2) + \mathrm{tr}(\Phi(V^TV - I)) + \mathrm{tr}(\Psi_i(U_i^TU_i - I)) + \mathrm{tr}(\Theta_i\Delta_i).$$

Solving for an individual $U_i$ using the partial derivative of $\mathcal{L}$ with respect to $U_i$ gives

$$\frac{\partial}{\partial U_i}\mathcal{L}(U, \Delta, V) = -2D_iV\Delta_i + U_i(\Psi_i^T + \Psi_i)$$
$$U_i = D_iV\Delta_i((D_iV\Delta_i)^T(D_iV\Delta_i))^{-1/2}. \tag{1}$$

Solving $V$ using the partial derivative of $\mathcal{L}$ with respect to $V$ gives

$$\frac{\partial}{\partial V}\mathcal{L}(U, \Delta, V) = \sum_{i=1}^{k} -2D_i^TU_i\Delta_i + V(\Phi^T + \Phi)$$
$$V = \sum_{i=1}^{k} D_i^TU_i\Delta_i((\sum_{i=1}^{k} D_i^TU_i\Delta_i)^T(\sum_{i=1}^{k} D_i^TU_i\Delta_i))^{-1/2}. \tag{2}$$

Solving for an individual $\Delta_i$ using the partial derivative of $\mathcal{L}$ with respect to $\Delta_i$ gives

$$\frac{\partial}{\partial \Delta_i}\mathcal{L}(U, \Delta, V) = -2U_i^TD_iV + 2\Delta_i + \Theta_i$$
$$\Delta_i = U_i^TD_iV. \tag{3}$$

#### 2 Method derivation

The section 1 shows that given the other two factors, we can find the optimal third factor: left basis matrices (equation 1), the right basis matrix (equation 2), and the diagonal matrices (equation 3). Now, given any two factors we will discuss how to efficiently update the third factor and minimize the total factorization error. In practice, we do not compute the square root inverse to update $U_i$ and $V$. We can make the optimal update of our matrix in each iteration based on the singular value decomposition (SVD) of other matrices. The first two results below show that, given either the left or right basis matrix, we can find the other matrix (optimal) with the required properties.

**Proposition 1.** Consider matrices $A \in \mathbb{R}^{m\times n}$ with rank $n$ and $Q \in \mathbb{R}^{n\times n}$. If $X\Sigma Y^T$ is the SVD of $AQ^T$, then among matrices $B \in \mathbb{R}^{m\times n}$ with orthonormal columns, $\|A - BQ\|_F$ is minimized when $B = XY^T$.

*Proof.* The norm and its square have the same minimum value, so we consider the minimum of

$$\|A - BQ\|_F^2 = \mathrm{tr}(A^TA) + \mathrm{tr}(Q^TQ) - 2\,\mathrm{tr}(AQ^TB^T),$$

which coincides with the maximum value of $\mathrm{tr}(AQ^TB^T)$. With $X\Sigma Y^T$ the SVD of $AQ^T$, define $Z = Y^TB^TX$. Since $B^TB = I$, we have $ZZ^T = I$. We can bound the maximum value of $\mathrm{tr}(AQ^TB^T)$ as follows:

$$\mathrm{tr}(AQ^TB^T) = \mathrm{tr}(X\Sigma Y^TB^T) = \mathrm{tr}(Z\Sigma) = \textstyle\sum_{i=1}^{n} Z_{ii}\sigma_i \leq \sum_{i=1}^{n} \sigma_i,$$

and this bound is attained when $Z = I$, and thus $B = XY^T$. □

In OSBF updates, we use the proposition 1 to find the optimal left basis matrices. In proposition 1, let us substitute $A = D_i, B = U_i$, and $Q = \Delta_i V^T$. Now we have $AQ^T = D_i V \Delta_i = X\Sigma Y^T$. The $U_i$ with orthonormal columns is given by $XY^T$. The same can also be obtained by substituting $D_i V \Delta_i = X\Sigma Y^T$ in equation 1:

$$U_i = D_i V \Delta_i ((D_i V \Delta_i)^T (D_i V \Delta_i))^{-1/2}$$
$$U_i = X\Sigma Y^T ((X\Sigma Y^T)^T (X\Sigma Y^T))^{-1/2} = XY^T.$$

**Proposition 2.** Consider matrices $A \in \mathbb{R}^{m\times n}$ with rank $n$ and $B \in \mathbb{R}^{m\times n}$. If $X\Sigma Y^T$ is the SVD of $A^T B$, then among orthogonal matrices $Q \in \mathbb{R}^{n\times n}$, $\|A - BQ^T\|_F$ is minimized when $Q = XY^T$.

*Proof.* The norm and its square have the same minimum value, so we consider the minimum of

$$\|A - BQ^T\|_F^2 = \mathrm{tr}(A^T A) + \mathrm{tr}(B^T B) - 2\,\mathrm{tr}(A^T B Q^T),$$

which coincides with the maximum value of $\mathrm{tr}(A^T B Q^T)$. With $X\Sigma Y^T$ the SVD of $A^T B$, define orthogonal matrix $Z = Y^T Q^T X$. We bound the maximum of $\mathrm{tr}(A^T B Q^T)$ as

$$\mathrm{tr}(A^T B Q^T) = \mathrm{tr}(X\Sigma Y^T Q^T) = \mathrm{tr}(Z\Sigma) = \textstyle\sum_{i=1}^n Z_{ii}\sigma_i \le \sum_{i=1}^n \sigma_i,$$

and this bound is attained when $Z = I$ and thus $Q = XY^T$. □

Using proposition 2, we can find the orthogonal right matrix for each factorization ($\Omega_i$) involved in the OSBF. In proposition 2, let us substitute $A = D_i, B = U_i\Delta_i$, and $Q = \Omega_i$. Now we have $A^T B = D_i^T U_i \Delta_i = X\Sigma Y^T$ and orthogonal $\Omega_i$ is given by $XY^T$. The computed matrices using proposition 2 are not shared and also does not guarantee that the total factorization error involved $\sum_{i=1}^k \|D_i^T U_i \Delta_i - \Omega_i\|_F^2$ is minimized. We will now further expand the proposition 2 to find the optimal shared right basis matrix.

**Proposition 3.** For a set of $n$ matrices $A_1, A_2, \ldots, A_n$, where $A_i \in \mathbb{R}^{k\times k}$, the orthogonal matrix $Q \in \mathbb{R}^{k\times k}$ minimizing $\sum_{i=1}^n \|A_i - Q\|_F^2$ is given by $Q = XY^T$, where SVD of $\sum_{i=1}^n A_i = X\Sigma Y^T$.

*Proof.* We seek to minimize

$$\textstyle\sum_{i=1}^n \|A_i - Q\|_F^2 \text{ subject to } Q^T Q = I.$$

Defining $A = \sum_{i=1}^n A_i$, this objective function can be rewritten

$$\mathrm{tr}(\textstyle\sum_{i=1}^n A_i^T A_i) - 2\,\mathrm{tr}(A^T Q) + n\,\mathrm{tr}(Q^T Q).$$

Let $\Phi$ be a matrix of Lagrange multipliers for $Q^T Q - I = 0$. The Lagrangian $\mathcal{L}$ is then

$$\mathcal{L}(Q, \Phi) = \mathrm{tr}(\textstyle\sum_{i=1}^n A_i^T A_i) - 2\,\mathrm{tr}(A^T Q) + \mathrm{tr}(Q^T Q) + \mathrm{tr}(\Phi(Q^T Q - I)).$$

Since $Q$ is orthogonal, the partial derivatives of $\mathcal{L}(Q, \Phi)$ with respect to $Q$ are as follows:

$$\frac{\partial \mathcal{L}(Q, \Phi)}{\partial Q} = -2A + Q(\Phi^T + \Phi).$$

Setting this equation to 0 yields:

$$Q = A\left(\frac{\Phi^T + \Phi}{2}\right)^{-1} \text{ and } A = Q\left(\frac{\Phi^T + \Phi}{2}\right).$$

Again because $Q$ is orthogonal,

$$A^T A = \left(\frac{\Phi^T + \Phi}{2}\right)^2 \quad \text{and} \quad \left(\frac{\Phi^T + \Phi}{2}\right) = (A^T A)^{1/2},$$

which implies $Q = A(A^T A)^{-1/2}$. Since $X\Sigma Y^T$ is the SVD of $A$ we conclude

$$Q = X\Sigma Y^T (Y\Sigma^2 Y^T)^{-1/2} = XY^T.$$

□

In OSBF, $V$ is shared across the $k$ factorizations. To find the closest shared $V$, we will use proposition 3. In proposition 3, let us substitute $A_i = D_i^T U_i \Delta_i$ and $Q = V$. Now we have $\sum_{i=1}^n A_i = \sum_{i=1}^k D_i^T U_i \Delta_i = X\Sigma Y^T$ and the optimal shared right basis in OSBF is given by $V = XY^T$.

#### 3 Iterative algorithm to minimize factorization error

Based on the equations 1-3 and propositions 1-3, we devised an iterative algorithm to minimize the total decomposition error. In each iteration, we make the optimal update, and since the objective function value is bounded below by zero; our procedure will converge to a stable local optimum. Steps for the iterative updates of the factorization are shown below:

**Algorithm**

1: Input: $D_i, i = 1, \ldots, k$
2: Output: $U_i, \Delta_i$, and $V$, where $U_i^T U_i = I$ and $V^T V = VV^T = I$
3: Initialize $U_i^j$, $\Delta_i^j$, and $V^j$. Initialize $\epsilon = \infty$. Let $\|\epsilon^{j,j,j}\|_F = \sum_{i=1}^k \|D_i - U_i^j \Delta_i^j V^{j^T}\|_F$
4: **Update $\mathbf{U_i}$**:
   $U_i^{j+1} \leftarrow ZY^T$, where SVD of $D_i V^j \Delta_i^j = Z\Sigma Y^T$. Compute $\|\epsilon^{j+1,j,j}\|_F$
   If $\|\epsilon^{j+1,j,j}\|_F < \epsilon$:
      $U_i \leftarrow U_i^{j+1}$
      $\epsilon \leftarrow \|\epsilon^{j+1,j,j}\|_F$
5: **Update $\mathbf{\Delta_i}$**:
   $\Delta_i^{j+1} \leftarrow \mathrm{diag}((U_i)^T D_i V^j)$. Compute $\|\epsilon^{j,j+1,j}\|_F$
   If $\|\epsilon^{j,j+1,j}\|_F < \epsilon$:
      $\Delta_i \leftarrow \Delta_i^{j+1}$
      $\epsilon \leftarrow \|\epsilon^{j,j+1,j}\|_F$
6: **Update $\mathbf{V}$**:
   $V^{j+1} \leftarrow MQ^T$, where SVD of $\sum_{i=1}^k D_i^T U_i \Delta_i = M\Phi Q^T$. Compute $\|\epsilon^{j,j,j+1}\|_F$
   If $\|\epsilon^{j,j,j+1}\|_F < \epsilon$:
      $V \leftarrow V^{j+1}$
      $\epsilon \leftarrow \|\epsilon^{j,j,j+1}\|_F$
7: Repeat steps 4-6 until convergence.

For a gene expression profile $Y_i$ from species $i$, minimizing the factorization error provides us the configuration of $U_i$'s and $\Delta_i$'s such that when we project a gene expression profile using $Y_i^T U_i \Delta_i^{-1}$, it lies close to the estimated common expression subspace $V$. In addition, our update strategy reduces the initial factorization error by maintaining the properties of the factors. For the common expression subspace, let the initial estimate of the common expression subspace using the eigenvalue decomposition of the expected $E(R)$ matrix be $V$. Let the optimized shared space be $\tilde{V}$. We have

$$R_\theta V = \tilde{V}, \text{ where}$$

$R_\theta \in \mathbb{R}^{n \times n}$ is a matrix. In OSBF, based on proposition 2 and 3, we express all $V$'s including our initial estimate as the product of a $XY^T$, where $X$ is a left singular matrix and $Y$ is a right singular matrix. For a given $V$, this is in fact the SVD with the diagonal matrix storing singular values being the identity matrix. So the singular values are 1 and hence all the eigenvalues are 1. Thus the determinant of $V$'s and $\tilde{V}$ is 1. This makes $R_\theta$ a pure rotational matrix. We can show that optimized space $\tilde{V}$ represents the expected inter-tissue correlation relationship in this rotated space. For instance, if we apply the same rotation to our standardized gene expression matrices, we have $XR_\theta^T$. Now we have,

$$\begin{aligned}
\mathrm{E}(R) &= \frac{1}{k}\sum_{i=1}^{k}\frac{1}{m_i}(X_i R_\theta^T)^T X_i R_\theta^T \\
&= \frac{1}{k}\sum_{i=1}^{k}\frac{1}{m_i} R_\theta X_i^T X_i R_\theta^T \\
&= R_\theta \frac{1}{k}\sum_{i=1}^{k}\frac{1}{m_i} X_i^T X_i R_\theta^T \\
&= (R_\theta V)\Lambda(R_\theta V)^T \\
&= \tilde{V}\Lambda\tilde{V}
\end{aligned}$$

In summary, the optimization strategy can be summarized as a procedure where we apply rotation to the initial estimate of common expression subspace and finding the corresponding optimal $U_i$'s and $\Delta_i$'s. We also show that the optimized $V$ has similar properties to that of the initial estimate and optimized $U_i$'s and $\Delta_i$'s are easily interpretable using cross-species gene expression analysis (see R package vignettes).

#### 3.1 Optimal learning rate in each update

In gradient descent, we have the general update rule:

$$\theta_{t+1} = \theta_t - \eta_t \nabla \mathcal{F}(\theta_t),$$

where $t$ is the iteration counter, $\eta$ is the learning rate at iteration $t$, and $\nabla \mathcal{F}$ is the gradient of the objective function. In our algorithm, the gradient of the objective function with respect to $U$ is $-2DV\Delta$. So we have the update rule as

$$U_{t+1} = U_t + \eta_t{}' DV\Delta.$$

By setting $\eta_t{}' = \frac{XY^T - U_t}{DV\Delta}$, we have $U_{t+1} = DV\Delta$, where SVD of $DV\Delta = X\Sigma Y^T$. From proposition 1, this is the optimal value for $U$, for a given $D_i$, $V$, and $\Delta_i$. The learning rate determines the size of the steps. If the value of $U_t$ is very different from $XY^T$, $\eta$ will be high, and we will take larger steps. As it becomes closer to $XY^T$, the learning rate also decreases. We have a similar case for updating $V$ and $\Delta_i$.

### 4 Properties of Orthogonal shared basis factorization

In this section, we discuss some critical mathematical properties of the factors estimated in the OSBF.

Property 1. The matrix $E$ is non-defective, the eigenvectors and eigenvalues ($\lambda_i$) are real with $\lambda_i > 0$.

*Proof.* We have $E = \mathrm{E}(R) = \frac{1}{k}\sum_{i=1}^{k} R_i$, where $R_i = X_i^T X_i / m_i$. Let $A_i = X_i^T X_i$. Now for any non zero vector $y$, we have

$$y^T A_i y = y^T X_i^T X_i y = (X_i y)^T (X_i y) = \|X_i y\|^2.$$

Columns of $X_i$ are linearly independent, then $y^T A_i y > 0$ for $y \neq 0$. So $A_i$ is a positive definite matrix. Also, the sum of $A_i$ is positive definite. Thus $E(R)$ is a non-singular matrix with positive eigenvalues. □

Property 2. The shared right basis matrix $V$ is a orthogonal matrix.

*Proof.* $E$ is a symmetric non-defective matrix. Eigenvectors of $E$ are orthogonal. □

Property 3. The species-specific eigengenes ($U$ matrices) are orthonormal.

Orthonormal eigengenes have the following advantages.

- The species-specific matrix $U_{m \times n} = \begin{bmatrix} u_1 & u_2 & \dots & u_n \end{bmatrix}$ is a non-square matrix with $m \gg n$. To project expression profiles to the common expression subspace $V$, computation of the inverse of $U_i$ matrices is required. Since the column vectors of the $U_i$ matrix are orthonormal, we can achieve this without computing a generalized inverse. Thus, a gene expression profile $Y_j$ from species $j$ can be projected to the common expression subspace using $Y_i^T U_i \Delta_i^{-1}$.

- The orthonormal property of the $U_i$ matrices summarizes the column space of $D$ using un-correlated vectors. Each $u_i$ corresponds to loadings for an independent linear combination of genes that transforms $D$ to the common expression subspace. The genes with significant contributions to the individual dimensions of the common expression subspace can be identified using coefficients (loadings) values. The un-correlated $u_i$'s help to minimize the redundancy of the biological processes represented by the common subspace and easily differentiate the contribution of genes/features across different dimensions of the common expression subspace.

#### Comparison of OSBF with other methods

Comparison of OSBF with Joint non-negative factorization (jNMF, Zhang et al., 2012) and HO GSVD (Ponnapalli et al., 2011) is summarized in Table 1. In OSBF, we achieve the following goals.

1. Biologically meaningful common expression subspace: The OSBF common expression subspace is a low-dimensional space capturing conserved inter-sample correlation relationships. In OSBF, how individual dimensions of the common subspace relate to the columns of the gene expression matrices from different species and what conserved biological processes individual dimensions represent can be effortlessly learned.

2. Interpretable factors: The independence of the columns of the species-specific factors and the common subspace helps to differentiate the contribution of genes/features across different dimensions of the common subspace. The diagonal matrices represent the amount of correlation information represented by individual dimensions of the common subspace.

3. Reducing the factorization error: Like many other joint matrix factorization approaches, OSBF is not an exact factorization. The total factorization error is minimized using an iterative algorithm.

Table 1: Comparison of joint matrix factorization approaches

| | **OSBF** | **HO GSVD** | **jNMF** |
|---|---|---|---|
| **Definition** | $\sum_{i=1}^{k} D_i \approx U_i \Delta_i V^T$ | $\sum_{i=1}^{N} D_i = U_i \Sigma_i V^T$ | $\sum_i X_i \approx W H_i$ |
| **Factorization type** | Not exact | Exact | Not exact |
| **Left basis** | Orthonormal | Not orthonormal | Not orthonormal, $W \geq 0$ |
| **Right basis** | Orthogonal | Not orthogonal | Not orthogonal, $H_i \geq 0$ |
| **Shared basis estimation** | $E = V \Lambda V^T$, where $E = \frac{1}{k} \sum_{i=1}^{k} R_i$ | $S = V \Lambda V^T$, where $S = \frac{1}{N(N-1)} \sum_{i=1}^{N} \sum_{j>i} (A_i A_j^{-1} + A_j A_i^{-1})$ | $W_{ia} = \frac{W_{ia}(\sum_i X_i H_i^T)_{ia}}{W(\sum_i (H_i H_i^T)_{ia}}$ $H_i = \frac{H_i W^T X_i}{W^T W H_i}$ |

Compared with jNMF-based approaches, the gene loading matrices in OSBF can have positive and negative loadings. In our analysis, we take advantage of the signs of genes to distinguish whether a gene is contributing to the phenotype represented by the common expression subspace or not. In jNMF, the shared factor cannot be associated with a general mathematical property related to the columns of the data matrix. In OSBF, independent of data, the common expression subspace dimensions represent directions maximizing the inter-sample correlation relationships. In HO GSVD, the steps in estimating the shared factor require complex procedures. As shown in Table 1, estimating the HO GSVD shared factor requires the computation of all pairwise quotients and their arithmetic mean. The pairwise quotient is defined as $A_i A_j^{-1}$, where $A_i = D_i^T D_i$. Compared to the inter-sample correlation relationship in OSBF, the pairwise quotients do not have any biological interpretations. Also, the estimated shared factor in HO GSVD is non-orthogonal and different dimensions of the HO GSVD shared factor do not necessarily represent properties related to conserved phenotypes. As a result, the biological interpretation of the shared factor is difficult in HO GSVD. Moreover, the independence of the columns of the species-specific factors is not guaranteed, making it challenging to differentiate the contribution of genes/features across different dimensions of the shared factor. The non-independence also requires the computation of generalized inverses to project expression profiles to the HO GSVD common subspace. These limitations restrict its application in cross-species gene expression analysis. A comparison of cross-species analysis of OSBF with HO GSVD using expression profiles of different tissue types is demonstrated in the R package vignette.

### 5 Quality checks of RNA-Seq libraries

For each project, a species-wise exploratory analysis based on principal component analysis and hierarchical clustering were performed to detect outlier libraries and fix mislabeling, if any. In each project, clustering plots: both principal component analysis (PCA) and hierarchical clustering heatmaps for all samples belonging to different species were generated separately for three approaches: variance stabilizing transformation after DESeq2 normalization (Love et al., 2014), TMM normalization, and upper-quartile normalization. TMM and upper-quartile normalization were performed using edgeR package (Robinson et al., 2010). For projects where data from multiple species are present, one-to-one orthologous protein-coding genes shared by all species were used to combine the gene expression matrix for quality checks.

Libraries from different projects belonging to the same species were analyzed to check any project-specific biases due to different experimental conditions. After appropriate normalization, samples clustered by tissue type in hierarchical clustering and PCA analysis indicated no strong biases. The analysis was repeated by performing batch correction at the project level using sva (Leek et al., 2012), and similar results were observed.

#### References


Leek JT, Johnson WE, Parker HS, Jaffe AE, Storey JD (2012) The sva package for removing batch effects and other unwanted variation in high-throughput experiments. *Bioinformatics* 28:882–883.

Love MI, Huber W, Anders S (2014) Moderated estimation of fold change and dispersion for RNA-seq data with DESeq2. *Genome Biology* 15:550.

Ponnapalli SP, Saunders MA, Van Loan CF, Alter O (2011) A higher-order generalized singular value decomposition for comparison of global mRNA expression from multiple organisms. *PLOS ONE* 6:e28072.

Robinson MD, McCarthy DJ, Smyth GK (2010) edgeR: a Bioconductor package for differential expression analysis of digital gene expression data. *Bioinformatics* 26:139–140.

Zhang S, Liu CC, Li W, Shen H, Laird PW, Zhou XJ (2012) Discovery of multi-dimensional modules by integrative analysis of cancer genomic data. *Nucleic Acids Research* 40:9379–9391.
