## Additional File 3 for "Orthogonal Shared Basis Factorization: Cross-species gene expression analysis using a common expression subspace"

### Additional File 3: Orthogonal Shared Basis Factorization

Amal Thomas

Quantitative and Computational Biology, University of Southern California, Los Angeles, CA, 90007 USA

The RNA-Seq profiles used in this study are shown in the following table.

Table 1: RNA-Seq profiles of different tissues

| **SRP ID** | **Species** | **Tissue** | **GSM ID** |
|---|---|---|---|
| SRP097223 [36] | *B.taurus* | muscle | GSM2463991 |
| SRP097223 | *B.taurus* | lung | GSM2463992 |
| SRP097223 | *B.taurus* | muscle | GSM2463993 |
| SRP097223 | *B.taurus* | lung | GSM2463994 |
| SRP097223 | *B.taurus* | muscle | GSM2463995 |
| SRP097223 | *B.taurus* | lung | GSM2463996 |
| SRP097223 | *B.taurus* | spleen | GSM2464066 |
| SRP097223 | *B.taurus* | spleen | GSM2464067 |
| SRP041696 [23] | *B.taurus* | muscle | GSM1379561 |
| SRP041696 | *B.taurus* | muscle | GSM1379562 |
| SRP041696 | *B.taurus* | muscle | GSM1379563 |
| SRP041696 | *B.taurus* | muscle | GSM1379564 |
| SRP041696 | *B.taurus* | muscle | GSM1379565 |
| SRP041696 | *B.taurus* | muscle | GSM1379566 |
| SRP052656 [33] | *B.taurus* | liver | GSM1587661 |
| SRP052656 | *B.taurus* | muscle | GSM1587663 |
| SRP052656 | *B.taurus* | liver | GSM1587665 |
| SRP052656 | *B.taurus* | muscle | GSM1587667 |
| SRP052656 | *B.taurus* | liver | GSM1587669 |
| SRP052656 | *B.taurus* | muscle | GSM1587671 |
| SRP052656 | *B.taurus* | liver | GSM1587673 |
| SRP052656 | *B.taurus* | muscle | GSM1587675 |
| SRP052656 | *B.taurus* | liver | GSM1587677 |
| SRP052656 | *B.taurus* | muscle | GSM1587679 |
| SRP052656 | *B.taurus* | liver | GSM1587681 |
| SRP052656 | *B.taurus* | muscle | GSM1587683 |
| SRP052656 | *B.taurus* | liver | GSM1587685 |
| SRP052656 | *B.taurus* | muscle | GSM1587687 |

| | | | |
|---|---|---|---|
| SRP052656 | *B.taurus* | liver | GSM1587689 |
| SRP052656 | *B.taurus* | liver | GSM1587693 |
| SRP052656 | *B.taurus* | muscle | GSM1587695 |
| SRP052656 | *B.taurus* | liver | GSM1587697 |
| SRP052656 | *B.taurus* | muscle | GSM1587699 |
| SRP056278 [17] | *B.taurus* | spleen | GSM1636044 |
| SRP056278 | *B.taurus* | spleen | GSM1636045 |
| SRP056278 | *B.taurus* | spleen | GSM1636047 |
| SRP068653 [36] | *B.taurus* | heart | GSM2042590 |
| SRP068653 | *B.taurus* | heart | GSM2042591 |
| SRP068653 | *B.taurus* | heart | GSM2042592 |
| SRP068653 | *B.taurus* | liver | GSM2042593 |
| SRP068653 | *B.taurus* | liver | GSM2042594 |
| SRP068653 | *B.taurus* | liver | GSM2042595 |
| SRP068653 | *B.taurus* | kidney | GSM2042596 |
| SRP068653 | *B.taurus* | kidney | GSM2042597 |
| SRP068653 | *B.taurus* | kidney | GSM2042598 |
| SRP083944 [40] | *B.taurus* | testis | GSM2300114 |
| SRP083944 | *B.taurus* | brain | GSM2300115 |
| SRP083944 | *B.taurus* | intestine | GSM2300119 |
| SRP012475 [24] | *B.taurus* | liver | GSM921419 |
| SRP012475 | *B.taurus* | liver | GSM921420 |
| SRP012475 | *B.taurus* | liver | GSM921421 |
| SRP012475 | *B.taurus* | liver | GSM921422 |
| SRP012475 | *B.taurus* | liver | GSM921423 |
| SRP017611 [10] | *B.taurus* | liver | GSM1054989 |
| SRP017611 | *B.taurus* | liver | GSM1054990 |
| SRP017611 | *B.taurus* | kidney | GSM1055039 |
| SRP017611 | *B.taurus* | kidney | GSM1055040 |
| SRP017611 | *B.taurus* | brain | GSM1055084 |
| SRP017611 | *B.taurus* | brain | GSM1055085 |
| SRP016501 [25] | *B.taurus* | brain | GSM1020720 |
| SRP016501 | *B.taurus* | intestine | GSM1020721 |
| SRP016501 | *B.taurus* | heart | GSM1020722 |
| SRP016501 | *B.taurus* | kidney | GSM1020723 |
| SRP016501 | *B.taurus* | liver | GSM1020724 |
| SRP016501 | *B.taurus* | lung | GSM1020725 |
| SRP016501 | *B.taurus* | muscle | GSM1020726 |
| SRP016501 | *B.taurus* | spleen | GSM1020727 |
| SRP016501 | *B.taurus* | testis | GSM1020728 |
| SRP016501 | *B.taurus* | brain | GSM1020729 |
| SRP016501 | *B.taurus* | intestine | GSM1020730 |
| SRP016501 | *B.taurus* | heart | GSM1020731 |

| | | | |
|---|---|---|---|
| SRP016501 | *B.taurus* | kidney | GSM1020732 |
| SRP016501 | *B.taurus* | liver | GSM1020733 |
| SRP016501 | *B.taurus* | lung | GSM1020734 |
| SRP016501 | *B.taurus* | muscle | GSM1020735 |
| SRP016501 | *B.taurus* | spleen | GSM1020736 |
| SRP016501 | *B.taurus* | testis | GSM1020737 |
| SRP016501 | *B.taurus* | brain | GSM1020738 |
| SRP016501 | *B.taurus* | intestine | GSM1020739 |
| SRP016501 | *B.taurus* | heart | GSM1020740 |
| SRP016501 | *B.taurus* | kidney | GSM1020741 |
| SRP016501 | *B.taurus* | liver | GSM1020742 |
| SRP016501 | *B.taurus* | lung | GSM1020743 |
| SRP016501 | *B.taurus* | spleen | GSM1020745 |
| SRP016501 | *B.taurus* | testis | GSM1020746 |
| SRP015997 [1] | *G.gallus* | brain | GSM1015156 |
| SRP015997 | *G.gallus* | liver | GSM1015157 |
| SRP015997 | *G.gallus* | kidney | GSM1015158 |
| SRP015997 | *G.gallus* | heart | GSM1015159 |
| SRP015997 | *G.gallus* | muscle | GSM1015160 |
| SRP007412 [5] | *G.gallus* | brain | GSM752557 |
| SRP007412 | *G.gallus* | brain | GSM752558 |
| SRP007412 | *G.gallus* | brain | GSM752559 |
| SRP007412 | *G.gallus* | brain | GSM752560 |
| SRP007412 | *G.gallus* | heart | GSM752561 |
| SRP007412 | *G.gallus* | heart | GSM752562 |
| SRP007412 | *G.gallus* | kidney | GSM752563 |
| SRP007412 | *G.gallus* | kidney | GSM752564 |
| SRP007412 | *G.gallus* | liver | GSM752565 |
| SRP007412 | *G.gallus* | liver | GSM752566 |
| SRP007412 | *G.gallus* | testis | GSM752567 |
| SRP007412 | *G.gallus* | testis | GSM752568 |
| SRP081121 [14] | *G.gallus* | brain | GSM2264624 |
| SRP081121 | *G.gallus* | brain | GSM2264625 |
| SRP081121 | *G.gallus* | brain | GSM2264626 |
| SRP081121 | *G.gallus* | brain | GSM2264627 |
| SRP081121 | *G.gallus* | brain | GSM2264628 |
| SRP081121 | *G.gallus* | brain | GSM2264629 |
| SRP081121 | *G.gallus* | brain | GSM2264630 |
| SRP081121 | *G.gallus* | brain | GSM2264631 |
| SRP081121 | *G.gallus* | brain | GSM2264632 |
| SRP081121 | *G.gallus* | brain | GSM2264633 |
| SRP081121 | *G.gallus* | brain | GSM2264634 |
| SRP081121 | *G.gallus* | brain | GSM2264635 |

| | | | |
|---|---|---|---|
| SRP081121 | *G.gallus* | heart | GSM2264636 |
| SRP081121 | *G.gallus* | heart | GSM2264637 |
| SRP081121 | *G.gallus* | heart | GSM2264639 |
| SRP081121 | *G.gallus* | heart | GSM2264640 |
| SRP081121 | *G.gallus* | heart | GSM2264641 |
| SRP081121 | *G.gallus* | heart | GSM2264642 |
| SRP081121 | *G.gallus* | heart | GSM2264643 |
| SRP081121 | *G.gallus* | heart | GSM2264644 |
| SRP081121 | *G.gallus* | heart | GSM2264645 |
| SRP081121 | *G.gallus* | heart | GSM2264646 |
| SRP081121 | *G.gallus* | heart | GSM2264647 |
| SRP081121 | *G.gallus* | liver | GSM2264648 |
| SRP081121 | *G.gallus* | liver | GSM2264649 |
| SRP081121 | *G.gallus* | liver | GSM2264650 |
| SRP081121 | *G.gallus* | liver | GSM2264651 |
| SRP081121 | *G.gallus* | liver | GSM2264652 |
| SRP081121 | *G.gallus* | liver | GSM2264653 |
| SRP081121 | *G.gallus* | liver | GSM2264654 |
| SRP081121 | *G.gallus* | liver | GSM2264655 |
| SRP081121 | *G.gallus* | liver | GSM2264656 |
| SRP081121 | *G.gallus* | liver | GSM2264657 |
| SRP081121 | *G.gallus* | liver | GSM2264658 |
| SRP081121 | *G.gallus* | liver | GSM2264659 |
| SRP081121 | *G.gallus* | lung | GSM2264660 |
| SRP081121 | *G.gallus* | lung | GSM2264661 |
| SRP081121 | *G.gallus* | lung | GSM2264663 |
| SRP081121 | *G.gallus* | lung | GSM2264664 |
| SRP081121 | *G.gallus* | lung | GSM2264665 |
| SRP081121 | *G.gallus* | lung | GSM2264666 |
| SRP081121 | *G.gallus* | lung | GSM2264667 |
| SRP081121 | *G.gallus* | lung | GSM2264668 |
| SRP081121 | *G.gallus* | lung | GSM2264669 |
| SRP081121 | *G.gallus* | lung | GSM2264671 |
| SRP097223 [36] | *G.gallus* | muscle | GSM2464015 |
| SRP097223 | *G.gallus* | heart | GSM2464016 |
| SRP097223 | *G.gallus* | lung | GSM2464017 |
| SRP097223 | *G.gallus* | muscle | GSM2464018 |
| SRP097223 | *G.gallus* | heart | GSM2464019 |
| SRP097223 | *G.gallus* | lung | GSM2464020 |
| SRP097223 | *G.gallus* | muscle | GSM2464021 |
| SRP097223 | *G.gallus* | heart | GSM2464022 |
| SRP097223 | *G.gallus* | muscle | GSM2464048 |
| SRP097223 | *G.gallus* | heart | GSM2464049 |

| | | | |
|---|---|---|---|
| SRP097223 | *G.gallus* | lung | GSM2464050 |
| SRP097223 | *G.gallus* | muscle | GSM2464051 |
| SRP097223 | *G.gallus* | heart | GSM2464052 |
| SRP097223 | *G.gallus* | lung | GSM2464053 |
| SRP097223 | *G.gallus* | heart | GSM2464055 |
| SRP097223 | *G.gallus* | lung | GSM2464056 |
| SRP097223 | *G.gallus* | liver | GSM2464087 |
| SRP097223 | *G.gallus* | liver | GSM2464089 |
| SRP097223 | *G.gallus* | liver | GSM2464091 |
| SRP097223 | *G.gallus* | liver | GSM2464111 |
| SRP097223 | *G.gallus* | liver | GSM2464113 |
| SRP097223 | *G.gallus* | liver | GSM2464115 |
| SRP097223 | *G.gallus* | kidney | GSM2464117 |
| SRP097223 | *G.gallus* | kidney | GSM2464118 |
| SRP097223 | *G.gallus* | kidney | GSM2464119 |
| SRP097223 | *G.gallus* | kidney | GSM2464126 |
| SRP097223 | *G.gallus* | kidney | GSM2464127 |
| SRP097223 | *G.gallus* | kidney | GSM2464128 |
| SRP159467 [28] | *G.gallus* | heart | GSM3374029 |
| SRP159467 | *G.gallus* | heart | GSM3374030 |
| SRP159467 | *G.gallus* | heart | GSM3374032 |
| SRP159467 | *G.gallus* | heart | GSM3374033 |
| SRP159467 | *G.gallus* | heart | GSM3374034 |
| SRP159467 | *G.gallus* | heart | GSM3374035 |
| SRP159467 | *G.gallus* | heart | GSM3374036 |
| SRP159467 | *G.gallus* | heart | GSM3374037 |
| SRP159467 | *G.gallus* | heart | GSM3374038 |
| SRP159467 | *G.gallus* | heart | GSM3374039 |
| SRP159467 | *G.gallus* | heart | GSM3374040 |
| SRP159467 | *G.gallus* | heart | GSM3374042 |
| SRP159467 | *G.gallus* | heart | GSM3374043 |
| SRP159467 | *G.gallus* | heart | GSM3374044 |
| SRP159467 | *G.gallus* | heart | GSM3374045 |
| SRP159467 | *G.gallus* | heart | GSM3374046 |
| SRP159467 | *G.gallus* | spleen | GSM3374065 |
| SRP159467 | *G.gallus* | spleen | GSM3374066 |
| SRP159467 | *G.gallus* | spleen | GSM3374067 |
| SRP159467 | *G.gallus* | spleen | GSM3374068 |
| SRP159467 | *G.gallus* | spleen | GSM3374069 |
| SRP159467 | *G.gallus* | spleen | GSM3374070 |
| SRP159467 | *G.gallus* | spleen | GSM3374071 |
| SRP159467 | *G.gallus* | spleen | GSM3374072 |
| SRP159467 | *G.gallus* | spleen | GSM3374073 |

| | | | |
|---|---|---|---|
| SRP159467 | *G.gallus* | spleen | GSM3374074 |
| SRP159467 | *G.gallus* | spleen | GSM3374075 |
| SRP159467 | *G.gallus* | spleen | GSM3374076 |
| SRP159467 | *G.gallus* | spleen | GSM3374077 |
| SRP159467 | *G.gallus* | spleen | GSM3374078 |
| SRP159467 | *G.gallus* | spleen | GSM3374079 |
| SRP159467 | *G.gallus* | spleen | GSM3374080 |
| SRP159467 | *G.gallus* | spleen | GSM3374081 |
| SRP159467 | *G.gallus* | spleen | GSM3374082 |
| SRP016501 [25] | *G.gallus* | brain | GSM1020747 |
| SRP016501 | *G.gallus* | intestine | GSM1020748 |
| SRP016501 | *G.gallus* | heart | GSM1020749 |
| SRP016501 | *G.gallus* | kidney | GSM1020750 |
| SRP016501 | *G.gallus* | liver | GSM1020751 |
| SRP016501 | *G.gallus* | spleen | GSM1020754 |
| SRP016501 | *G.gallus* | testis | GSM1020755 |
| SRP016501 | *G.gallus* | brain | GSM1020756 |
| SRP016501 | *G.gallus* | intestine | GSM1020757 |
| SRP016501 | *G.gallus* | heart | GSM1020758 |
| SRP016501 | *G.gallus* | kidney | GSM1020759 |
| SRP016501 | *G.gallus* | liver | GSM1020760 |
| SRP016501 | *G.gallus* | lung | GSM1020761 |
| SRP016501 | *G.gallus* | muscle | GSM1020762 |
| SRP016501 | *G.gallus* | spleen | GSM1020763 |
| SRP016501 | *G.gallus* | testis | GSM1020764 |
| SRP016501 | *G.gallus* | intestine | GSM1020766 |
| SRP016501 | *G.gallus* | heart | GSM1020767 |
| SRP016501 | *G.gallus* | kidney | GSM1020768 |
| SRP016501 | *G.gallus* | lung | GSM1020770 |
| SRP016501 | *G.gallus* | spleen | GSM1020772 |
| SRP016501 | *G.gallus* | testis | GSM1020773 |
| SRP017959 [27] | *G.gallus* | brain | GSM1064853 |
| SRP017959 | *G.gallus* | testis | GSM1064855 |
| SRP001558 [3] | *H.sapiens* | liver | GSM432598 |
| SRP001558 | *H.sapiens* | liver | GSM432599 |
| SRP001558 | *H.sapiens* | liver | GSM432600 |
| SRP001558 | *H.sapiens* | liver | GSM432601 |
| SRP001558 | *H.sapiens* | liver | GSM432602 |
| SRP001558 | *H.sapiens* | liver | GSM432603 |
| SRP001558 | *H.sapiens* | liver | GSM432604 |
| SRP001558 | *H.sapiens* | liver | GSM432605 |
| SRP001558 | *H.sapiens* | liver | GSM432606 |
| SRP001558 | *H.sapiens* | liver | GSM432607 |

| | | | |
|---|---|---|---|
| SRP001558 | *H.sapiens* | liver | GSM432608 |
| SRP001558 | *H.sapiens* | liver | GSM432609 |
| SRP007412 [5] | *H.sapiens* | brain | GSM752692 |
| SRP007412 | *H.sapiens* | brain | GSM752693 |
| SRP007412 | *H.sapiens* | brain | GSM752694 |
| SRP007412 | *H.sapiens* | brain | GSM752695 |
| SRP007412 | *H.sapiens* | brain | GSM752696 |
| SRP007412 | *H.sapiens* | brain | GSM752697 |
| SRP007412 | *H.sapiens* | brain | GSM752698 |
| SRP007412 | *H.sapiens* | heart | GSM752700 |
| SRP007412 | *H.sapiens* | heart | GSM752701 |
| SRP007412 | *H.sapiens* | kidney | GSM752702 |
| SRP007412 | *H.sapiens* | kidney | GSM752703 |
| SRP007412 | *H.sapiens* | kidney | GSM752704 |
| SRP007412 | *H.sapiens* | liver | GSM752705 |
| SRP007412 | *H.sapiens* | liver | GSM752706 |
| SRP007412 | *H.sapiens* | testis | GSM752707 |
| SRP007412 | *H.sapiens* | testis | GSM752708 |
| SRP145002 [6] | *H.sapiens* | brain | GSM3137335 |
| SRP145002 | *H.sapiens* | brain | GSM3137336 |
| SRP145002 | *H.sapiens* | brain | GSM3137337 |
| SRP145002 | *H.sapiens* | brain | GSM3137338 |
| SRP145002 | *H.sapiens* | heart | GSM3137339 |
| SRP145002 | *H.sapiens* | heart | GSM3137340 |
| SRP145002 | *H.sapiens* | heart | GSM3137341 |
| SRP145002 | *H.sapiens* | heart | GSM3137342 |
| SRP145002 | *H.sapiens* | kidney | GSM3137343 |
| SRP145002 | *H.sapiens* | kidney | GSM3137344 |
| SRP145002 | *H.sapiens* | liver | GSM3137346 |
| SRP145002 | *H.sapiens* | brain | GSM3137347 |
| SRP111096 [38] | *H.sapiens* | brain | GSM2693425 |
| SRP111096 | *H.sapiens* | brain | GSM2693433 |
| SRP111096 | *H.sapiens* | brain | GSM2693440 |
| SRP111096 | *H.sapiens* | brain | GSM2693449 |
| SRP043368 [22] | *H.sapiens* | muscle | GSM1415126 |
| SRP043368 | *H.sapiens* | muscle | GSM1415127 |
| SRP043368 | *H.sapiens* | muscle | GSM1415128 |
| SRP043368 | *H.sapiens* | muscle | GSM1415129 |
| SRP043368 | *H.sapiens* | muscle | GSM1415130 |
| SRP043368 | *H.sapiens* | muscle | GSM1415131 |
| SRP043368 | *H.sapiens* | muscle | GSM1415132 |
| SRP043368 | *H.sapiens* | muscle | GSM1415133 |
| SRP043368 | *H.sapiens* | muscle | GSM1415134 |

| | | | |
|---|---|---|---|
| SRP043368 | *H.sapiens* | muscle | GSM1415135 |
| SRP043368 | *H.sapiens* | muscle | GSM1415136 |
| SRP043368 | *H.sapiens* | muscle | GSM1415137 |
| SRP043368 | *H.sapiens* | muscle | GSM1415138 |
| SRP043368 | *H.sapiens* | muscle | GSM1415139 |
| SRP043368 | *H.sapiens* | muscle | GSM1415140 |
| SRP043368 | *H.sapiens* | muscle | GSM1415141 |
| SRP043368 | *H.sapiens* | muscle | GSM1415142 |
| SRP043368 | *H.sapiens* | muscle | GSM1415143 |
| SRP043368 | *H.sapiens* | muscle | GSM1415144 |
| SRP043368 | *H.sapiens* | muscle | GSM1415146 |
| SRP043368 | *H.sapiens* | muscle | GSM1415147 |
| SRP043368 | *H.sapiens* | muscle | GSM1415148 |
| SRP043368 | *H.sapiens* | muscle | GSM1415149 |
| SRP043108 [21] | *H.sapiens* | muscle | GSM1409687 |
| SRP043108 | *H.sapiens* | muscle | GSM1409688 |
| SRP043108 | *H.sapiens* | muscle | GSM1409689 |
| SRP043108 | *H.sapiens* | muscle | GSM1409690 |
| SRP043108 | *H.sapiens* | muscle | GSM1409691 |
| SRP043108 | *H.sapiens* | muscle | GSM1409692 |
| SRP043108 | *H.sapiens* | muscle | GSM1409693 |
| SRP043108 | *H.sapiens* | muscle | GSM1409694 |
| SRP043108 | *H.sapiens* | muscle | GSM1409695 |
| SRP043108 | *H.sapiens* | muscle | GSM1409696 |
| SRP043108 | *H.sapiens* | muscle | GSM1409697 |
| SRP043108 | *H.sapiens* | muscle | GSM1409698 |
| SRP043108 | *H.sapiens* | muscle | GSM1409699 |
| SRP043108 | *H.sapiens* | muscle | GSM1409702 |
| SRP043108 | *H.sapiens* | muscle | GSM1409703 |
| SRP043108 | *H.sapiens* | muscle | GSM1409704 |
| SRP043108 | *H.sapiens* | muscle | GSM1409705 |
| SRP043108 | *H.sapiens* | muscle | GSM1409706 |
| SRP043108 | *H.sapiens* | muscle | GSM1409707 |
| SRP043108 | *H.sapiens* | muscle | GSM1409708 |
| SRP043108 | *H.sapiens* | muscle | GSM1409709 |
| EMTAB2836 [12] | *H.sapiens* | intestine | ERS326931 |
| EMTAB2836 | *H.sapiens* | kidney | ERS326934 |
| EMTAB2836 | *H.sapiens* | liver | ERS326939 |
| EMTAB2836 | *H.sapiens* | lung | ERS326948 |
| EMTAB2836 | *H.sapiens* | lung | ERS326952 |
| EMTAB2836 | *H.sapiens* | testis | ERS326957 |
| EMTAB2836 | *H.sapiens* | liver | ERS326962 |
| EMTAB2836 | *H.sapiens* | testis | ERS326963 |

| | | | |
|---|---|---|---|
| EMTAB2836 | *H.sapiens* | spleen | ERS326970 |
| EMTAB2836 | *H.sapiens* | heart | ERS326973 |
| EMTAB2836 | *H.sapiens* | intestine | ERS326976 |
| EMTAB2836 | *H.sapiens* | intestine | ERS326978 |
| EMTAB2836 | *H.sapiens* | heart | ERS326979 |
| EMTAB2836 | *H.sapiens* | spleen | ERS326983 |
| EMTAB2836 | *H.sapiens* | testis | ERS326984 |
| EMTAB2836 | *H.sapiens* | heart | ERS326987 |
| EMTAB2836 | *H.sapiens* | liver | ERS326991 |
| EMTAB2836 | *H.sapiens* | intestine | ERS326992 |
| EMTAB2836 | *H.sapiens* | intestine | ERS326994 |
| EMTAB2836 | *H.sapiens* | testis | ERS326998 |
| EMTAB2836 | *H.sapiens* | heart | ERS327004 |
| EMTAB2836 | *H.sapiens* | spleen | ERS327006 |
| EMTAB2836 | *H.sapiens* | lung | ERS327011 |
| EMTAB2836 | *H.sapiens* | kidney | ERS327015 |
| EMTAB2836 | *H.sapiens* | testis | ERS327016 |
| EMTAB2836 | *H.sapiens* | testis | ERS327017 |
| EMTAB2836 | *H.sapiens* | lung | ERS327020 |
| EMTAB2836 | *H.sapiens* | kidney | ERS327021 |
| EMTAB2836 | *H.sapiens* | kidney | ERS327022 |
| EMTAB2836 | *H.sapiens* | testis | ERS327023 |
| EMTAB2836 | *H.sapiens* | spleen | ERS327025 |
| EMTAB513 [32] | *H.sapiens* | testis | ERR030873 |
| EMTAB513 | *H.sapiens* | brain | ERR030882 |
| EMTAB513 | *H.sapiens* | kidney | ERR030885 |
| EMTAB513 | *H.sapiens* | heart | ERR030886 |
| EMTAB513 | *H.sapiens* | liver | ERR030887 |
| EMTAB513 | *H.sapiens* | brain | ERR030890 |
| EMTAB513 | *H.sapiens* | kidney | ERR030893 |
| EMTAB513 | *H.sapiens* | heart | ERR030894 |
| EMTAB513 | *H.sapiens* | liver | ERR030895 |
| EMTAB513 | *H.sapiens* | testis | ERR030902 |
| SRP124265 [18] | *H.sapiens* | brain | GSM2842950 |
| SRP124265 | *H.sapiens* | muscle | GSM2842951 |
| SRP058740 [31] | *H.sapiens* | brain | GSM1695905 |
| SRP058740 | *H.sapiens* | brain | GSM1695906 |
| SRP058740 | *H.sapiens* | heart | GSM1695907 |
| SRP058740 | *H.sapiens* | heart | GSM1695908 |
| SRP058740 | *H.sapiens* | liver | GSM1695909 |
| SRP058740 | *H.sapiens* | liver | GSM1695910 |
| SRP058740 | *H.sapiens* | testis | GSM1695911 |
| SRP058740 | *H.sapiens* | testis | GSM1695912 |

| | | | |
|---|---|---|---|
| SRP008743 | *H.sapiens* | liver | SRX099952 |
| SRP008743 | *H.sapiens* | liver | SRX102937 |
| SRP008743 | *H.sapiens* | liver | SRX102961 |
| SRP008743 | *H.sapiens* | liver | SRX104333 |
| SRP136499 [2] | *H.sapiens* | liver | GSM3068288 |
| SRP136499 | *H.sapiens* | kidney | GSM3068289 |
| SRP136499 | *H.sapiens* | liver | GSM3068290 |
| SRP136499 | *H.sapiens* | lung | GSM3068291 |
| SRP136499 | *H.sapiens* | heart | GSM3068292 |
| SRP136499 | *H.sapiens* | kidney | GSM3068293 |
| SRP136499 | *H.sapiens* | liver | GSM3068294 |
| SRP136499 | *H.sapiens* | lung | GSM3068295 |
| SRP136499 | *H.sapiens* | heart | GSM3068296 |
| SRP136499 | *H.sapiens* | kidney | GSM3068297 |
| SRP136499 | *H.sapiens* | liver | GSM3068298 |
| SRP136499 | *H.sapiens* | lung | GSM3068299 |
| SRP136499 | *H.sapiens* | heart | GSM3068300 |
| SRP136499 | *H.sapiens* | kidney | GSM3068301 |
| SRP136499 | *H.sapiens* | liver | GSM3068302 |
| SRP136499 | *H.sapiens* | lung | GSM3068303 |
| SRP017959 [27] | *H.sapiens* | brain | GSM1064822 |
| SRP017959 | *H.sapiens* | brain | GSM1064824 |
| SRP017959 | *H.sapiens* | testis | GSM1064826 |
| SRP017959 | *H.sapiens* | brain | GSM1196040 |
| SRP017959 | *H.sapiens* | brain | GSM1196041 |
| SRP017959 | *H.sapiens* | brain | GSM1196042 |
| SRP001558 [3] | *M.mulatta* | liver | GSM432622 |
| SRP001558 | *M.mulatta* | liver | GSM432623 |
| SRP001558 | *M.mulatta* | liver | GSM432624 |
| SRP001558 | *M.mulatta* | liver | GSM432625 |
| SRP001558 | *M.mulatta* | liver | GSM432626 |
| SRP001558 | *M.mulatta* | liver | GSM432627 |
| SRP001558 | *M.mulatta* | liver | GSM432628 |
| SRP001558 | *M.mulatta* | liver | GSM432629 |
| SRP001558 | *M.mulatta* | liver | GSM432630 |
| SRP001558 | *M.mulatta* | liver | GSM432631 |
| SRP001558 | *M.mulatta* | liver | GSM432632 |
| SRP001558 | *M.mulatta* | liver | GSM432633 |
| SRP007412 [5] | *M.mulatta* | brain | GSM752631 |
| SRP007412 | *M.mulatta* | brain | GSM752632 |
| SRP007412 | *M.mulatta* | brain | GSM752633 |
| SRP007412 | *M.mulatta* | brain | GSM752634 |
| SRP007412 | *M.mulatta* | brain | GSM752635 |

| | | | |
|---|---|---|---|
| SRP007412 | *M.mulatta* | heart | GSM752636 |
| SRP007412 | *M.mulatta* | heart | GSM752637 |
| SRP007412 | *M.mulatta* | kidney | GSM752638 |
| SRP007412 | *M.mulatta* | kidney | GSM752639 |
| SRP007412 | *M.mulatta* | liver | GSM752640 |
| SRP007412 | *M.mulatta* | liver | GSM752641 |
| SRP007412 | *M.mulatta* | testis | GSM752642 |
| SRP007412 | *M.mulatta* | testis | GSM752643 |
| SRP145002 [6] | *M.mulatta* | brain | GSM3137348 |
| SRP145002 | *M.mulatta* | brain | GSM3137349 |
| SRP145002 | *M.mulatta* | brain | GSM3137350 |
| SRP145002 | *M.mulatta* | heart | GSM3137351 |
| SRP145002 | *M.mulatta* | heart | GSM3137352 |
| SRP145002 | *M.mulatta* | heart | GSM3137353 |
| SRP145002 | *M.mulatta* | kidney | GSM3137354 |
| SRP145002 | *M.mulatta* | kidney | GSM3137355 |
| SRP145002 | *M.mulatta* | kidney | GSM3137356 |
| SRP145002 | *M.mulatta* | liver | GSM3137357 |
| SRP145002 | *M.mulatta* | liver | GSM3137358 |
| SRP145002 | *M.mulatta* | liver | GSM3137359 |
| SRP145002 | *M.mulatta* | liver | GSM3137360 |
| SRP017517 [7] | *M.mulatta* | brain | GSM1051459 |
| SRP017517 | *M.mulatta* | kidney | GSM1051460 |
| SRP017517 | *M.mulatta* | lung | GSM1051461 |
| SRP019755 [19] | *M.mulatta* | brain | GSM1099737 |
| SRP019755 | *M.mulatta* | brain | GSM1099738 |
| SRP019755 | *M.mulatta* | brain | GSM1099739 |
| SRP019755 | *M.mulatta* | brain | GSM1099740 |
| SRP017517 | *M.mulatta* | heart | GSM1349923 |
| SRP017517 | *M.mulatta* | spleen | GSM1595913 |
| SRP071778 [37] | *M.mulatta* | muscle | GSM2088347 |
| SRP071778 | *M.mulatta* | muscle | GSM2088349 |
| SRP071778 | *M.mulatta* | muscle | GSM2088351 |
| SRP071778 | *M.mulatta* | muscle | GSM2088353 |
| SRP009818 [39] | *M.mulatta* | brain | GSM848932 |
| SRP009818 | *M.mulatta* | liver | GSM848933 |
| SRP009818 | *M.mulatta* | muscle | GSM848934 |
| SRP009818 | *M.mulatta* | testis | GSM848935 |
| SRP124265 | *M.mulatta* | liver | GSM2842952 |
| SRP016501 [25] | *M.mulatta* | brain | GSM1020693 |
| SRP016501 | *M.mulatta* | intestine | GSM1020694 |
| SRP016501 | *M.mulatta* | heart | GSM1020695 |
| SRP016501 | *M.mulatta* | kidney | GSM1020696 |

| | | | |
|---|---|---|---|
| SRP016501 | *M.mulatta* | liver | GSM1020697 |
| SRP016501 | *M.mulatta* | lung | GSM1020698 |
| SRP016501 | *M.mulatta* | spleen | GSM1020700 |
| SRP016501 | *M.mulatta* | testis | GSM1020701 |
| SRP016501 | *M.mulatta* | brain | GSM1020702 |
| SRP016501 | *M.mulatta* | intestine | GSM1020703 |
| SRP016501 | *M.mulatta* | heart | GSM1020704 |
| SRP016501 | *M.mulatta* | kidney | GSM1020705 |
| SRP016501 | *M.mulatta* | liver | GSM1020706 |
| SRP016501 | *M.mulatta* | lung | GSM1020707 |
| SRP016501 | *M.mulatta* | spleen | GSM1020709 |
| SRP016501 | *M.mulatta* | testis | GSM1020710 |
| SRP016501 | *M.mulatta* | brain | GSM1020711 |
| SRP016501 | *M.mulatta* | intestine | GSM1020712 |
| SRP016501 | *M.mulatta* | heart | GSM1020713 |
| SRP016501 | *M.mulatta* | kidney | GSM1020714 |
| SRP016501 | *M.mulatta* | liver | GSM1020715 |
| SRP016501 | *M.mulatta* | lung | GSM1020716 |
| SRP016501 | *M.mulatta* | spleen | GSM1020718 |
| SRP016501 | *M.mulatta* | testis | GSM1020719 |
| EMTAB6813 [26] | *M.mulatta* | brain | ERS2508654 |
| EMTAB6813 | *M.mulatta* | brain | ERS2508655 |
| EMTAB6813 | *M.mulatta* | heart | ERS2508656 |
| EMTAB6813 | *M.mulatta* | heart | ERS2508657 |
| EMTAB6813 | *M.mulatta* | brain | ERS2508658 |
| EMTAB6813 | *M.mulatta* | kidney | ERS2508659 |
| EMTAB6813 | *M.mulatta* | kidney | ERS2508660 |
| EMTAB6813 | *M.mulatta* | liver | ERS2508661 |
| EMTAB6813 | *M.mulatta* | liver | ERS2508662 |
| EMTAB6813 | *M.mulatta* | liver | ERS2508663 |
| EMTAB6813 | *M.mulatta* | testis | ERS2508664 |
| EMTAB6813 | *M.mulatta* | testis | ERS2508665 |
| EMTAB6813 | *M.mulatta* | brain | ERS2508666 |
| EMTAB6813 | *M.mulatta* | heart | ERS2508667 |
| EMTAB6813 | *M.mulatta* | heart | ERS2508668 |
| EMTAB6813 | *M.mulatta* | brain | ERS2508669 |
| EMTAB6813 | *M.mulatta* | kidney | ERS2508670 |
| EMTAB6813 | *M.mulatta* | kidney | ERS2508671 |
| EMTAB6813 | *M.mulatta* | liver | ERS2508672 |
| EMTAB6813 | *M.mulatta* | testis | ERS2508673 |
| EMTAB6813 | *M.mulatta* | brain | ERS2508674 |
| EMTAB6813 | *M.mulatta* | brain | ERS2508675 |
| EMTAB6813 | *M.mulatta* | liver | ERS2508676 |

| | | | |
|---|---|---|---|
| EMTAB6813 | *M.mulatta* | testis | ERS2508677 |
| SRP028336 [4] | *M.mulatta* | muscle | GSM1198520 |
| SRP028336 | *M.mulatta* | muscle | GSM1198521 |
| SRP028336 | *M.mulatta* | muscle | GSM1198522 |
| SRP028336 | *M.mulatta* | muscle | GSM1198523 |
| SRP028336 | *M.mulatta* | muscle | GSM1198524 |
| SRP028336 | *M.mulatta* | kidney | GSM1198525 |
| SRP028336 | *M.mulatta* | kidney | GSM1198526 |
| SRP028336 | *M.mulatta* | kidney | GSM1198527 |
| SRP028336 | *M.mulatta* | kidney | GSM1198528 |
| SRP028336 | *M.mulatta* | kidney | GSM1198529 |
| SRP028336 | *M.mulatta* | kidney | GSM1198530 |
| SRP058740 | *M.mulatta* | brain | GSM1695922 |
| SRP058740 | *M.mulatta* | heart | GSM1695923 |
| SRP058740 | *M.mulatta* | liver | GSM1695924 |
| SRP058740 | *M.mulatta* | testis | GSM1695925 |
| SRP008743 | *M.mulatta* | liver | SRX102930 |
| SRP008743 | *M.mulatta* | liver | SRX102935 |
| SRP008743 | *M.mulatta* | liver | SRX102962 |
| SRP008743 | *M.mulatta* | liver | SRX102963 |
| SRP136499 | *M.mulatta* | heart | GSM3068304 |
| SRP136499 | *M.mulatta* | kidney | GSM3068305 |
| SRP136499 | *M.mulatta* | liver | GSM3068306 |
| SRP136499 | *M.mulatta* | lung | GSM3068307 |
| SRP136499 | *M.mulatta* | heart | GSM3068308 |
| SRP136499 | *M.mulatta* | kidney | GSM3068309 |
| SRP136499 | *M.mulatta* | liver | GSM3068310 |
| SRP136499 | *M.mulatta* | lung | GSM3068311 |
| SRP136499 | *M.mulatta* | heart | GSM3068312 |
| SRP136499 | *M.mulatta* | kidney | GSM3068313 |
| SRP136499 | *M.mulatta* | liver | GSM3068314 |
| SRP136499 | *M.mulatta* | lung | GSM3068315 |
| SRP136499 | *M.mulatta* | heart | GSM3068316 |
| SRP136499 | *M.mulatta* | kidney | GSM3068317 |
| SRP136499 | *M.mulatta* | liver | GSM3068318 |
| SRP136499 | *M.mulatta* | lung | GSM3068319 |
| SRP017959 [27] | *M.mulatta* | brain | GSM1064837 |
| SRP017959 | *M.mulatta* | testis | GSM1064840 |
| EMTAB599 [13] | *M.musculus* | heart | ERX012344 |
| EMTAB599 | *M.musculus* | brain | ERX012345 |
| EMTAB599 | *M.musculus* | liver | ERX012346 |
| EMTAB599 | *M.musculus* | lung | ERX012349 |
| EMTAB599 | *M.musculus* | brain | ERX012351 |

| | | | |
|---|---|---|---|
| EMTAB599 | *M.musculus* | lung | ERX012352 |
| EMTAB599 | *M.musculus* | spleen | ERX012354 |
| EMTAB599 | *M.musculus* | spleen | ERX012356 |
| EMTAB599 | *M.musculus* | lung | ERX012357 |
| EMTAB599 | *M.musculus* | brain | ERX012358 |
| EMTAB599 | *M.musculus* | spleen | ERX012359 |
| EMTAB599 | *M.musculus* | liver | ERX012361 |
| EMTAB599 | *M.musculus* | heart | ERX012362 |
| EMTAB599 | *M.musculus* | heart | ERX012363 |
| EMTAB599 | *M.musculus* | liver | ERX012364 |
| EMTAB599 | *M.musculus* | spleen | ERX012365 |
| EMTAB599 | *M.musculus* | brain | ERX012366 |
| EMTAB599 | *M.musculus* | lung | ERX012367 |
| EMTAB599 | *M.musculus* | lung | ERX012368 |
| EMTAB599 | *M.musculus* | brain | ERX012369 |
| EMTAB599 | *M.musculus* | liver | ERX012370 |
| EMTAB599 | *M.musculus* | lung | ERX012371 |
| EMTAB599 | *M.musculus* | heart | ERX012374 |
| EMTAB599 | *M.musculus* | liver | ERX012375 |
| EMTAB599 | *M.musculus* | liver | ERX012377 |
| EMTAB599 | *M.musculus* | heart | ERX012378 |
| EMTAB599 | *M.musculus* | brain | ERX012379 |
| SRP015997 | *M.musculus* | brain | GSM1015150 |
| SRP015997 | *M.musculus* | brain | GSM1015151 |
| SRP015997 | *M.musculus* | liver | GSM1015152 |
| SRP015997 | *M.musculus* | kidney | GSM1015153 |
| SRP015997 | *M.musculus* | heart | GSM1015154 |
| SRP015997 | *M.musculus* | muscle | GSM1015155 |
| SRP007412 [5] | *M.musculus* | brain | GSM752614 |
| SRP007412 | *M.musculus* | brain | GSM752615 |
| SRP007412 | *M.musculus* | brain | GSM752616 |
| SRP007412 | *M.musculus* | brain | GSM752617 |
| SRP007412 | *M.musculus* | brain | GSM752618 |
| SRP007412 | *M.musculus* | brain | GSM752619 |
| SRP007412 | *M.musculus* | heart | GSM752620 |
| SRP007412 | *M.musculus* | heart | GSM752621 |
| SRP007412 | *M.musculus* | heart | GSM752622 |
| SRP007412 | *M.musculus* | kidney | GSM752623 |
| SRP007412 | *M.musculus* | kidney | GSM752624 |
| SRP007412 | *M.musculus* | kidney | GSM752625 |
| SRP007412 | *M.musculus* | liver | GSM752626 |
| SRP007412 | *M.musculus* | liver | GSM752627 |
| SRP007412 | *M.musculus* | liver | GSM752628 |

| | | | |
|---|---|---|---|
| SRP007412 | *M.musculus* | testis | GSM752629 |
| SRP007412 | *M.musculus* | testis | GSM752630 |
| SRP145002 [6] | *M.musculus* | brain | GSM3137373 |
| SRP145002 | *M.musculus* | brain | GSM3137374 |
| SRP145002 | *M.musculus* | heart | GSM3137375 |
| SRP145002 | *M.musculus* | heart | GSM3137376 |
| SRP145002 | *M.musculus* | kidney | GSM3137377 |
| SRP145002 | *M.musculus* | kidney | GSM3137378 |
| SRP145002 | *M.musculus* | kidney | GSM3137379 |
| SRP145002 | *M.musculus* | liver | GSM3137380 |
| SRP145002 | *M.musculus* | liver | GSM3137381 |
| SRP017611 [10] | *M.musculus* | liver | GSM1055018 |
| SRP017611 | *M.musculus* | liver | GSM1055019 |
| SRP017611 | *M.musculus* | liver | GSM1055020 |
| SRP017611 | *M.musculus* | kidney | GSM1055066 |
| SRP017611 | *M.musculus* | kidney | GSM1055067 |
| SRP017611 | *M.musculus* | kidney | GSM1055068 |
| SRP017611 | *M.musculus* | brain | GSM1055111 |
| SRP017611 | *M.musculus* | brain | GSM1055112 |
| SRP017611 | *M.musculus* | brain | GSM1055113 |
| SRP124265 | *M.musculus* | brain | GSM2842957 |
| SRP124265 | *M.musculus* | heart | GSM2842958 |
| SRP124265 | *M.musculus* | kidney | GSM2842959 |
| SRP124265 | *M.musculus* | liver | GSM2842960 |
| SRP016501 | *M.musculus* | brain | GSM1020640 |
| SRP016501 | *M.musculus* | intestine | GSM1020641 |
| SRP016501 | *M.musculus* | heart | GSM1020642 |
| SRP016501 | *M.musculus* | kidney | GSM1020643 |
| SRP016501 | *M.musculus* | liver | GSM1020644 |
| SRP016501 | *M.musculus* | lung | GSM1020645 |
| SRP016501 | *M.musculus* | muscle | GSM1020646 |
| SRP016501 | *M.musculus* | spleen | GSM1020647 |
| SRP016501 | *M.musculus* | testis | GSM1020648 |
| SRP016501 | *M.musculus* | brain | GSM1020649 |
| SRP016501 | *M.musculus* | intestine | GSM1020650 |
| SRP016501 | *M.musculus* | kidney | GSM1020651 |
| SRP016501 | *M.musculus* | liver | GSM1020652 |
| SRP016501 | *M.musculus* | lung | GSM1020653 |
| SRP016501 | *M.musculus* | muscle | GSM1020654 |
| SRP016501 | *M.musculus* | spleen | GSM1020655 |
| SRP016501 | *M.musculus* | testis | GSM1020656 |
| SRP016501 | *M.musculus* | brain | GSM1020657 |
| SRP016501 | *M.musculus* | intestine | GSM1020658 |

| | | | |
|---|---|---|---|
| SRP016501 | *M.musculus* | heart | GSM1020659 |
| SRP016501 | *M.musculus* | kidney | GSM1020660 |
| SRP016501 | *M.musculus* | liver | GSM1020661 |
| SRP016501 | *M.musculus* | lung | GSM1020662 |
| SRP016501 | *M.musculus* | muscle | GSM1020663 |
| SRP016501 | *M.musculus* | spleen | GSM1020664 |
| SRP016501 | *M.musculus* | testis | GSM1020665 |
| EMTAB6081 [34] | *M.musculus* | brain | ERR2130614 |
| EMTAB6081 | *M.musculus* | brain | ERR2130615 |
| EMTAB6081 | *M.musculus* | brain | ERR2130616 |
| EMTAB6081 | *M.musculus* | intestine | ERR2130617 |
| EMTAB6081 | *M.musculus* | intestine | ERR2130618 |
| EMTAB6081 | *M.musculus* | intestine | ERR2130619 |
| EMTAB6081 | *M.musculus* | intestine | ERR2130620 |
| EMTAB6081 | *M.musculus* | intestine | ERR2130621 |
| EMTAB6081 | *M.musculus* | intestine | ERR2130622 |
| EMTAB6081 | *M.musculus* | heart | ERR2130626 |
| EMTAB6081 | *M.musculus* | heart | ERR2130627 |
| EMTAB6081 | *M.musculus* | intestine | ERR2130628 |
| EMTAB6081 | *M.musculus* | intestine | ERR2130629 |
| EMTAB6081 | *M.musculus* | intestine | ERR2130630 |
| EMTAB6081 | *M.musculus* | intestine | ERR2130631 |
| EMTAB6081 | *M.musculus* | intestine | ERR2130632 |
| EMTAB6081 | *M.musculus* | intestine | ERR2130633 |
| EMTAB6081 | *M.musculus* | kidney | ERR2130634 |
| EMTAB6081 | *M.musculus* | kidney | ERR2130635 |
| EMTAB6081 | *M.musculus* | kidney | ERR2130636 |
| EMTAB6081 | *M.musculus* | liver | ERR2130637 |
| EMTAB6081 | *M.musculus* | liver | ERR2130638 |
| EMTAB6081 | *M.musculus* | liver | ERR2130639 |
| EMTAB6081 | *M.musculus* | muscle | ERR2130643 |
| EMTAB6081 | *M.musculus* | muscle | ERR2130644 |
| EMTAB6081 | *M.musculus* | muscle | ERR2130645 |
| SRP018127 [35] | *M.musculus* | brain | GSM1069683 |
| SRP018127 | *M.musculus* | liver | GSM1069684 |
| SRP018127 | *M.musculus* | testis | GSM1069685 |
| SRP058740 | *M.musculus* | heart | GSM1695929 |
| SRP058740 | *M.musculus* | liver | GSM1695930 |
| SRP058740 | *M.musculus* | testis | GSM1695931 |
| SRP008743 | *M.musculus* | liver | SRX104350 |
| SRP008743 | *M.musculus* | liver | SRX104354 |
| SRP008743 | *M.musculus* | liver | SRX104356 |
| SRP008743 | *M.musculus* | liver | SRX104785 |

| | | | |
|---|---|---|---|
| SRP017959 [27] | *M.musculus* | brain | GSM1064841 |
| SRP017959 | *M.musculus* | testis | GSM1064845 |
| SRP001558 [3] | *P.troglodytes* | liver | GSM432610 |
| SRP001558 | *P.troglodytes* | liver | GSM432611 |
| SRP001558 | *P.troglodytes* | liver | GSM432612 |
| SRP001558 | *P.troglodytes* | liver | GSM432613 |
| SRP001558 | *P.troglodytes* | liver | GSM432614 |
| SRP001558 | *P.troglodytes* | liver | GSM432615 |
| SRP001558 | *P.troglodytes* | liver | GSM432616 |
| SRP001558 | *P.troglodytes* | liver | GSM432617 |
| SRP001558 | *P.troglodytes* | liver | GSM432618 |
| SRP001558 | *P.troglodytes* | liver | GSM432619 |
| SRP001558 | *P.troglodytes* | liver | GSM432620 |
| SRP001558 | *P.troglodytes* | liver | GSM432621 |
| SRP007412 [5] | *P.troglodytes* | brain | GSM752664 |
| SRP007412 | *P.troglodytes* | brain | GSM752665 |
| SRP007412 | *P.troglodytes* | brain | GSM752666 |
| SRP007412 | *P.troglodytes* | brain | GSM752667 |
| SRP007412 | *P.troglodytes* | brain | GSM752668 |
| SRP007412 | *P.troglodytes* | brain | GSM752669 |
| SRP007412 | *P.troglodytes* | brain | GSM752670 |
| SRP007412 | *P.troglodytes* | brain | GSM752671 |
| SRP007412 | *P.troglodytes* | heart | GSM752672 |
| SRP007412 | *P.troglodytes* | heart | GSM752673 |
| SRP007412 | *P.troglodytes* | kidney | GSM752674 |
| SRP007412 | *P.troglodytes* | kidney | GSM752675 |
| SRP007412 | *P.troglodytes* | liver | GSM752676 |
| SRP007412 | *P.troglodytes* | liver | GSM752677 |
| SRP007412 | *P.troglodytes* | testis | GSM752678 |
| SRP028336 | *P.troglodytes* | kidney | GSM1198495 |
| SRP028336 | *P.troglodytes* | kidney | GSM1198496 |
| SRP028336 | *P.troglodytes* | kidney | GSM1198497 |
| SRP028336 | *P.troglodytes* | kidney | GSM1198498 |
| SRP028336 | *P.troglodytes* | kidney | GSM1198499 |
| SRP028336 | *P.troglodytes* | kidney | GSM1198500 |
| SRP058740 | *P.troglodytes* | brain | GSM1695914 |
| SRP058740 | *P.troglodytes* | brain | GSM1695915 |
| SRP058740 | *P.troglodytes* | heart | GSM1695916 |
| SRP058740 | *P.troglodytes* | heart | GSM1695917 |
| SRP058740 | *P.troglodytes* | liver | GSM1695918 |
| SRP058740 | *P.troglodytes* | liver | GSM1695919 |
| SRP058740 | *P.troglodytes* | testis | GSM1695920 |
| SRP058740 | *P.troglodytes* | testis | GSM1695921 |

| | | | |
|---|---|---|---|
| SRP008743 | *P.troglodytes* | liver | SRX102929 |
| SRP008743 | *P.troglodytes* | liver | SRX102938 |
| SRP008743 | *P.troglodytes* | liver | SRX102956 |
| SRP008743 | *P.troglodytes* | liver | SRX102964 |
| SRP136499 | *P.troglodytes* | heart | GSM3068272 |
| SRP136499 | *P.troglodytes* | kidney | GSM3068273 |
| SRP136499 | *P.troglodytes* | liver | GSM3068274 |
| SRP136499 | *P.troglodytes* | lung | GSM3068275 |
| SRP136499 | *P.troglodytes* | heart | GSM3068276 |
| SRP136499 | *P.troglodytes* | kidney | GSM3068277 |
| SRP136499 | *P.troglodytes* | liver | GSM3068278 |
| SRP136499 | *P.troglodytes* | lung | GSM3068279 |
| SRP136499 | *P.troglodytes* | heart | GSM3068280 |
| SRP136499 | *P.troglodytes* | kidney | GSM3068281 |
| SRP136499 | *P.troglodytes* | liver | GSM3068282 |
| SRP136499 | *P.troglodytes* | lung | GSM3068283 |
| SRP136499 | *P.troglodytes* | heart | GSM3068284 |
| SRP136499 | *P.troglodytes* | kidney | GSM3068285 |
| SRP136499 | *P.troglodytes* | liver | GSM3068286 |
| SRP136499 | *P.troglodytes* | lung | GSM3068287 |
| SRP145002 [6] | *R.norvegicus* | brain | GSM3137382 |
| SRP145002 | *R.norvegicus* | brain | GSM3137383 |
| SRP145002 | *R.norvegicus* | brain | GSM3137384 |
| SRP145002 | *R.norvegicus* | brain | GSM3137385 |
| SRP145002 | *R.norvegicus* | heart | GSM3137386 |
| SRP145002 | *R.norvegicus* | heart | GSM3137387 |
| SRP145002 | *R.norvegicus* | heart | GSM3137388 |
| SRP145002 | *R.norvegicus* | heart | GSM3137389 |
| SRP145002 | *R.norvegicus* | kidney | GSM3137390 |
| SRP145002 | *R.norvegicus* | kidney | GSM3137391 |
| SRP145002 | *R.norvegicus* | kidney | GSM3137392 |
| SRP145002 | *R.norvegicus* | kidney | GSM3137393 |
| SRP145002 | *R.norvegicus* | liver | GSM3137394 |
| SRP145002 | *R.norvegicus* | liver | GSM3137395 |
| SRP145002 | *R.norvegicus* | liver | GSM3137396 |
| SRP145002 | *R.norvegicus* | liver | GSM3137397 |
| SRP029760 [8] | *R.norvegicus* | brain | GSM1278047 |
| SRP029760 | *R.norvegicus* | brain | GSM1278048 |
| SRP029760 | *R.norvegicus* | brain | GSM1278050 |
| SRP029760 | *R.norvegicus* | heart | GSM1278051 |
| SRP029760 | *R.norvegicus* | heart | GSM1278052 |
| SRP029760 | *R.norvegicus* | kidney | GSM1278053 |
| SRP029760 | *R.norvegicus* | kidney | GSM1278054 |

| | | | |
|---|---|---|---|
| SRP029760 | *R.norvegicus* | liver | GSM1278055 |
| SRP029760 | *R.norvegicus* | liver | GSM1278056 |
| SRP029760 | *R.norvegicus* | testis | GSM1278058 |
| SRP017611 | *R.norvegicus* | liver | GSM1055025 |
| SRP017611 | *R.norvegicus* | liver | GSM1055026 |
| SRP017611 | *R.norvegicus* | liver | GSM1055027 |
| SRP017611 | *R.norvegicus* | kidney | GSM1055075 |
| SRP017611 | *R.norvegicus* | kidney | GSM1055076 |
| SRP017611 | *R.norvegicus* | kidney | GSM1055077 |
| SRP017611 | *R.norvegicus* | brain | GSM1055120 |
| SRP017611 | *R.norvegicus* | brain | GSM1055121 |
| SRP017611 | *R.norvegicus* | brain | GSM1055122 |
| SRP016501 | *R.norvegicus* | brain | GSM1020666 |
| SRP016501 | *R.norvegicus* | intestine | GSM1020667 |
| SRP016501 | *R.norvegicus* | heart | GSM1020668 |
| SRP016501 | *R.norvegicus* | kidney | GSM1020669 |
| SRP016501 | *R.norvegicus* | liver | GSM1020670 |
| SRP016501 | *R.norvegicus* | lung | GSM1020671 |
| SRP016501 | *R.norvegicus* | muscle | GSM1020672 |
| SRP016501 | *R.norvegicus* | spleen | GSM1020673 |
| SRP016501 | *R.norvegicus* | testis | GSM1020674 |
| SRP016501 | *R.norvegicus* | brain | GSM1020675 |
| SRP016501 | *R.norvegicus* | heart | GSM1020677 |
| SRP016501 | *R.norvegicus* | kidney | GSM1020678 |
| SRP016501 | *R.norvegicus* | liver | GSM1020679 |
| SRP016501 | *R.norvegicus* | lung | GSM1020680 |
| SRP016501 | *R.norvegicus* | muscle | GSM1020681 |
| SRP016501 | *R.norvegicus* | spleen | GSM1020682 |
| SRP016501 | *R.norvegicus* | testis | GSM1020683 |
| SRP016501 | *R.norvegicus* | brain | GSM1020684 |
| SRP016501 | *R.norvegicus* | intestine | GSM1020685 |
| SRP016501 | *R.norvegicus* | heart | GSM1020686 |
| SRP016501 | *R.norvegicus* | kidney | GSM1020687 |
| SRP016501 | *R.norvegicus* | lung | GSM1020689 |
| SRP016501 | *R.norvegicus* | muscle | GSM1020690 |
| SRP016501 | *R.norvegicus* | spleen | GSM1020691 |
| SRP016501 | *R.norvegicus* | testis | GSM1020692 |
| EMTAB6081 | *R.norvegicus* | brain | ERR2130652 |
| EMTAB6081 | *R.norvegicus* | brain | ERR2130653 |
| EMTAB6081 | *R.norvegicus* | brain | ERR2130654 |
| EMTAB6081 | *R.norvegicus* | intestine | ERR2130655 |
| EMTAB6081 | *R.norvegicus* | intestine | ERR2130656 |
| EMTAB6081 | *R.norvegicus* | intestine | ERR2130657 |

| | | | |
|---|---|---|---|
| EMTAB6081 | *R.norvegicus* | intestine | ERR2130658 |
| EMTAB6081 | *R.norvegicus* | intestine | ERR2130659 |
| EMTAB6081 | *R.norvegicus* | intestine | ERR2130660 |
| EMTAB6081 | *R.norvegicus* | heart | ERR2130664 |
| EMTAB6081 | *R.norvegicus* | heart | ERR2130665 |
| EMTAB6081 | *R.norvegicus* | heart | ERR2130666 |
| EMTAB6081 | *R.norvegicus* | intestine | ERR2130667 |
| EMTAB6081 | *R.norvegicus* | intestine | ERR2130668 |
| EMTAB6081 | *R.norvegicus* | intestine | ERR2130670 |
| EMTAB6081 | *R.norvegicus* | intestine | ERR2130671 |
| EMTAB6081 | *R.norvegicus* | intestine | ERR2130672 |
| EMTAB6081 | *R.norvegicus* | kidney | ERR2130673 |
| EMTAB6081 | *R.norvegicus* | kidney | ERR2130674 |
| EMTAB6081 | *R.norvegicus* | kidney | ERR2130675 |
| EMTAB6081 | *R.norvegicus* | liver | ERR2130676 |
| EMTAB6081 | *R.norvegicus* | liver | ERR2130677 |
| EMTAB6081 | *R.norvegicus* | liver | ERR2130678 |
| EMTAB6081 | *R.norvegicus* | muscle | ERR2130682 |
| EMTAB6081 | *R.norvegicus* | muscle | ERR2130683 |
| EMTAB6081 | *R.norvegicus* | muscle | ERR2130684 |
| SRP097223 | *S.scrofa* | muscle | GSM2464007 |
| SRP097223 | *S.scrofa* | heart | GSM2464008 |
| SRP097223 | *S.scrofa* | lung | GSM2464009 |
| SRP097223 | *S.scrofa* | heart | GSM2464010 |
| SRP097223 | *S.scrofa* | lung | GSM2464011 |
| SRP097223 | *S.scrofa* | muscle | GSM2464012 |
| SRP097223 | *S.scrofa* | lung | GSM2464014 |
| SRP097223 | *S.scrofa* | heart | GSM2464024 |
| SRP097223 | *S.scrofa* | lung | GSM2464025 |
| SRP097223 | *S.scrofa* | muscle | GSM2464026 |
| SRP097223 | *S.scrofa* | muscle | GSM2464029 |
| SRP097223 | *S.scrofa* | heart | GSM2464030 |
| SRP097223 | *S.scrofa* | lung | GSM2464031 |
| SRP097223 | *S.scrofa* | muscle | GSM2464032 |
| SRP097223 | *S.scrofa* | kidney | GSM2464063 |
| SRP097223 | *S.scrofa* | kidney | GSM2464064 |
| SRP097223 | *S.scrofa* | kidney | GSM2464065 |
| SRP097223 | *S.scrofa* | liver | GSM2464081 |
| SRP097223 | *S.scrofa* | spleen | GSM2464082 |
| SRP097223 | *S.scrofa* | liver | GSM2464083 |
| SRP097223 | *S.scrofa* | spleen | GSM2464084 |
| SRP097223 | *S.scrofa* | liver | GSM2464085 |
| SRP097223 | *S.scrofa* | spleen | GSM2464086 |

| | | | |
|---|---|---|---|
| SRP097223 | *S.scrofa* | liver | GSM2464093 |
| SRP097223 | *S.scrofa* | spleen | GSM2464094 |
| SRP097223 | *S.scrofa* | liver | GSM2464095 |
| SRP097223 | *S.scrofa* | spleen | GSM2464096 |
| SRP097223 | *S.scrofa* | liver | GSM2464097 |
| SRP097223 | *S.scrofa* | spleen | GSM2464098 |
| SRP097223 | *S.scrofa* | kidney | GSM2464120 |
| SRP097223 | *S.scrofa* | kidney | GSM2464121 |
| SRP097223 | *S.scrofa* | kidney | GSM2464122 |
| SRP018524 [11] | *S.scrofa* | testis | GSM1080023 |
| SRP018524 | *S.scrofa* | testis | GSM1080024 |
| SRP018524 | *S.scrofa* | testis | GSM1080025 |
| SRP018524 | *S.scrofa* | testis | GSM1080026 |
| SRP018524 | *S.scrofa* | testis | GSM1080027 |
| SRP018524 | *S.scrofa* | testis | GSM1080028 |
| SRP018524 | *S.scrofa* | testis | GSM1080029 |
| SRP018524 | *S.scrofa* | testis | GSM1080030 |
| SRP018524 | *S.scrofa* | testis | GSM1080031 |
| SRP018524 | *S.scrofa* | testis | GSM1080032 |
| SRP018524 | *S.scrofa* | liver | GSM1080033 |
| SRP018524 | *S.scrofa* | liver | GSM1080034 |
| SRP018524 | *S.scrofa* | liver | GSM1080035 |
| SRP018524 | *S.scrofa* | liver | GSM1080036 |
| SRP018524 | *S.scrofa* | liver | GSM1080037 |
| SRP018524 | *S.scrofa* | liver | GSM1080038 |
| SRP018524 | *S.scrofa* | liver | GSM1080039 |
| SRP018524 | *S.scrofa* | liver | GSM1080040 |
| SRP018524 | *S.scrofa* | liver | GSM1080041 |
| SRP018524 | *S.scrofa* | liver | GSM1080042 |
| SRP032451 [29] | *S.scrofa* | brain | GSM1256170 |
| SRP032451 | *S.scrofa* | brain | GSM1256171 |
| SRP032451 | *S.scrofa* | brain | GSM1256172 |
| SRP032451 | *S.scrofa* | brain | GSM1256173 |
| SRP032451 | *S.scrofa* | brain | GSM1256174 |
| SRP032451 | *S.scrofa* | brain | GSM1256175 |
| SRP032451 | *S.scrofa* | brain | GSM1256176 |
| SRP032451 | *S.scrofa* | brain | GSM1256177 |
| SRP032451 | *S.scrofa* | brain | GSM1256178 |
| SRP032451 | *S.scrofa* | brain | GSM1256179 |
| SRP068231 [16] | *S.scrofa* | testis | GSM2033155 |
| SRP068231 | *S.scrofa* | testis | GSM2033156 |
| SRP068231 | *S.scrofa* | testis | GSM2033157 |
| SRP068231 | *S.scrofa* | testis | GSM2033161 |

| | | | |
|---|---|---|---|
| SRP068231 | *S.scrofa* | testis | GSM2033162 |
| SRP068231 | *S.scrofa* | testis | GSM2033163 |
| SRP083002 [20] | *S.scrofa* | muscle | GSM2293304 |
| SRP083002 | *S.scrofa* | muscle | GSM2293305 |
| SRP083002 | *S.scrofa* | muscle | GSM2293306 |
| SRP083002 | *S.scrofa* | muscle | GSM2293307 |
| SRP083002 | *S.scrofa* | muscle | GSM2293308 |
| SRP083002 | *S.scrofa* | muscle | GSM2293309 |
| SRP141481 [9] | *S.scrofa* | liver | GSM2461207 |
| SRP141481 | *S.scrofa* | liver | GSM2461208 |
| SRP141481 | *S.scrofa* | liver | GSM2461209 |
| SRP141481 | *S.scrofa* | liver | GSM2461210 |
| SRP141481 | *S.scrofa* | liver | GSM2461211 |
| SRP141481 | *S.scrofa* | liver | GSM2461212 |
| SRP141481 | *S.scrofa* | liver | GSM2461213 |
| SRP141481 | *S.scrofa* | liver | GSM2461214 |
| SRP141481 | *S.scrofa* | liver | GSM2461215 |
| SRP141481 | *S.scrofa* | liver | GSM2461216 |
| SRP141481 | *S.scrofa* | liver | GSM2461217 |
| SRP141481 | *S.scrofa* | liver | GSM2461218 |
| SRP141481 | *S.scrofa* | liver | GSM2461219 |
| SRP141481 | *S.scrofa* | liver | GSM2461220 |
| SRP141481 | *S.scrofa* | liver | GSM2461221 |
| SRP141481 | *S.scrofa* | liver | GSM2461222 |
| SRP141481 | *S.scrofa* | liver | GSM2461223 |
| SRP141481 | *S.scrofa* | liver | GSM2461224 |
| SRP141481 | *S.scrofa* | liver | GSM2461225 |
| SRP141481 | *S.scrofa* | liver | GSM2461226 |
| SRP141481 | *S.scrofa* | liver | GSM2461227 |
| SRP141481 | *S.scrofa* | liver | GSM2461228 |
| SRP141481 | *S.scrofa* | liver | GSM2461229 |
| SRP141481 | *S.scrofa* | liver | GSM2461230 |
| SRP141481 | *S.scrofa* | liver | GSM2461231 |
| SRP141481 | *S.scrofa* | liver | GSM2461232 |
| SRP141481 | *S.scrofa* | liver | GSM2461233 |
| SRP141481 | *S.scrofa* | liver | GSM2461234 |
| SRP141481 | *S.scrofa* | liver | GSM2461235 |
| SRP141481 | *S.scrofa* | liver | GSM2461236 |
| SRP141481 | *S.scrofa* | liver | GSM2461237 |
| SRP141481 | *S.scrofa* | liver | GSM2461238 |
| SRP141481 | *S.scrofa* | liver | GSM2461239 |
| SRP141481 | *S.scrofa* | liver | GSM2461240 |
| SRP141481 | *S.scrofa* | liver | GSM2461241 |

| | | | |
|---|---|---|---|
| SRP141481 | *S.scrofa* | liver | GSM2461242 |
| SRP141481 | *S.scrofa* | liver | GSM2461243 |
| SRP141481 | *S.scrofa* | liver | GSM2461244 |
| SRP141481 | *S.scrofa* | liver | GSM2461245 |
| SRP141481 | *S.scrofa* | liver | GSM2461246 |
| SRP141481 | *S.scrofa* | liver | GSM2461247 |
| SRP141481 | *S.scrofa* | liver | GSM2461248 |
| SRP141481 | *S.scrofa* | liver | GSM2461249 |
| SRP141481 | *S.scrofa* | liver | GSM2461250 |
| SRP141481 | *S.scrofa* | liver | GSM2461251 |
| SRP141481 | *S.scrofa* | liver | GSM2461252 |
| SRP141481 | *S.scrofa* | liver | GSM2461253 |
| SRP141481 | *S.scrofa* | liver | GSM2461254 |
| SRP141481 | *S.scrofa* | testis | GSM2461255 |
| SRP141481 | *S.scrofa* | testis | GSM2461256 |
| SRP141481 | *S.scrofa* | testis | GSM2461257 |
| SRP141481 | *S.scrofa* | testis | GSM2461258 |
| SRP141481 | *S.scrofa* | testis | GSM2461259 |
| SRP141481 | *S.scrofa* | testis | GSM2461260 |
| SRP141481 | *S.scrofa* | testis | GSM2461261 |
| SRP141481 | *S.scrofa* | testis | GSM2461262 |
| SRP141481 | *S.scrofa* | testis | GSM2461263 |
| SRP141481 | *S.scrofa* | testis | GSM2461264 |
| SRP141481 | *S.scrofa* | testis | GSM2461265 |
| SRP141481 | *S.scrofa* | testis | GSM2461266 |
| SRP141481 | *S.scrofa* | testis | GSM2461267 |
| SRP141481 | *S.scrofa* | testis | GSM2461268 |
| SRP141481 | *S.scrofa* | testis | GSM2461269 |
| SRP141481 | *S.scrofa* | testis | GSM2461270 |
| SRP141481 | *S.scrofa* | testis | GSM2461271 |
| SRP141481 | *S.scrofa* | testis | GSM2461272 |
| SRP141481 | *S.scrofa* | testis | GSM2461273 |
| SRP141481 | *S.scrofa* | testis | GSM2461274 |
| SRP141481 | *S.scrofa* | testis | GSM2461275 |
| SRP141481 | *S.scrofa* | testis | GSM2461277 |
| SRP141481 | *S.scrofa* | testis | GSM2461278 |
| SRP141481 | *S.scrofa* | testis | GSM2461279 |
| SRP141481 | *S.scrofa* | testis | GSM2461280 |
| SRP141481 | *S.scrofa* | testis | GSM2461281 |
| SRP141481 | *S.scrofa* | testis | GSM2461282 |
| SRP141481 | *S.scrofa* | testis | GSM2461283 |
| SRP141481 | *S.scrofa* | testis | GSM2461284 |
| SRP141481 | *S.scrofa* | testis | GSM2461285 |

| | | | |
|---|---|---|---|
| SRP141481 | *S.scrofa* | testis | GSM2461286 |
| SRP141481 | *S.scrofa* | testis | GSM2461287 |
| SRP141481 | *S.scrofa* | testis | GSM2461288 |
| SRP141481 | *S.scrofa* | testis | GSM2461289 |
| SRP141481 | *S.scrofa* | testis | GSM2461290 |
| SRP141481 | *S.scrofa* | testis | GSM2461291 |
| SRP141481 | *S.scrofa* | testis | GSM2461292 |
| SRP141481 | *S.scrofa* | testis | GSM2461293 |
| SRP141481 | *S.scrofa* | testis | GSM2461294 |
| SRP141481 | *S.scrofa* | testis | GSM2461295 |
| SRP141481 | *S.scrofa* | testis | GSM2461296 |
| SRP141481 | *S.scrofa* | testis | GSM2461297 |
| SRP141481 | *S.scrofa* | testis | GSM2461298 |
| SRP141481 | *S.scrofa* | testis | GSM2461299 |
| SRP141481 | *S.scrofa* | testis | GSM2461300 |
| SRP141481 | *S.scrofa* | testis | GSM2461301 |
| SRP141481 | *S.scrofa* | testis | GSM2461302 |
| SRP125390 [30] | *S.scrofa* | muscle | GSM2862765 |
| SRP125390 | *S.scrofa* | muscle | GSM2862766 |
| SRP125390 | *S.scrofa* | muscle | GSM2862767 |
| SRP125390 | *S.scrofa* | muscle | GSM2862768 |
| SRP125390 | *S.scrofa* | muscle | GSM2862769 |
| SRP125390 | *S.scrofa* | muscle | GSM2862770 |
| SRP125390 | *S.scrofa* | muscle | GSM2862771 |
| SRP125390 | *S.scrofa* | muscle | GSM2862772 |
| SRP125390 | *S.scrofa* | muscle | GSM2862773 |
| SRP125390 | *S.scrofa* | muscle | GSM2862774 |
| SRP125390 | *S.scrofa* | muscle | GSM2862775 |
| SRP125390 | *S.scrofa* | muscle | GSM2862776 |
| SRP125390 | *S.scrofa* | muscle | GSM2862777 |
| SRP017611 | *S.scrofa* | liver | GSM1055031 |
| SRP017611 | *S.scrofa* | liver | GSM1055032 |
| SRP017611 | *S.scrofa* | kidney | GSM1055078 |
| SRP017611 | *S.scrofa* | kidney | GSM1055079 |
| SRP017611 | *S.scrofa* | brain | GSM1055123 |
| SRP017611 | *S.scrofa* | brain | GSM1055124 |
| SRP124265 | *S.scrofa* | brain | GSM2842962 |
| SRP124265 | *S.scrofa* | heart | GSM2842963 |
| SRP124265 | *S.scrofa* | kidney | GSM2842964 |
| SRP124265 | *S.scrofa* | liver | GSM2842965 |
| SRP124265 | *S.scrofa* | muscle | GSM2842966 |
| SRP018288 [15] | *S.scrofa* | heart | GSM1074091 |
| SRP018288 | *S.scrofa* | liver | GSM1074092 |

| | | | |
|---|---|---|---|
| SRP018288 | *S.scrofa* | kidney | GSM1074094 |

### References


[1] Nuno L Barbosa-Morais, Manuel Irimia, Qun Pan, Hui Y Xiong, Serge Gueroussov, Leo J Lee, Valentina Slobodeniuc, Claudia Kutter, Stephen Watt, Recep Colak, et al. The evolutionary landscape of alternative splicing in vertebrate species. *Science*, 338(6114):1587–1593, 2012.

[2] Lauren E Blake, Julien Roux, Irene Hernando-Herraez, Nicholas E Banovich, Raquel Garcia Perez, Chiaowen Joyce Hsiao, Ittai Eres, Claudia Cuevas, Tomas Marques-Bonet, and Yoav Gilad. A comparison of gene expression and DNA methylation patterns across tissues and species. *Genome Research*, 30(2):250–262, 2020.

[3] Ran Blekhman, John C Marioni, Paul Zumbo, Matthew Stephens, and Yoav Gilad. Sex-specific and lineage-specific alternative splicing in primates. *Genome Research*, 20(2):180–189, 2010.

[4] Katarzyna Bozek, Yuning Wei, Zheng Yan, Xiling Liu, Jieyi Xiong, Masahiro Sugimoto, Masaru Tomita, Svante Pääbo, Raik Pieszek, Chet C Sherwood, et al. Exceptional evolutionary divergence of human muscle and brain metabolomes parallels human cognitive and physical uniqueness. *PLoS Biology*, 12(5):e1001871, 2014.

[5] David Brawand, Magali Soumillon, Anamaria Necsulea, Philippe Julien, Gábor Csárdi, Patrick Harrigan, Manuela Weier, Angélica Liechti, Ayinuer Aximu-Petri, Martin Kircher, et al. The evolution of gene expression levels in mammalian organs. *Nature*, 478(7369):343–348, 2011.

[6] Francesco N Carelli, Angélica Liechti, Jean Halbert, Maria Warnefors, and Henrik Kaessmann. Repurposing of promoters and enhancers during mammalian evolution. *Nature Communications*, 9(1): 1–11, 2018.

[7] Jia-Yu Chen, Zhiyu Peng, Rongli Zhang, Xin-Zhuang Yang, Bertrand Chin-Ming Tan, Huaying Fang, Chu-Jun Liu, Mingming Shi, Zhi-Qiang Ye, Yong E Zhang, et al. RNA editome in rhesus macaque shaped by purifying selection. *PLoS Genetics*, 10(4):e1004274, 2014.

[8] Diego Cortez, Ray Marin, Deborah Toledo-Flores, Laure Froidevaux, Angelica Liechti, Paul D Waters, Frank Grützner, and Henrik Kaessmann. Origins and functional evolution of y chromosomes across mammals. *Nature*, 508(7497):488–493, 2014.

[9] Markus Drag, Ruta Skinkyté-Juskiené, Duy N Do, Lisette JA Kogelman, and Haja N Kadarmideen. Differential expression and co-expression gene networks reveal candidate biomarkers of boar taint in non-castrated pigs. *Scientific Reports*, 7(1):1–18, 2017.

[10] Alexey A Fushan, Anton A Turanov, Sang-Goo Lee, Eun Bae Kim, Alexei V Lobanov, Sun Hee Yim, Rochelle Buffenstein, Sang-Rae Lee, Kyu-Tae Chang, Hwanseok Rhee, et al. Gene expression defines natural changes in mammalian lifespan. *Aging Cell*, 14(3):352–365, 2015.

[11] Asep Gunawan, Sudeep Sahadevan, Christiane Neuhoff, Christine Große-Brinkhaus, Ahmed Gad, Luc Frieden, Dawit Tesfaye, Ernst Tholen, Christian Looft, Muhammad Jasim Uddin, et al. RNA deep

sequencing reveals novel candidate genes and polymorphisms in boar testis and liver tissues with divergent androstenone levels. *PloS One*, 8(5):e63259, 2013.

[12] Björn Hallström. RNA-seq of coding RNA from tissue samples of 122 human individuals representing 32 different tissues. `https://www.ebi.ac.uk/biostudies/arrayexpress/studies/E-MTAB-2836`, 2015.

[13] Thomas M Keane, Leo Goodstadt, Petr Danecek, Michael A White, Kim Wong, Binnaz Yalcin, Andreas Heger, Avigail Agam, Guy Slater, Martin Goodson, et al. Mouse genomic variation and its effect on phenotypes and gene regulation. *Nature*, 477(7364):289–294, 2011.

[14] Diyan Li, Yan Li, Miao Li, Tiandong Che, Shilin Tian, Binlong Chen, Xuming Zhou, Guolong Zhang, Uma Gaur, Majing Luo, et al. Population genomics identifies patterns of genetic diversity and selection in chicken. *BMC Genomics*, 20(1):1–12, 2019.

[15] Mingzhou Li, Shilin Tian, Long Jin, Guangyu Zhou, Ying Li, Yuan Zhang, Tao Wang, Carol KL Yeung, Lei Chen, Jideng Ma, et al. Genomic analyses identify distinct patterns of selection in domesticated pigs and Tibetan wild boars. *Nature Genetics*, 45(12):1431–1438, 2013.

[16] Yao Li, Jialian Li, Chengchi Fang, Liang Shi, Jiajian Tan, Yuanzhu Xiong, Bin Fan, and Changchun Li. Genome-wide differential expression of genes and small rnas in testis of two different porcine breeds and at two different ages. *Scientific Reports*, 6(1):1–16, 2016.

[17] Yaokun Li, José A Carrillo, Yi Ding, Yanghua He, Chunping Zhao, Jianan Liu, George E Liu, Linsen Zan, and Jiuzhou Song. Transcriptomic profiling of spleen in grass-fed and grain-fed angus cattle. *PLoS One*, 10(9):e0135670, 2015.

[18] Yumei Li, Chen Li, Shuxian Li, Qi Peng, Ni A An, Aibin He, and Chuan-Yun Li. Human exonization through differential nucleosome occupancy. *Proceedings of the National Academy of Sciences*, 115 (35):8817–8822, 2018.

[19] Zhongshan Li, Hindrike Bammann, Mingshuang Li, Hongyu Liang, Zheng Yan, Yi-Ping Phoebe Chen, Min Zhao, and Philipp Khaitovich. Evolutionary and ontogenetic changes in RNA editing in human, chimpanzee, and macaque brains. *RNA*, 19(12):1693–1702, 2013.

[20] KS Lim, KT Lee, JE Park, WH Chung, GW Jang, BH Choi, Ki Chang Hong, and TH Kim. Identification of differentially expressed genes in longissimus muscle of pigs with high and low intramuscular fat content using rna sequencing. *Animal Genetics*, 48(2):166–174, 2017.

[21] Malene E Lindholm, Mikael Huss, Beata W Solnestam, Sanela Kjellqvist, Joakim Lundeberg, and Carl J Sundberg. The human skeletal muscle transcriptome: sex differences, alternative splicing, and tissue homogeneity assessed with RNA sequencing. *The FASEB Journal*, 28(10):4571–4581, 2014.

[22] Maléne E Lindholm, Francesco Marabita, David Gomez-Cabrero, Helene Rundqvist, Tomas J Ekström, Jesper Tegnér, and Carl Johan Sundberg. An integrative analysis reveals coordinated reprogramming of the epigenome and the transcriptome in human skeletal muscle after training. *Epigenetics*, 9 (12):1557–1569, 2014.

[23] GF Liu, HJ Cheng, W You, EL Song, XM Liu, and FC Wan. Transcriptome profiling of muscle by rna-seq reveals significant differences in digital gene expression profiling between angus and luxi cattle. *Animal Production Science*, 55(9):1172–1178, 2015.

[24] Matthew McCabe, Sinéad Waters, Dermot Morris, David Kenny, David Lynn, and Chris Creevey. Rna-seq analysis of differential gene expression in liver from lactating dairy cows divergent in negative energy balance. *BMC Genomics*, 13(1):1–11, 2012.

[25] Jason Merkin, Caitlin Russell, Ping Chen, and Christopher B Burge. Evolutionary dynamics of gene and isoform regulation in mammalian tissues. *Science*, 338(6114):1593–1599, 2012.

[26] Margarida Cardoso Moreira. Rhesus macaque RNA-seq time-series of the development of six major organs. `https://www.ebi.ac.uk/biostudies/arrayexpress/studies/E-MTAB-6813`, 2019.

[27] Anamaria Necsulea, Magali Soumillon, Maria Warnefors, Angélica Liechti, Tasman Daish, Ulrich Zeller, Julie C Baker, Frank Grützner, and Henrik Kaessmann. The evolution of lncrna repertoires and expression patterns in tetrapods. *Nature*, 505(7485):635–640, 2014.

[28] W Park, K Srikanth, D Lim, M Park, T Hur, S Kemp, Tadelle Dessie, MS Kim, S-R Lee, MFW Te Pas, et al. Comparative transcriptome analysis of ethiopian indigenous chickens from low and high altitudes under heat stress condition reveals differential immune response. *Animal Genetics*, 50(1):42–53, 2019.

[29] Dafne Pérez-Montarelo, Ole Madsen, Estefânia Alves, M Carmen Rodríguez, Josep María Folch, José Luis Noguera, Martien AM Groenen, and Ana I Fernández. Identification of genes regulating growth and fatness traits in pig through hypothalamic transcriptome analysis. *Physiological Genomics*, 46(6):195–206, 2014.

[30] Katarzyna Ropka-Molik, Klaudia Pawlina-Tyszko, Kacper Żukowski, Katarzyna Piórkowska, Grzegorz Żak, Artur Gurgul, Natalia Derebecka, and Joanna Wesoły. Examining the genetic background of porcine muscle growth and development based on transcriptome and mirnaome data. *International journal of molecular sciences*, 19(4):1208, 2018.

[31] Jorge Ruiz-Orera, Jessica Hernandez-Rodriguez, Cristina Chiva, Eduard Sabidó, Ivanela Kondova, Ronald Bontrop, Tomàs Marqués-Bonet, and M Mar Albà. Origins of de novo genes in human and chimpanzee. *PLoS Genetics*, 11(12):e1005721, 2015.

[32] Gary Schroth. RNA-Seq of human individual tissues and mixture of 16 tissues (Illumina Body Map). `https://www.ebi.ac.uk/biostudies/arrayexpress/studies/E-MTAB-513`, 2011.

[33] Minseok Seo, Kelsey Caetano-Anolles, Sandra Rodriguez-Zas, Sojeong Ka, Jin Young Jeong, Sungkwon Park, Min Ji Kim, Whan-Gook Nho, Seoae Cho, Heebal Kim, et al. Comprehensive identification of sexually dimorphic genes in diverse cattle tissues using RNA-seq. *BMC genomics*, 17(1):1–18, 2016.

[34] Julia F Söllner, German Leparc, Tobias Hildebrandt, Holger Klein, Leo Thomas, Elia Stupka, and Eric Simon. An RNA-Seq atlas of gene expression in mouse and rat normal tissues. *Scientific data*, 4(1): 1–11, 2017.

[35] Magali Soumillon, Anamaria Necsulea, Manuela Weier, David Brawand, Xiaolan Zhang, Hongcang Gu, Pauline Barthes, Maria Kokkinaki, Serge Nef, Andreas Gnirke, et al. Cellular source and mechanisms of high transcriptome complexity in the mammalian testis. *Cell Reports*, 3(6):2179–2190, 2013.

[36] Qianzi Tang, Yiren Gu, Xuming Zhou, Long Jin, Jiuqiang Guan, Rui Liu, Jing Li, Kereng Long, Shilin Tian, Tiandong Che, et al. Comparative transcriptomics of 5 high-altitude vertebrates and their low-altitude relatives. *Gigascience*, 6(12):gix105, 2017.

[37] Yao Xiao, Can Wang, Jia-Yu Chen, Fujian Lu, Jue Wang, Ning Hou, Xiaomin Hu, Fanxin Zeng, Dongwei Ma, Xueting Sun, et al. Deficiency of prkd2 triggers hyperinsulinemia and metabolic disorders. *Nature Communications*, 9(1):1–11, 2018.

[38] Chuan Xu, Qian Li, Olga Efimova, Liu He, Shoji Tatsumoto, Vita Stepanova, Takao Oishi, Toshifumi Udono, Katsushi Yamaguchi, Shuji Shigenobu, et al. Human-specific features of spatial gene expression and regulation in eight brain regions. *Genome Research*, 28(8):1097–1110, 2018.

[39] Shi-Jian Zhang, Chu-Jun Liu, Mingming Shi, Lei Kong, Jia-Yu Chen, Wei-Zhen Zhou, Xiaotong Zhu, Peng Yu, Jue Wang, Xinzhuang Yang, et al. RhesusBase: a knowledgebase for the monkey research community. *Nucleic Acids Research*, 41(D1):D892–D905, 2013.

[40] Yang Zhou, Lingyang Xu, Derek M Bickhart, Steven G Schroeder, Erin E Connor, Leeson J Alexander, Tad S Sonstegard, Curtis P Van Tassell, Hong Chen, George E Liu, et al. Reduced representation bisulphite sequencing of ten bovine somatic tissues reveals dna methylation patterns and their impacts on gene expression. *BMC Genomics*, 17(1):1–11, 2016.
